# Resource distribution unifies optimal offspring size and bacterial aging

**DOI:** 10.1101/2023.06.28.546398

**Authors:** Taom Sakal, Stephen Proulx

## Abstract

Models of optimal offspring size and bacterial aging share the same underlying mathematical problem: how should a parent optimally distribute limited resources among its offspring? Optimal offspring size theory has long explored the trade-off between offspring number and size in higher organisms. Meanwhile, the emerging field of bacterial aging examines whether and under what conditions cells evolve unequal sharing of old cellular components. Despite addressing similar problems, these models remain constrained by field-specific assumptions. We unify them in a generalized resource-distribution framework that yields insights and predictions unreachable by either field alone. Our central finding is that the convexity of the function relating resources to offspring survivorship determines the optimal resource distribution strategy. Furthermore, we show that these strategies evolve, characterize their robustness to fluctuating environments, and uncover the conditions that select for producing a “runt of the litter.”

## 1. Introduction

A newly hatched chick requires food from its parent, else it is sure to die. It cries out in hunger, as do its nestmates, and the parent only has so many worms to go around. Feed it too little, and the chick will grow weak and die. Feed it too much, and little remains for the other chicks, who grow weak and die instead. The parent faces a central question: how should it distribute the food to maximize its number of surviving offspring? This central question underlies optimal offspring size (OOS) theory.

A parallel story unfolds in the microscopic world. When a bacterium replicates, it must duplicate its components and distribute them between its two daughter cells. Newly created components perform better than older ones, so the daughter inheriting more new components will grow faster and have a better chance of survival. However, if a daughter receives too many components of too old an age then it will die, as if by senescence. The mother could distribute the old and new components equally between the daughters. Or she could favor one daughter with all new components and offload the old components onto the other. Or she could choose any of the infinite strategies between these extremes. A good choice is essential; like the mother bird, our bacterium seeks a distribution strategy that maximizes the number of surviving offspring. The study of this, and the conditions under which asymmetric division of components leads to senescence, is the domain of bacterial aging (BA) theory.

Optimal offspring size (OOS) theory and bacterial aging (BA) theory emerged separately and share little history, yet both address the same fundamental question: How should parents optimally distribute limited resources among offspring? This naturally raises another question: will these optimal strategies actually evolve?

To answer these questions, we build an optimization model that integrates macroscopic (offspring size) and microscopic (bacterial aging) perspectives within a generalized convexity framework. We show that the convexity (second derivative) of *survivorship*—the relationship between resources and offspring survival—determines whether symmetric or asymmetric resource distributions evolve. Specifically, concave survivorship favors symmetric distributions, while convex survivorship favors asymmetric distributions. These strategies consistently evolve and are summarized in Table 2 and Table 3.

**Table 1.** Terms and symbols for the constant environment model.

| Term | Definition |
| --- | --- |
| Resource | Something that is distributed among offspring that affects the offspring survival probability. Despite the positive connotation of the word, resources do not always improve offspring survival probability. |
| Fitness | A parent's expected number of surviving offspring. |
| Survivorship | The probability that an offspring will survive to reproductive age. |
| Strategy | How a parent distributes resources among its offspring. Represented by the vector $\vec{r}$. |
| Offspring on the curved region | An offspring that receives $r_{low} \leq r \leq r_{high}$ resources. |
| Symmetric Strategy | A distribution strategy that equally distributes resources among offspring in the curved region. |
| Asymmetric Strategy | A distribution strategy where all offspring on the curved region have either $r_{low}$ or $r_{high}$ resources, with at most one offspring receiving an amount $r_r$ between these two. |
| $r_i$ | The resources the $i$th offspring receives. Also called the offspring's size. |
| $\vec{r}$ | The resource distribution strategy. This is a vector of resource amounts $(r_1, r_2, \dots, r_n)$ where the $i$th offspring receives $r_i$ resources (or, depending on context, $r_i$ resources above $r_{low}$.) |
| $s(r)$ | The survivorship function. Outputs the survival probability of an offspring given $r$ resources. |
| $s_c(r)$ | The curved region of $s$. This is the function $s$ restricted to offspring such that $r_{low} \leq r \leq r_{high}$. |
| $R$ | The total amount of resources the parent has available to distribute. |
| $r_{low}$ | The minimum threshold, after which resources begin affecting survivorship. The lower border of the curved region. Must be greater than or equal to zero. |
| $r_{high}$ | The maximum threshold, after which resources stop affecting survivorship. The upper border of the curved region. Must be greater than $r_{low}$ but can be infinite. |
| $r_{max}$ | The maximum amount of resource an offspring can receive. |
| $\gamma_{low}$ | The survival probability of offspring with $r_{low}$ or less resources. |
| $\gamma_{high}$ | The survival probability of offspring with $r_{high}$ or more resources. |
| $n$ | The length of $\vec{r}$. Can be interpreted either as the number of offspring or the *maximum* number of offspring. In the latter case, we interpret offspring with size $r_{low}$ or less as offspring that are not born. |
| $w(\vec{r})$ | The fitness of distribution strategy $\vec{r}$. The expected number of surviving offspring in a constant environment. |

**Table 2.** Forms of optimal strategies when the parent is not required to distribute all resources. In the example strategies, the number of offspring is *n* = 6.

| $s_c$ | Optimal Strategy<br>(may withhold resources) | Example<br>strategy vector |
| --- | --- | --- |
| Concave-increasing | Calculate the optimal offspring number as in the Smith-Fretwell model. Distribute all resources equally among these offspring. Give all remaining offspring zero resources. | $(r^*, r^*, r^*, r^*, 0, 0)$ |
| Concave-decreasing | Give each offspring no more than $r_{low}$ resources. | $(0, 0, 0, 0, 0, 0)$ |
| Concave-increasing-decreasing | Same strategy as concave-increasing, except never give an offspring enough resources to enter the decreasing part of the curve. | $(r^*, r^*, r^*, r^*, 0, 0)$ |
| Convex-increasing | Create as many offspring with $r_{max}$ resources as possible. Put all remaining resources into a single offspring. | $(r_{max}, r_{max}, r_{max}, r_r, 0, 0)$ |
| Convex-decreasing | Give each offspring no more than $r_{low}$ resources. | $(0, 0, 0, 0, 0, 0)$ |
| Convex-decreasing-increasing | If $\gamma_{low} \geq \gamma_{high}$ then give each offspring no more than $r_{low}$ resources. Otherwise the optimal strategy depends on the details of $s$ and $R$. | *Multiple Forms* |

**Table 3.** Forms of optimal strategies when the parent is required to distribute all resources. For the example forms, the number of offspring is *n* = 6.

| $s_c$ | Optimal Strategy<br>(must distribute all resources) | Example<br>strategy vector |
| --- | --- | --- |
| Concave-increasing | Same as when not required to distribute all resources. | $(r^*, r^*, r^*, r^*, 0, 0)$ |
| Concave-decreasing | Find the optimal anti-resource offspring size $\bar{r}$. Make as many offspring of size $r_{max} - \bar{r}$ as possible. All other offspring will be size $r_{max}$. | $(r_{max}, r_{max}, r_{max}, r_{max}, r_{max} - \bar{r}, r_{max} - \bar{r})$ |
| Concave-increasing-decreasing | Same as the concave decreasing case but with a different way of calculating the optimal anti-resource offspring size ($\bar{r}'$). | $(r_{max}, r_{max}, r_{max}, r_{max}, r_{max} - \bar{r}', r_{max} - \bar{r}')$ |
| Convex-increasing | Same as when not required to distribute all resources. | $(r_{max}, r_{max}, r_{max}, r_r, 0, 0)$ |
| Convex-decreasing | Create as many offspring with $r_{low}$ resources as possible. For any remaining resources, sequentially take each offspring and fill them up to $r_{max}$, or as high as possible. Place all remaining resources in a $r_r$ sized offspring. | $(r_{max}, r_{max}, r_r, r_{low}, r_{low}, r_{low})$ |
| Convex-decreasing-increasing | Same as either the concave-increasing or concave-decreasing case if in the optimal strategy there exists an offspring with size between $r_{low}$ and $r_{high}$. Otherwise, the strategy depends on specific parameter values. | *Multiple Forms* |

This model (1) agrees with and strengthens existing models by making the same predictions in more general settings, (2) organizes existing models into a unifying concavity-based framework, (3) makes new predictions, including when runts evolve, and (4) reveals universal properties of resource distribution models. By virtue of being an optimization model, our results extend beyond the original scope and apply to any system with an analogous mathematical structure. Optimal Foraging Theory is one such example (Charnov, 1976), but the applications need not be biological.

In Section 2, we review the history of OOS and BA. Section 3 sets up the generalized resource distribution model that applies to both OOS and BA, then presents the core mathematical results. Section 4 extends these results to fluctuating environments.

## 2. Background

### A. Optimal Offspring Size (OOS) Theory

A parent can either make many offspring with few resources each, or few offspring with many resources each. OOS theory studies this trade-off, and likely began with Lack’s observation of bird clutch sizes (Lack, 1954).

The Smith-Fretwell model forms the mathematical cornerstone of this theory, predicting that (1) an optimal offspring size exists, (2) the parent should make as many offspring of the optimal size as possible, and (3) the optimal offspring size is independent of the parent’s total available resources (Smith and Fretwell, 1974). In the years since, biologists have refined this model to account for allometrics (Charnov and Ernest, 2006), multiple types of resources (McGinley and Charnov, 1988), stage-based effects of resources (Mangel et al., 1994; Hendry et al., 2001), and variable environments (Forbes, 1991; Olofsson et al., 2009; Monro et al., 2010) to name but a few. See Roff, 1993 for a rich review of empirical and theoretical studies, including many applications and variations of the Smith-Fretwell model. Marshall et al., 2018 offers a more modern review.

In these models, resources improve offspring survivorship but have diminishing returns, resulting in a survivorship function of the form shown in Figure 1. There are also situations where we expect survivorship to have a more sigmoidal shape (Dani and Kodandaramaiah, 2017), or to decrease once an offspring is given too many resources (Hendry et al., 2001; Marshall et al., 2008).

**Figure 1.**
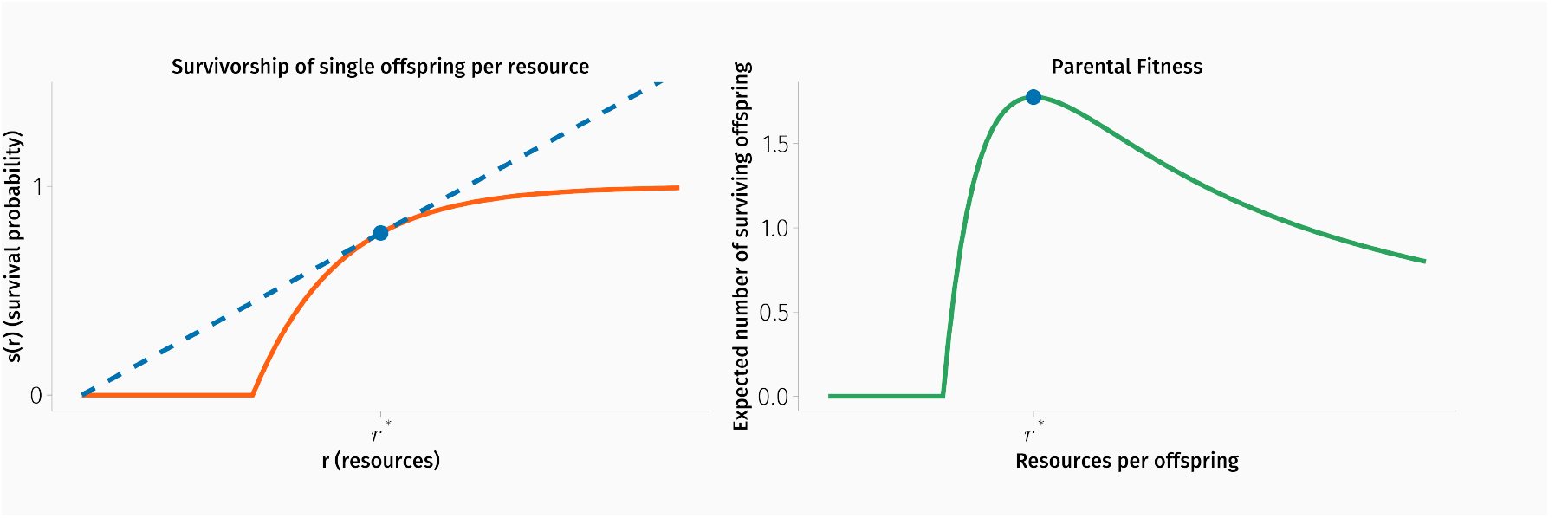
(Left) The survivorship function of the Smith-Fretwell model. This is a concave-increasing survivorship curve. The optimal offspring size can be found by drawing the blue dashed line from the origin so that it lies tangent to the curve. (Right) The fitness of a parent employing a symmetric strategy where the parent makes as many offspring of size *r*\* as possible. All offspring receive the same amount of resources.

**Figure 2.**
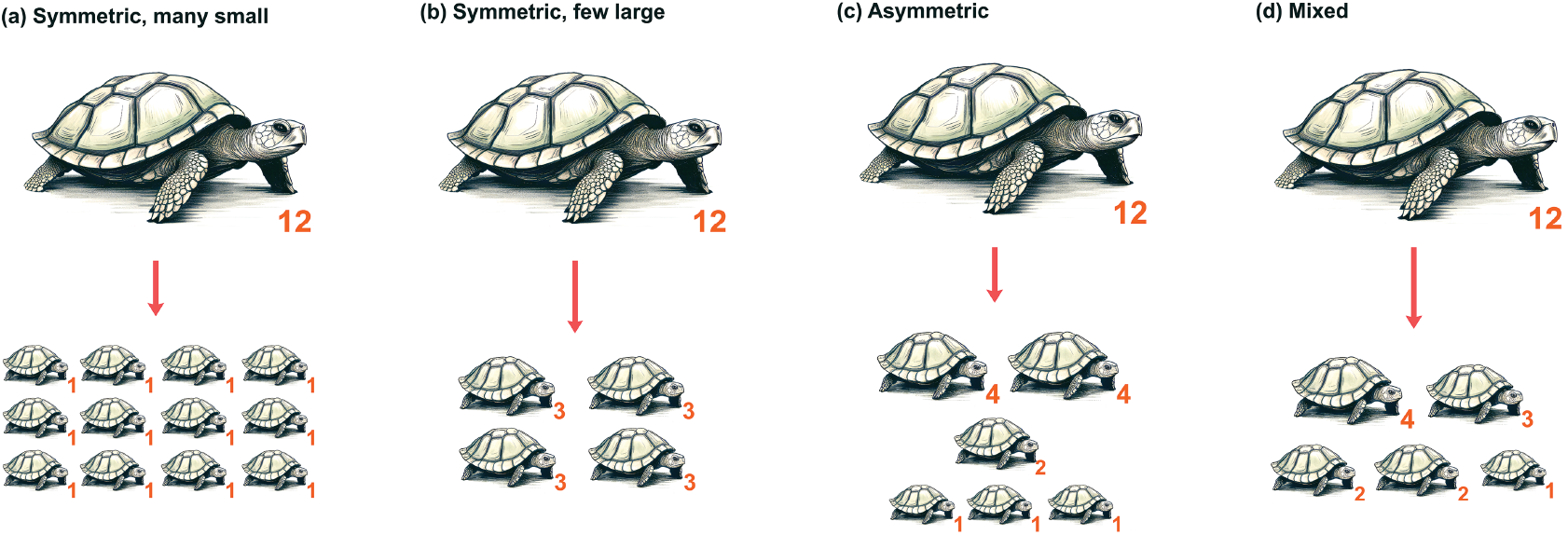
Four example strategies for distributing twelve resources when *r*_*low*_ = 1 and *r*_*high*_ = 4. The two figures on the left illustrate different symmetric strategies: in (a), all offspring are size one or zero, while in (b), all offspring are size three or zero. Figure (c) depicts an asymmetric strategy, in which there are many offspring of sizes *r*_*low*_ and *r*_*high*_ and at most one intermediate-sized offspring. Figure (d) demonstrates a mixed strategy, which is neither symmetric nor asymmetric.

These models also implicitly assume a *symmetric strategy* in which parents distribute resources equally among all their offspring (Roff, 1993, though exceptions exist, as seen in Marshall et al., 2008). This is a strong assumption, one which nature often ignores. In certain pea varieties and other plants, seeds located toward the middle of the pod tend to be larger (Harper et al., 1970; McGinley et al., 1987; Roach and Wulff, 1987). In animals, the offspring’s position in the ovary can affect size (Telfer and Rutberg, 1960; McLaren and Michie, 1960; McKeown et al., 1976). Even factors such as the number of adjacent eggs can impact the size of offspring, as observed in keelback snakes (Brown and Shine, 2009). Having offspring of different sizes may have evolved as a form of bet-hedging in uncertain environments, although the evidence here is mixed (Einum and Fleming, 2004; Marshall et al., 2008; Olofsson et al., 2009; Crean and Marshall, 2009; Morrongiello et al., 2012; Childs et al., 2010).

OOS models also assume that resources never harm offspring. However, if we generalize resources to be anything that is distributed to offspring, then we can include toxins like PCBs which accumulate within the blubber of marine mammals (Muir et al., 1996; Ross et al., 2000; Boon et al., 2002; Houde et al., 2006) and are offloaded to offspring during gestation and lactation (Aguilar and Borrell, 1994; Hickie et al., 1999; Klanjscek et al., 2007; Noonburg et al., 2010).

### B. Bacterial Aging (BA) Theory

Scientists had long believed that most cells distributed their components symmetrically. Each daughter would receive equal proportions of old and new components. Of course, some classic exceptions were known. *Saccharomyces cerevisiae* buds off small cells, with the larger cell inheriting old cytoskeleton and organelles (Herskowitz, 1988; Gershon and Gershon, 2000). *Caulobacter crescentus* asymmetrically divides into swarmer and stalked cells (Ackermann et al., 2003). Diatoms do not divide evenly, giving new cell walls to their younger daughters (Laney et al., 2012; Moger-Reischer and Lennon, 2019).

But more extensive research in asymmetric division has appeared only in the last three decades. Researchers observed subpopulations of *Escherichia coli* age and die, even with abundant resources. When *E. coli* reproduce, they expand their cell wall from one end, thus giving one daughter a new pole while the other daughter inherits the old pole. Damaged proteins segregate toward this old pole and can reduce the reproduction rate by more than 30% (Moger-Reischer and Lennon, 2019; Stewart et al., 2005; Winkler et al., 2010; Lindner et al., 2008). *E. coli* are not alone; asymmetric division has evolved repeatedly (Kysela et al., 2013; Ackermann et al., 2007). Damaged proteins accumulate with age in *Saccharomyces cerevisiae* and *Schizosaccharomyces Pombe* (Erjavec et al., 2008). Animal cells preferentially segregate genomic DNA, organelles, and damaged proteins into one cell (Fuentealba et al., 2008; Moore and Jessberger, 2017; Ogrodnik et al., 2014; Rujano et al., 2006; Moger-Reischer and Lennon, 2019). Cells also asymmetrically segregate RNA, histones, chromatids, mitochondria, and other organelles (Shlyakhtina et al., 2019). Polarization of the cell may enable this differential inheritance, and polarity itself may have arisen as a general strategy to restrict senescence to one daughter (Sunchu and Cabernard, 2020; Macara and Mili, 2008).

The disadvantage of asymmetric distribution is senescence. By consistently allocating old components into a single daughter, one lineage becomes a “garbage dump” for damaged components (Stephens, 2005). A cell eventually accumulates components so old and worn that they no longer support the functions of life. This unlucky daughter dies, as if from old age (Proenca et al., 2018). Empirical evidence suggests that such an asymmetric strategy increases population growth rate for certain species of bacteria (Winkler et al., 2010; Stephens, 2005; Kysela et al., 2013; Moger-Reischer and Lennon, 2019) and thus may have been selected for.

Models from the last two decades support this idea. In 2006, Watve demonstrated the evolution of asymmetry through a Leslie matrix stage-structured model (Watve et al., 2006) and in 2007, Evans did the same with a diffusion approach (Evans and Steinsaltz, 2007). In 2007, Ackermann tracked an abstract unit of damage that accumulates and passes on to offspring where damage decreased cell survival in a concave, convex, or linear manner (Ackermann et al., 2007). In a model by Erjavec et al., cells must accumulate a protein to a threshold to reproduce, but these proteins can degrade to damaged ones, slowing this process (Erjavec et al., 2008). Chao created a similar model, but where damage linearly reduced the rate of accumulation (Chao, 2010). Lin et al. tracked cell volume growth rate as a Hill-type function of the concentration of a key protein. They found a phase transition from symmetric to asymmetric strategies as the protein uptake rate changed (Lin et al., 2019). Unlike OOS models, these models consider the entire spectrum of strategies between symmetric and asymmetric. However, they are limited to two daughter cells, and thus cannot give any information on the optimal number of offspring.

#### Box 1.

The classic Smith-Fretwell model serves as the foundation of our model and makes the following assumptions.

1. Parents have a fixed amount of resources *R* to invest in reproduction.
2. Parents can choose the number of offspring *n* to produce.
3. The parent distributes resources equally among the offspring.
4. An offspring’s survival probability increases with resources, but with diminishing returns. The survival probability of an offspring given *r* resources is *s*(*r*).
5. Offspring require a minimum amount of resources *r*_*low*_ to survive.
6. Once an individual reaches maturity, its survival and reproduction are independent of the resource allocation it received as a juvenile.

Smith and Fretwell showed that there exists an optimal amount of resources (*r*\*) to give each offspring and that this amount maximizes fitness per unit resource. The parent’s optimal strategy is to produce as many offspring of size *r*\* as possible, yielding a total of *R/r*\* offspring. If this is not an integer, one simply compares the fitness of the two nearest integers and chooses the higher (Charnov et al., 1995). This optimal size is independent of *R* meaning that parents, when given additional resources, should create more offspring of size *r*\* rather than larger offspring. Note that this is an instance of the marginal value theorem (Charnov, 1976).

## 3. A General Resource Distribution Model

We model an abstract scenario without system-specific details. During reproduction, physical material is distributed among offspring. It may be food, space, time, attention, new cellular components, or even waste/refuse like toxins or damaged proteins. We generically call these *resources*, and they have two core properties:

1. The parent has only a finite amount of resources available to distribute.
2. An offspring’s probability of survival depends on how many resources it receives.

Parental fitness depends on their resource allocation strategy, and if the strategy is heritable, then populations will evolve to increase the expected number of surviving offspring.

We remove the constraints of OOS and BA to allow:

1. Any distribution strategies, from symmetric to asymmetric to anything in between.
2. Any integer number of offspring.
3. Arbitrary concave or convex survivorship functions, either increasing or decreasing.

### A. Model Setup

Suppose a parent has *R* total resources which it can distribute among their offspring. Parents control how they distribute their resources and the total number of offspring they create. We encode this information in the vector 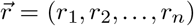, where offspring *i* receives *r*_*i*_ units of resource and *n* is the number of offspring created. We call 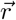 the parent’s *distribution strategy*. (For clarity we call anything the parent distributes *resources*, regardless of whether they help or harm the offspring. Mnemonically, one could think of *r* as standing for *refuse* when they harm. As shorthand, we sometimes refer to an offspring with *r* resources as being of *size r*.)

The parent’s expected number of surviving offspring for a particular strategy 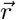 can then be written as

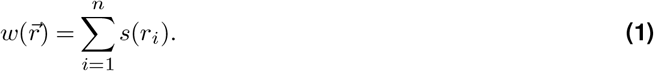

where *s*(*r*) is the survival probability of an offspring receiving *r* resources. Rather than restricting *s* to a specific functional form, we allow *s* to be an arbitrary continuous piecewise function made up of a curved region and two flat regions.

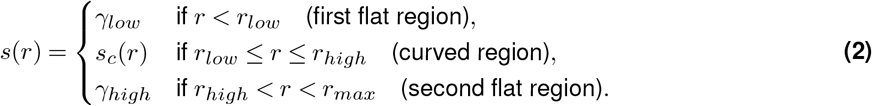

Figure 4 shows the six general shapes that *s* can take. Figure 3 contains a diagram of all the parameters and parts. All the information about how resources affect offspring survival probability is contained in *s*. Resources only affect survival; future performance is independent of how many resources an offspring initially received.

**Figure 3.**
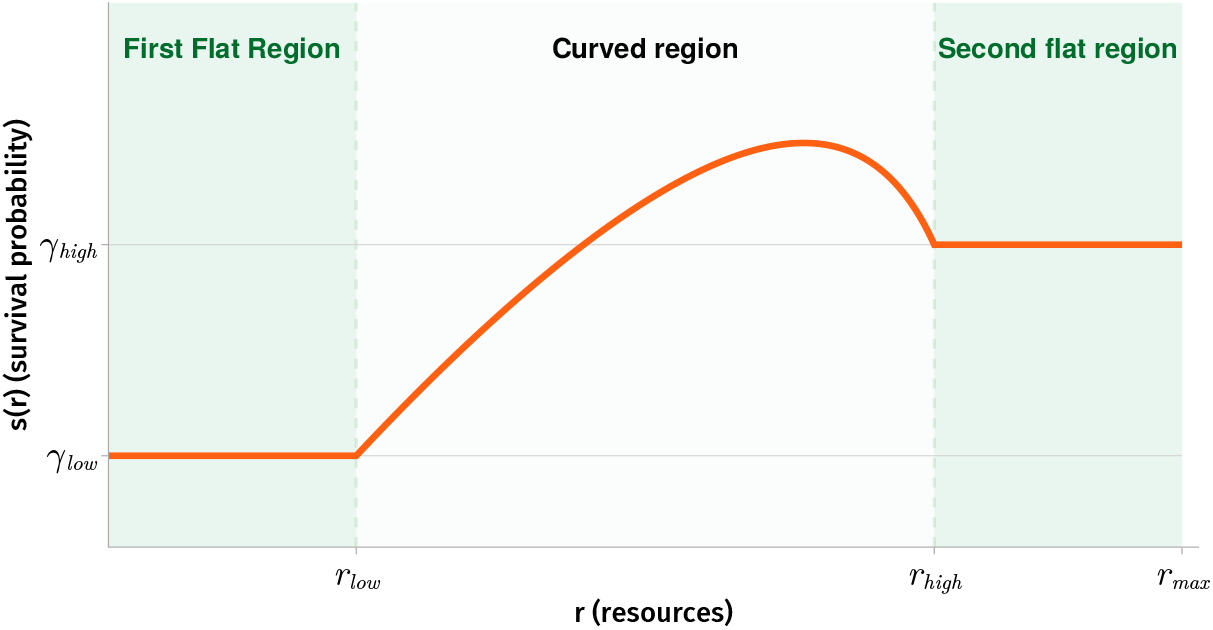
Anatomy of the survivorship function *s*. This function takes in a resource value between zero and *r*_*max*_ and outputs a survival probability. Note that *r*_*low*_ may be zero and *r*_*max*_ or *r*_*high*_ may be infinite. The constants *γ*_*low*_ and *γ*_*high*_ can take any value between zero and one.

**Figure 4.**
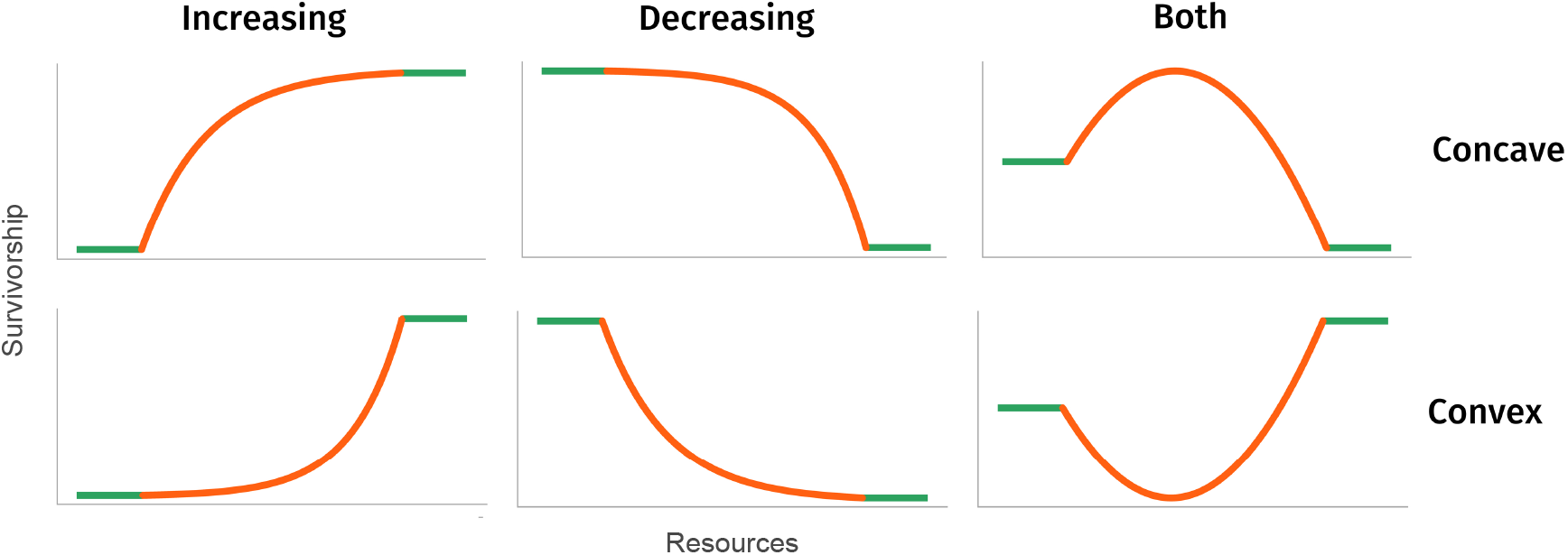
The six forms the survivorship function *s* can take, named according to the shape of their curved region (orange): concave-increasing, concave-decreasing, concave-increasing-decreasing, convex-increasing, convex-decreasing, convex-decreasing-increasing.

Equation 1 is a natural formulation of the resource distribution problem, though as far as we know this form has been analyzed only in Marshall et al., 2008. An optimal distribution strategy is an 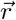 that maximizes 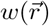 under one of the following constraints:

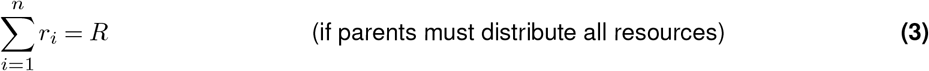

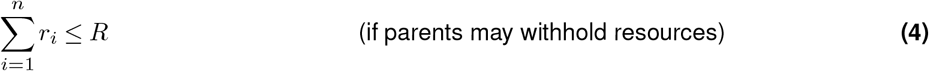

Which constraint we choose depends on the system. The first appears when the parent must use *all R* of their resources, such as when a bacterium distributes components, or if the resource is some kind of toxin that the parent must offload onto the offspring. The second constraint occurs when the parent can distribute *up to R* resources, but not necessarily all, such as when the resource is food, attention, or uterine capacity.

Furthermore, we limit the maximum size of each offspring to *r*_*max*_. Such a constraint could exist for many reasons, e.g., the offspring being too large for the mother to physically birth. If no such size limit exists, we set *r*_*max*_ equal to *R* because no offspring can receive more than the total amount of resources. The total number of offspring *n* may also be constrained. This constraint may have evolved in some species: carrying too many offspring could affect a fly’s ability to fly (Berrigan, 1991), a snake’s ability to move (Crews et al., 1988), or a lizard’s ability to climb (Andrews and Rand, 1974), hide in crevices (Vitt, 1981), and run (Garland Jr, 1985; Van Damme et al., 1989). Mathematically, we do not need to worry about these details; it is enough to fix the number of offspring at exactly *n*. Once we derive the form of the optimal strategy for any *n*, it is a simple matter to scan through all biologically reasonable values of *n* and select the one that yields the greatest fitness.

We assume natural selection acts to maximize the expected number of surviving offspring in constant environments, and that there is enough standing variation for this to happen. What natural selection maximizes (if anything) depends on many factors, including the genetic system, the levels of stochasticity, whether transgenerational effects exist, and whether the population is stage-structured, spatially structured, frequency dependent, or has overlapping generations (Proulx and Adler, 2010; Smith, 1978; Frank, 2011; Mangel et al., 1994). We further assume resources affect only offspring survival probability. This would be broken if adult state depended on initial resources received (e.g., resources affect size at maturity) or if resource distribution affected the parent (e.g. a parent starves or forgoes future reproduction to invest more resources into its offspring). We also ignore any evolution of the total resources *R*. However, if *R* did change as the parent ages, as expected in many organisms (Roff, 1993), then the analysis still holds in that the overall form of the strategy remains optimal (see Table 2 and Table 3) rather than the specific strategy itself (which depends on the exact *R* value).

To summarize, this model is a non-linear optimization problem where the solutions are the optimal resource distribution strategies that evolve. For a given number of offspring *n* we wish to find the distribution strategy 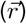 that maximizes parental fitness (Equation 1) while using up exactly all (Constraint 3), or no more than all (Constraint 4), total resources (*R*), without making offspring larger than the maximum size (*r*_*max*_).

#### A.1. Six shapes of survivorship

While in principle survivorship can be any arbitrary function, we restrict ourselves to the six possible classes shown in Figure 4 because they are both biologically relevant and mathematically tractable. The centerpiece of the survivorship function, both in location and importance, is the curved region given by *s*_*c*_. This is a concave or convex function that is continuous and twice differentiable everywhere except at the boundaries where it touches the flat regions. In the analysis section we will discover that the concavity of *s*_*c*_ is the main factor in determining the optimal strategy.

The first flat region of *s* describes what happens when resources are below the threshold *r*_*low*_. For example, *r*_*low*_ may represent the minimum amount of resources that an offspring needs for life; less than this and it is guaranteed to starve and die (Smith and Fretwell, 1974). Alternatively, if resources are damaged cellular components (as in bacteria), enough might need to accumulate before they begin harming the offspring. An offspring that receives below *r*_*low*_ resources has *γ*_*low*_ survivorship. Similarly *r*_*high*_ is the threshold after which resources saturate and no longer have an effect, and offspring above this threshold always have *γ*_*high*_ survivorship.

One can effectively remove the first flat region by setting *r*_*low*_ equal to zero. Likewise one can remove the second flat region by setting *r*_*high*_ equal to *r*_*max*_. This does not change any of the following analysis.

Concave-increasing survivorship captures diminishing returns of investment in offspring and thus is common in the OOS literature. When *r*_*high*_ and *r*_*max*_ are infinite, *s* is the same as a classic Smith-Fretwell survival function.

Concave-increasing-decreasing survivorship is seen in stage-structured populations. Sockeye salmon offspring have two developmental stages: egg and fry. The egg stage follows a concave-decreasing function. Large egg size decreases survivorship as the embryo requires a large surface-area-to-volume ratio to supply oxygen; too large an egg size, and the embryo asphyxiates and dies. But the opposite is true during the post-hatching stage; larger egg size correlates with larger fry and thus higher survivorship in a concave-increasing manner. When combined, these two forces create a concave-increasing-decreasing survivorship function (Hendry et al., 2001 but see Hendry and Day, 2003).

Concave-decreasing and convex-decreasing survivorship are both seen in BA. These decrease because the mother distributes a waste product to offspring. These could be toxins or damaged proteins in the cell, but more often are old cellular components. There tends to be a threshold amount where, once passed, they start harming the offspring.

Where convex-increasing and convex-decreasing-increasing survivorship arises is more speculative. Convex-increasing survivorship may apply where, for example, size follows a sigmoidal survivorship curve, but the inflection point is beyond the physiological limits of what the mother can birth. The survivorship curve, from the mother’s perspective, is effectively convex. For convex-decreasing-increasing survivorship, it would be associated with something that is beneficial in high and low doses, but detrimental in medium doses. For example, a small offspring may be able to hide from predators while a large offspring can fight them off, but a medium-sized offspring is more vulnerable to predation.

### B. Analysis

We have two central questions: what strategies are optimal, and will organisms evolve toward these optimal strategies? Our analysis strategy is as follows.

1. Restrict offspring to the curved region of *s*. Using the convexity of the curved region, apply Jensen’s inequality to find the optimal distribution strategy, *when restricted to the curved region*.
2. Remove the restriction to allow the two flat regions. Work through each of the six survivorship classes to arrive at the specific optimal distribution strategies for each class.

In all cases, we will find that the fitness landscape is simple and “Mt. Fuji-like”, meaning the optimal strategy is unique (up to some trivial cases). The optimal strategies are summarized in Table 2 and Table 3

To streamline the analysis, we focus on the case where the curved region *s*_*c*_ is *strictly* concave or convex. This ensures that the optima we derive are unique. The same analysis works for the non-strict case, but the optima need not be unique or strict. Natural selection may instead become stuck at different optima or neutrally drift along flat regions.

### C. Convexity drives Symmetry vs Asymmetry

Here we derive our two central results.

1. Concave survivorship functions select for **symmetric** distribution, in which parents make a set number of offspring and distribute resources equally between them.
2. Convex functions select for **asymmetric** distribution, in which resources are distributed as unevenly as possible. This can create a group of small offspring and a group of large offspring, with potentially a single middle-sized offspring. Alternatively, this can create a group of large offspring with a single runt.

Convexity causes different outcomes due to Jensen’s inequality, which controls evolution for offspring on the curved region *s*_*c*_. This allows us to derive the optimal strategy if all offspring were limited to the curved region, and to prove that populations evolve toward our optimal strategy. Because this depends only on the convexity, it does not matter whether the survival functions are increasing or decreasing.

#### C.1. Symmetric Strategies are Optimal under Concave Survival

Suppose *s* is a concave-increasing function.

Consider an optimal distribution strategy 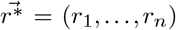. Each offspring must fall into one of three categories:

1. offspring on the first flat region of *s* (who receive 0 ≤ *r < r*_*low*_ resources)
2. offspring on the curved region *s*_*c*_ (who receive *r*_*low*_ ≤ *r* ≤ *r*_*high*_ resources)
3. offspring on the second flat region of *s* (who receive *r*_*high*_ *< r* ≤ *r*_*max*_ resources).

If we decompose the fitness contribution into a term for each of these categories, the parental fitness (Equation 1) becomes

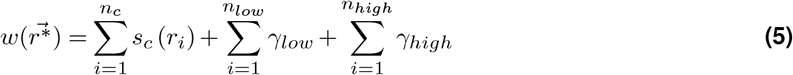

where *n*_*c*_, *n*_*low*_ and *n*_*high*_ are the number of offspring on the curved, first flat, and second flat regions respectively. As all offspring on the flat regions have the same survival probability, this simplifies to

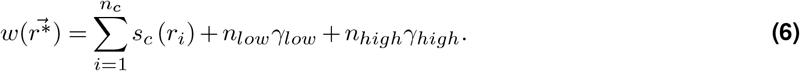

Focus now on the summation term 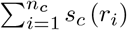. Because *s*_*c*_(*r*) is a strictly concave function, Jensen’s inequality states that

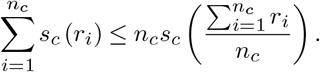

Then, defining *R*_*c*_ as the total amount of resources invested in the curved-region offspring, we can rewrite this as

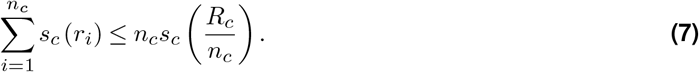

For a strictly concave function *s*_*c*_, equality holds if and only if all *r*_*i*_ are identical (i.e., *r*_*i*_ = *R*_*c*_*/n*_*c*_ for all *i*). This implies that any optimal strategy *must* equally distribute the *R*_*c*_ resources to the *n*_*c*_ curved-region-offspring. This is a *symmetric* distribution strategy and looks like

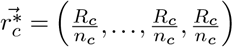

where 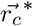 denotes the optimal strategy for offspring on the curved region.

Here, a symmetric strategy is the unique and strict global optimum when considering only the offspring on the curved region. This is because, if a multivariate function is strictly concave, and if the collection of feasible strategies lies on a convex set, then there is a single extremum that is strict. Our constraints are linear, and so the feasible strategies lie on a linear set that is by definition convex. If the curved region is instead *non-strictly* concave then the symmetric strategy remains a global optimum but it is no longer unique or strict. As an extreme example, if *s*_*c*_ is linear so that *s*_*c*_(*r*) = *r*, then all ways of distributing the offspring on the curved region perform equally well (Hillier and Lieberman, 2010).

The fact that there is only one optimum under strict concavity hints that the fitness landscape is a simple, single-peaked one. This would imply our population evolves toward the optimum from any starting point. We show that this is indeed the case in Supplement A. For fixed *R*_*c*_ and *n*_*c*_, mutations that move the curved region’s offspring towards the symmetric distribution are beneficial. These mutations do not have to be small; a large mutation is beneficial as long as it makes the distribution more symmetric. (Or, if *s*_*c*_ is non-strictly concave, such mutations are at worst neutral.)

Note that we have not yet shown what values of *R*_*c*_ and *n*_*c*_ give the optimal strategy. All we know is that, given an *R*_*c*_ and *n*_*c*_, the optimal strategy is symmetric.

#### C.2. Asymmetric strategies are optimal under convex survival

For strictly convex survivorship, we mimic the argument for the concave case. Split the offspring into three types: those on the two flat regions and those on the curved region. Consider only those on the curved region and again apply Jensen’s inequality. The inequality sign is reversed because *s*_*c*_ is strictly convex, and we conclude that

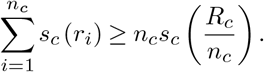

For a strictly convex function *s*_*c*_, equality holds if and only if all *r*_*i*_ are identical. This is to say that the symmetric strategy performs worse than all other strategies. Furthermore, it is strict and is the unique global minimum for the sum ∑*s*_*c*_(*r*_*i*_). This is because, if a multivariate function is strictly convex and the collection of feasible strategies lies on a convex set, then there is a single extremum, and it is strict (Hillier and Lieberman, 2010; Boyd and Vandenberghe, 2004).

This is the opposite of the concave case. Mutations that move the strategy *away* from symmetry – and thus make the strategy more *asymmetric* – are selected for. If *s*_*c*_ is non-strictly convex, such mutations are at worst neutral. Climbing the fitness landscape through a series of small mutations, we end up at a distribution where the curved region’s strategy has the form

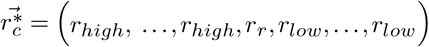

where *r*_*low*_ ≤ *r*_*r*_ ≤ *r*_*high*_. We call such a strategy an *asymmetric strategy*. We can produce this strategy with a simple algorithm: give all offspring *r*_*low*_ resources, then sequentially make as many *r*_*high*_ sized offspring as possible, and finally give any remaining resources *r*_*r*_ to a single offspring.

This strategy is a strict global optimum, is unique (up to reorderings), and we expect it to evolve. We have placed the proofs for these statements in Supplement B, as they are tedious and unenlightening.

### D. Optimal strategy for specific cases

The convexity of the survivorship function determines whether resources for offspring on the curved region should be distributed symmetrically or asymmetrically. However, we do not yet know how many offspring or resources should initially be placed on the curved region, nor how offspring placed on the flat regions influence the optimal solution. To resolve this, we must analyze each case in Figure 4 individually. The details of these analyses, along with algorithms used to calculate the optima, are given in Supplement A.

We have placed these derivations in the supplement because the additional derivations contain no major biological insights. They are lengthy and primarily involve working through edge-cases that arise from considering the flat regions. Generally, we find a unique optimum for non-extreme values of *R*. At extreme values, however, all offspring may be situated exclusively on the lower or upper flat regions, resulting in multiple equivalent optimal distribution strategies.

We also studied when these results hold in stage-structured populations. We have confined this analysis to Supplement D due to scope.

## 4. Fluctuating Environments

In constant environments, our model predicts that mixed strategies never arise; evolution prefers purely symmetric or purely asymmetric strategies. However, mixed strategies exist in nature. This may be due to sibling competition, spatial constraints, parasitism, differing offspring quality, or imperfect information, to name a few. Here we focus on one of the most common explanations: that fluctuating environments have selected for diversifying bet-hedging (Roff, 1993; Crean and Marshall, 2009; Olofsson et al., 2009).

If strategies are nonplastic, the parent is forced to commit to a single strategy across all environments it encounters. Mathematically, the parent chooses an 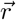 and *n* that remain constant across all environments they encounter. The optimal strategy will be the 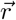 that maximizes the geometric mean fitness 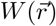 (Haldane and Jayakar, 1963; Karlin and Liberman, 1975; Proulx and Adler, 2010; Frank, 2011; Proulx and Teotónio, 2017; Proulx et al., 2019).

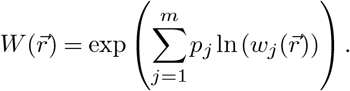

Here *p*_*j*_ is the frequency of encountering environment *j* out of the *m* possible environments, and 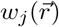 is the parental fitness of applying strategy 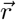 in environment *j*.

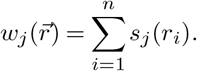

As before, *n* is fixed, *i* indexes the offspring, and parents cannot allocate more than the total resources available in a given environment. Let *R*_*j*_ denote the total resources in environment *j*. Rather than track *R*_*j*_ explicitly, we measure allocations as proportions so that *r*_*i*_ is the proportion of total resources given to offspring *i*. After this rescaling, every environment has *R* = 1, and *s*_*j*_(*r*_*i*_) is the probability that an offspring in environment *j* survives when it receives that fraction of its parent’s resources. Environmental variation now enters only through the functions *s*_*j*_, simplifying the analysis.

**Table 4.** Additional terms and symbols for the fluctuating environments model. Note that *j* indexes the environments and *m* is the total number of environments while *i* still indexes offspring and *n* is the total number of offspring.

| Term | Definition |
| --- | --- |
| $s_j(r)$ | The survivorship of an offspring given $r$ resources in environment $j$. |
| $w_j(\vec{r})$ | The fitness of distribution strategy $\vec{r}$ in environment $j$. |
| $W(\vec{r})$ | The geometric mean fitness of distribution strategy $\vec{r}$ across all environments. |
| $m$ | The number of environments |
| $p_j$ | The probability of environment $j$. |

### A. Analysis of Fluctuating Environments

We analyze this model in three parts, building in complexity. The analysis here is informal – otherwise one requires an unenlightening mess of notation and definitions. We quarantine the formal derivation of the general case to Supplement A.

#### A.1 Environments share resource thresholds and convexity

Assume that the thresholds *r*_*low*_ and *r*_*high*_ remain the same across all environments. We can then define a curved section of the full fitness landscape *W* by restricting all the *r*_*i*_ to be between *r*_*low*_ and *r*_*high*_. Denote this restricted fitness landscape as *W*_*c*_.

*W*_*c*_ is concave if all the *s*_*j*_ are concave, as in Figure 5(a). This is because *W*_*c*_ is the composition of the weighted geometric mean (a non-decreasing concave function) with the parental fitness for each environment *w*_*j*_ (a concave function on the restricted domain, as it is the sum of concave *s*_*j*_).

**Figure 5.**
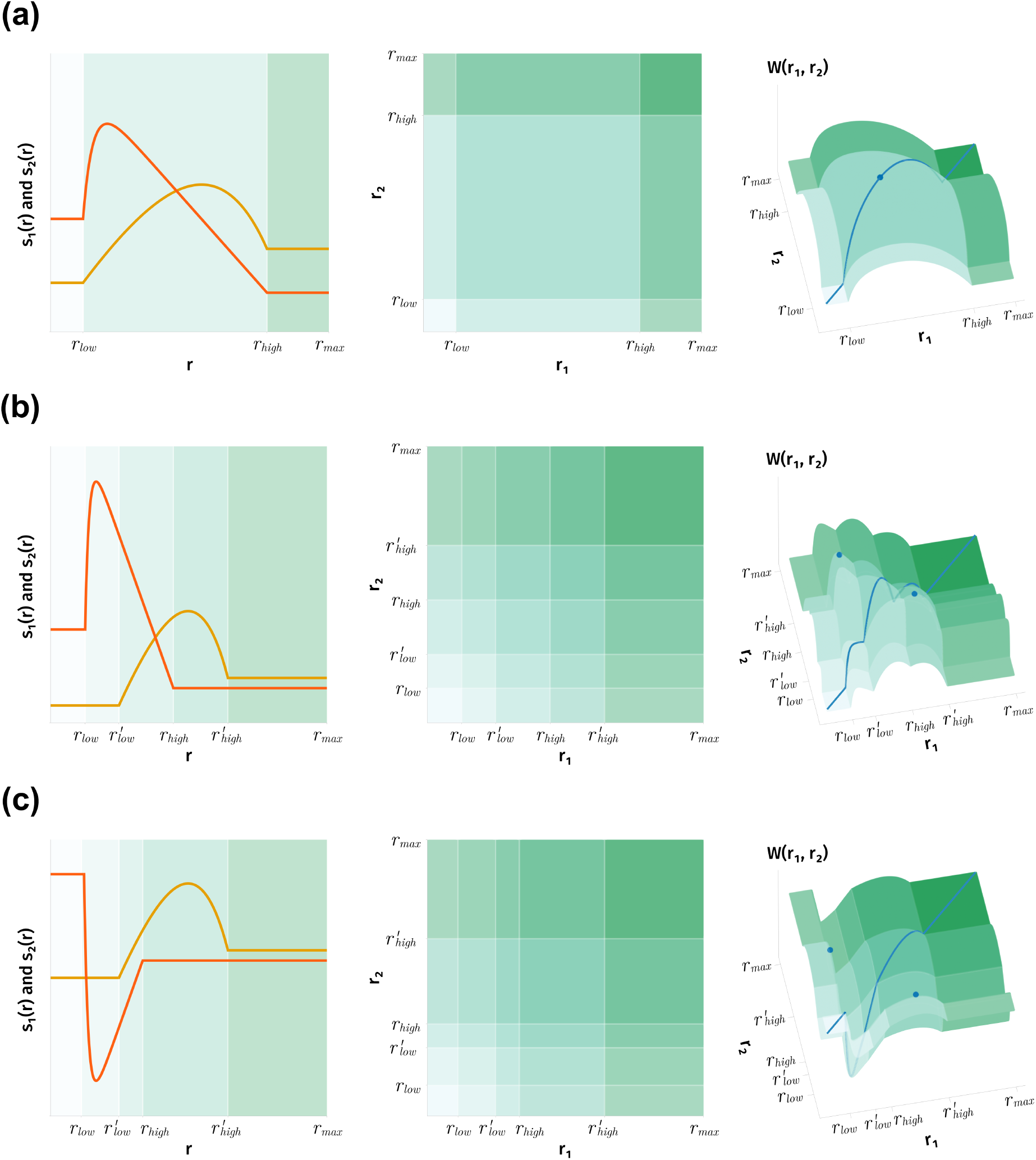
Each row shows a system with two offspring and two equally probable environments. The dark orange survivorship curve corresponds to the first environment and the light orange survivorship curve to the second. Along the axes, the ticks of the second environment are denoted by primes. In (a), the survivorship functions are both concave and share resource thresholds – that is, boundaries of their curved regions (*r*_*low*_ and *r*_*high*_). In (b), they are both concave but do not share boundaries. In (c), they do not share convexity nor boundaries. The first column illustrates how the thresholds at 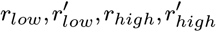 define regions. The second column shows how these regions divide up the space of possible strategies – in the case of two offspring this is a two-dimensional grid. For example, the cell in the upper left corner of each grid represents all strategies where the first offspring has less than *r*_*low*_ resources and the second offspring has more than 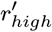 or *r*_*high*_. The third column shows the fitness landscape *W*. Here (a) and (b) demonstrate how these cells map exactly onto concave regions when both the survivorship functions are concave. The blue dots are the global optima. The blue line represents all symmetric strategies. Each point of this line is always a local optimum when restricted to the slice of the landscape that uses the same amount of resources.

As in the constant environment case, a concave *W*_*c*_ implies the optimal strategy is symmetric. Given any non-symmetric strategy within this curved region, we can make two of the offspring more symmetric without reducing geometric mean fitness. By Jensen’s inequality applied to each *s*_*j*_, this action does not decrease each *w*_*j*_ and thus does not decrease *W*_*c*_ (since the geometric mean is non-decreasing in its arguments).

Thus, the “concave survivorship implies symmetric strategies are optimal” result remains robust, even in the face of fluctuating environments. Bet-hedging does not occur under these conditions, and the parent makes as many offspring of the optimal size as possible.

This contrasts with the classic result that bet-hedging evolves if *s*_1_ and *s*_2_ are Gaussian survivorship curves such that their peaks are at different points and they are narrow enough that they are sufficiently non-overlapping. We can reconcile this with our result by observing how immensely restrictive our two conditions are. All the *s*_*j*_ must have the same *r*_*high*_ and *r*_*low*_, and all the *s*_*j*_ must be concave within this shared range. This prevents us from having two different peaks and low overlap at the same time. The *s*_*j*_ being concave on [*r*_*low*_, *r*_*high*_] always forces a large amount of overlap. (E.g., if one places peaks as far apart as possible, one peak at *r*_*low*_ and the other at *r*_*high*_, we would get something akin to two mirrored right triangles, ensuring high overlap.) This does not happen if the *s*_*j*_ are Gaussian – the long convex tails can sufficiently reduce overlap.

Conversely, if *W*_*c*_ is convex, a similar argument as in the concave case can show that an asymmetric strategy must be optimal. However, if each of the *s*_*j*_ is convex, *W*_*c*_ is not *guaranteed* to be convex. There exist examples where convex *s*_*j*_ induce a sigmoidal *W*_*c*_. In such cases, the sigmoidal shape can be subtle and appear convex to the naked eye.

#### A.2. Environments share convexity but not resource thresholds

Suppose that all the individual environmental survival functions *s*_*j*_ are concave, but their thresholds *r*_*low*,*j*_ and *r*_*high*,*j*_ differ across environments, as in Figure 5(b). The curved regions of survival functions may overlap partially or not at all. In this case, the full fitness landscape *W* is made up of many concave sections.

We can view each *s*_*j*_ as being made up of a first flat region, its curved region, and a second flat region. Consider an arbitrary strategy 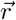. For each environment *j*, each *r*_*i*_ falls on either the curved region or one of the flat regions of *s*_*j*_. Imagine now restricting each *r*_*i*_ to remain in the region (flat or curved) it initially occupied for each *s*_*j*_ simultaneously. Geometrically, we can visualize these regions by partitioning the resource allocation space. Each axis (representing an *r*_*i*_) is segmented by all unique *r*_*low*,*j*_ and *r*_*high*,*j*_. This creates a multi-dimensional grid of cells, each corresponding to a distinct section of *W*, as shown in the middle column of Figure 5.

This partitions the domain of possible strategies into cells. Within each cell the fitness landscape is concave because it is a composition of concave survivorship segments with the geometric mean. This preserves concavity, as discussed in the previous case. (Recall the flat regions do not disrupt this, as the constant function is both concave and convex).

This paints *W* as a rugged landscape, but one with predictable regularity. If all the *s*_*j*_ are concave, the fitness landscape within each cell is a hill, and a 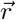 starting within a cell is expected to stay in that cell, evolving toward the top of that cell’s hill. Cells thus act as evolutionary basins of attraction. There are between 1 and at most (2*m* + 1)^*n*^ of these cells.

If the *s*_*j*_ are instead convex, then *typically* we have fitness valleys and basins of repulsion within the cells. The strategy is then expected to evolve to the border of the cells and move along it. However, this is not always true, as it is not guaranteed that the fitness landscapes within these cells are convex (since the geometric mean of convex functions is not necessarily convex). Many open questions remain for this case.

#### A.3. Environments share neither resource thresholds nor convexity

Consider a mix of concave and convex *s*_*j*_ with no shared thresholds, as in Figure 5(c). As before, we can split each *s*_*j*_ into flat and curved regions to form cells.

The fitness landscape here is a complex world of mountains, valleys, and rolling sigmoidal hills. Yet despite this complexity, symmetric strategies are always extrema when restricted to the set of strategies that use *exactly the same amount of resources*. Symmetric strategies may shift from being a minimum to a saddle point or a maximum, but they always remain local extrema.

This is due to the *Purkiss Principle*, which (loosely) states that symmetric problems have symmetric solutions. Below is the principle as written in Waterhouse, 1983. (The source contains far stronger and more general versions of this theorem, but we only require the basic one.)

##### Purkiss Principle

*Let* 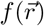 *and* 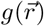 *be symmetric functions (i.e*., *the order of r*_*i*_ *does not matter) and consider some vector* 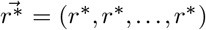 *with all entries equal. Say that f and g both have continuous second derivatives in the neighborhood* 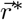.*Then, when constrained to vectors* 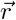 *such that* 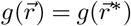, *the function* 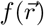 *will have a local maximum or minimum at* 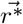 *as long as*

1. 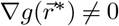.
2. *All second order terms of f* −*λg, are non-zero in directions perpendicular to* 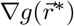, *for all constants λ*.

For us, *f* represents the geometric mean fitness *W* and 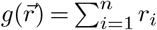 represents our resource constraint. These are both symmetric functions that meet the conditions above, thus the Purkiss Principle applies. If we limit *n* to be the number of non-zero-sized offspring, we can also show that *any* symmetric strategy is a local extremum when restricted to non-zero sized offspring.

## 5. Discussion

OOS and BA both study the same problem of resource distribution, yet their literature rarely intersects. This is perhaps unsurprising; they arose from different parts of biology and make distinct mathematical assumptions about survivorship and the number of offspring allowed. Here we have abstracted away the system-specific details to build a more general and unifying model, showing that concave survivorship implies symmetric distribution and convex survivorship implies asymmetric distribution. This framework:

1. agrees with and strengthens predictions of existing models.
2. organizes existing models into a unifying framework.
3. makes new predictions.
4. reveals universal patterns in resource distribution models.

Concerning point (1), previous results still hold in our more general setting. We have shown the Smith-Fretwell model’s implicit assumption that a symmetric strategy is optimal does not invalidate its conclusions under concave survivorship. We have shown, in BA, that asymmetric strategies can evolve without requiring binary fission or damage accumulation across generations (Angert, 2005).

Furthermore, we show that any mutation that places the strategy closer to optimal is selected for. This implies not only that optimal strategies evolve via small mutations (something not always explicitly shown) but that they can also evolve by large mutations or recombination. Even in the face of reproductive constraints, we expect evolution to get as close to the optimal strategy as possible. In any given organism, symmetric and asymmetric strategies stand at the extremes of fitness. One is the best strategy, and the other must be the worst (Supplement E). We have also shown which of these results carry over to fluctuating environments.

Our model also (2) organizes existing models into a unifying framework. Counterintuitively, whether resources help or harm the offspring matters less than whether they affect the offspring in a concave or convex manner. Convexity determines whether symmetric or asymmetric strategies are selected for.

This shifts the biological focus. It is not enough to know whether resources help or harm. We need to understand the convexity. The upside is a powerful tactic: assuming our model assumptions hold, we can ignore much information. For example, to decide the optimal strategy for the convex-increasing and convex-decreasing cases, we only require *r*_*low*_, *r*_*high*_, and *r*_*max*_. The rest of the survival function is irrelevant except for determining the fitness of the single runt.

The idea of runts transitions us to how our model (3) makes new predictions. One of which is that resource distribution is an evolutionary mechanism selecting for the birth of runts (a single, significantly smaller, offspring appearing in a litter). The “runt of the litter” in dogs is common in popular culture, and the frequency of runts has been studied in pigs (van der Lende et al., 1990). Yet the literature on runts is sparse, and we found no studies proposing an adaptive reason why a single runt would evolve. We can now give a reason: creating a single runt maximizes expected fitness under convex-increasing survivorship.

However, it is unclear where such convex-increasing survivorship occurs in nature. One possibility is found at the start of a sigmoidal function. Sigmoidal functions arise when we allow for sibling rivalry and parent preference for the larger offspring (Dani and Kodandaramaiah, 2017; Jørgensen et al., 2011). In this case, if a mother is so starved of resources that she cannot or will not create any offspring on the concave region, then she is effectively working with convex-increasing survivorship. In this specific scenario, we predict that increasing total resources will create a “phase transition”, where the mother abruptly switches from an asymmetric strategy to a symmetric one.

Another prediction concerns cases where animals vertically transmit toxins to their offspring. Assuming toxins induce concave-decreasing survivorship, we expect the mother to produce two classes of offspring – those with large amounts of toxins and those with few toxins. This pattern can be seen in a dynamic energy budget model of the North Atlantic right whale (Klanjscek et al., 2007).

A third prediction is that convexity alone is enough to evolve asymmetric reproduction in bacteria. We do not need to follow the age of the components or the accumulation of damage over multiple generations. Local information about survival after one generation is enough.

Imagine a bacterium that, upon birth, immediately repairs any old components it inherited to the “new” state. The energy cost of this initial repair induces a convex survivorship function. After this initial repair, it allows its components to age normally so that, by reproduction, its components are “old”. The bacterium duplicates its components and divides. In this case asymmetry still evolves, even though components never grow older than one generation. This is not to say component aging and repair can be ignored, but rather that the shape of the survivorship function (i.e., its convexity or concavity) alone implies a default evolutionary trajectory which component aging and other effects can then modify.

For a fourth prediction, we note that bacteria may have originally divided symmetrically, and it is only with the evolution of asymmetry that the concept of “parent” emerged (Ackermann et al., 2003; Ackermann et al., 2007). But if symmetry was the historical stable state, then survivorship may have originally been concave, with symmetric division being a local optimum. To evolve asymmetric division by small mutations, the survivorship function may have become convex. We would then predict that the evolution of symmetric to asymmetric division is then the evolution of a concave survivorship to a convex one.

A final, speculative, prediction is that we can select for specific strategies in real systems by manipulating the convexity of the survivorship function. For example, enforcing concave survivorship could reduce variability in farming applications, and enforcing a convex function could slow the runaway symmetric division found in stem cell cancers (Clevers, 2005; Morrison and Kimble, 2006; Wodarz and Gonzalez, 2006; Clevers, 2011; López-Lázaro, 2015; López-Lázaro, 2018; Shlyakhtina et al., 2019).

Analysis of our model also (4) reveals universal patterns in resource distribution models. In particular, symmetric strategies frequently emerge as local extrema. This reflects the deep mathematical symmetry of resource distribution models: all offspring play by the same rules and take from the same resource pool in the same manner. Mathematically, this means the parental fitness and resource constraint are symmetric functions. The Purkiss Principle then tells us (with caveats) that symmetric strategies are extrema. Even if offspring mutate or migrate to different regions, they share rules in how they mutate or move. Organisms following the same rules, interacting with the environment in the same way, is an almost universal simplification in evolutionary and ecological modeling. This implies symmetry, and hints that the generalized versions of the Purkiss Principle may apply widely.

### A. Future Directions and Open Problems

While this convexity framework unifies OOS and BA, we have ignored transgenerational accumulation of damage, a cornerstone of BA research. Specifically, our model treats components as “new” or “old.” We do not track how they become progressively older and more damaged over generations. Rather, we show that such transgenerational effects are not *required* for symmetric or asymmetric strategies to emerge. These effects could still reinforce or change the predicted optimal strategy (and some early pilot simulations suggest this happens in nontrivial ways). Old degraded components could even change the ability to reach the optimum; aging has been shown to reduce a cell’s ability to divide asymmetrically (Lai et al., 2002; Moger-Reischer and Lennon, 2019).

However, this disregard for transgenerational damage may still approximate certain bacterial systems. Suppose a bacterium’s survivorship depends on its internal burden of damaged proteins. The older the protein, the more damage it contributes. While daughter cells might inherit varying initial damage loads, cellular processes like protein recycling, repair, and dilution via growth can lead the total quantity of such damage to converge towards a steady state (Erjavec et al., 2008). If division is triggered when cells reach a certain physiological state (e.g., have damage load beneath a certain threshold per unit volume), then most parent cells would possess a similar, characteristic total amount *R* of this “damage load” to distribute. Our model’s assumption of a fixed *R* for the dividing parent captures this scenario.

Transgenerational effects are not limited to bacteria either. Larger offspring may mature faster and gain more resources (Roach and Wulff, 1987; Mousseau and Dingle, 1991; Roff, 1993; Donohue, 2009). Current models of offspring size often ignore this aspect by assuming an offspring’s fitness as a parent is independent of how many resources they were given as a child. To fix this, we could incorporate transgenerational effects in the style of BA models.

Our model also lacks repair. As an alternative to distributing damaged components, cells may repair them at the cost of energy (Ackermann et al., 2007; Watve et al., 2006; Evans and Steinsaltz, 2007; Proenca et al., 2018). With repair, one can map out at which energy cost the optimal strategy switches from the symmetric to the asymmetric strategy. We do not consider repair in our model, again being content to show it is not *required* to evolve symmetry or asymmetry. But repair may reinforce or modify the predicted course of evolution. Repair might also extend to OOS theory; vertical transfer of toxins to offspring has been seen in butterflies Paula et al., 2014, minks (Janelle C. Restum John P. Giesy James A. Render Elizabeth B. Shipp David A. Verbrugge Richard J. Aulerich, 1998), and marine mammals (Hickie et al., 1999; Klanjscek et al., 2007). The parent may be able to spend resources or energy to remove such toxins before they are transferred (Noonburg et al., 2010).

We also ignored environmental stress, which increases the rate of cellular damage and is known to favor asymmetric division (Vedel et al., 2016). We postulate that rising stress is equivalent to morphing the survivorship function to become progressively more convex. Fluctuating environmental stress would then be equivalent to our fluctuating environments model.

Incorporating the interplay of transgenerational effects, repair, environmental stress, and other effects would further unify BA and OOS theory. And even then the work is not done – our approach is fundamentally limited because optimization models rarely tell the full story once we add in environmental feedback (Parker and Smith, 1990; Bernardo, 1996; Metz et al., 2008b; Metz et al., 2008a).

Further open problems surround how our results change under various modifications. We can let the survivorship be sigmoidal or Gaussian, or allow the order of offspring to matter. We can also add stage-structure under fluctuating environments or consider systems with multiple resources rather than one. These extensions, and their biological motivation, are discussed in Supplement A.

## 6. Conclusion

We have unified and expanded the theories of Optimal Offspring Size (OOS) and Bacterial Aging (BA) through a resource distribution framework that leverages the powerful and well-developed theory of convex optimization. In constant environments, we classified optimal distribution strategies and showed that they evolve. These findings highlight the central role of convexity: concave survivorship selects for symmetric distribution while convex survivorship selects for asymmetric distribution. We partially extended these principles to fluctuating environments.

There remains much to explore. Future research should incorporate key BA mechanisms like transgenerational effects, repair, and environmental stress responses. This will strengthen the bridge between OOS and BA, fostering the flow of ideas and insights between the two fields.

## Supporting information

Convex Patches Counterexample

## Acknowledgments

We thank Alexandra Brown for figure design advice and the fruitful suggestion to use Jensen’s inequality. Roger Nisbet provided many excellent edits and references. Holly Moeller and Kelly Thomasson gave helpful discussions. Lina Kim and University of California, Santa Barbara’s Research Mentorship Program matched us with excellent student research assistants. Of these, Arjun Patrawala, Ruchi Dixit, Heather Wei, and Kevin Guo performed early pilot simulations and studies for the main model. Victor Zhou provided the tentative results on transgenerational effects. Kaival Shah and Austin Park helped create the conjectures in Supplement F.1 and Supplement F.2 respectively.

## Supplementary Information

### Supplement A. Optimal strategies for constant environments

#### A. Concave-increasing

Concave-increasing functions are common in optimal offspring size theory. The classic Smith-Fretwell survivorship function is an example of such a function for the special case where *r*_*max*_ equals *R*. Such functions model resources that are always beneficial or neutral.

Because more resources never harm the offspring, any proposed optimal strategy that uses less than *R* resources could be improved – or at least not made worse – by distributing all *R* resources. Thus we can focus only on strategies that use all *R* resources because a globally optimal strategy exists among them.

Any strategy partitions the offspring into one of three regions: the first flat region, the curved region, and those on the second flat region. If *R* is high enough that we can distribute the resources so that all *n* offspring receive *r*_*high*_ or greater resources, then any such distribution is an optimal strategy and we are done. Otherwise, focus on the first flat region. An offspring in this region with size above zero is wasting resources – we could make that offspring size exactly zero without decreasing its survivorship and use those resources on other offspring. Likewise it is never benificial to give an offspring more than *r*_*high*_ resources because the resources above *r*_*high*_ could be given elsewhere without decreasing fitness. In other words, if we have a strategy that gives offspring *i* a resource value 0 *< r*_*i*_ ≤ *r*_*low*_ or *r*_*i*_ *> r*_*high*_ then we can replace *r*_*i*_ with zero or *r*_*high*_ respectively. We can then reallocate the remaining resources someplace else to obtain a strategy with the same or better fitness.

This implies that an optimal strategy lies among those strategies where the offspring in the first flat region are all exactly size zero and offspring in the second flat region are all exactly size *r*_*high*_. Note that offspring of size *r*_*high*_ are technically not part of the flat region, but are (just barely) on the curved region. Thus the second flat region has no offspring.

From our analysis of concave functions in the main paper, we already know that an optimal strategy distributes resources among curved region offspring symmetrically. This means at least one optimal strategy has only zero sized offspring on the first flat region, no offspring on the second flat region, and offspring on the curved region all having the same size. Call such a strategy 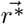. From Equation 6, we see the fitness of this strategy is

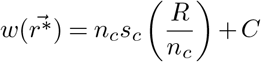

where *C* = (*n* − *n*_*c*_ − 1)*γ*_*low*_ is the fitness contributed by all the zero resource offspring.

As a technical point it is never a bad idea for a parent to make 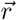 as large (in length) as possible because *γ*_*low*_ ≥ 0. But biologically speaking *γ*_*low*_ is usually zero as zero resource offspring have no chance of survival. In this case, regardless of the length of 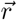, we would say *n*_*c*_ is the actual number of offspring produced by the parent. This means we wish to find the *n*_*c*_ that maximizes *n*_*c*_*s*_*c*_(*R/n*_*c*_). But this is the exact problem that the original Smith-Fretwell model solves. The optimal resources *r*\* to give each offspring should satisfy

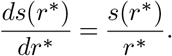

An easy way to find this *r*\* is to graph *s*(*r*) and find the unique line that intersects both the origin and a *single* point of the curved region of *s*(*r*). The *r* coordinate of the point where the line intersects the curve is *r*\*.

Once we know *r*\* we can calculate that *n*_*c*_ = *R/r*\*. If *n*_*c*_ is a non-integer value then we compare the two closest integers and choose the one that yields highest fitness (Charnov et al., 1995).

It may at first seem we have just repeated the original result of Smith and Fretwell. But because of the generality of our setup we have shown that even with the flexibility of distributing unequally, or if *n* is fixed, or if all of *R* must be used, the Smith-Fretwell strategy remains optimal. The algorithm for finding this optimum is

1. Find the *r*\* that satisfies *ds*(*r*\*)*/dr*\* = *s*(*r*\*)*/r*\*.
2. If *r*\* is not an integer, evaluate the fitness for the two nearest integers and select the one which yields highest fitness.

The form of the optimum is

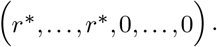

#### B. Concave-decreasing

Concave-decreasing functions (and decreasing functions in general) model detrimental proteins or damaged cellular components (Moger-Reischer and Lennon, 2019). They could also model toxins that multicellular animals pass onto their offspring Klanjscek et al., 2007; Noonburg et al., 2010; Paula et al., 2014, or heavy metals that accumulate in fish eggs (Jezierska et al., 2009) and cross the placental barrier in mammals (Caserta et al., 2013).

Because resources never benefit the offspring in this case, if the parent has the option to withhold resources then any strategy that distributes no more than *r*_*low*_ resources to each offspring is optimal.

If the parent instead *must* distribute all its resources then the situation is more complex. To find an optimum we will transform this problem into one of finding a concave-increasing optimum – a problem that we know how to solve from the previous section. This transformation consists of mirroring *s* across the vertical line at *r*_*max*_ to obtain

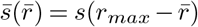

which is a concave-increasing function. The function 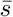 takes in how many resources an offspring is away from *r*_*max*_ and returns its survival probability. If we previously measured survival in units of food a chick received we now measure it in units of food that the chick *didn’t* receive, relative to the maximum.

We have recast the system in units of “anti-resource,” which we denote as 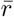. This means 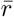 anti-resources corresponds to 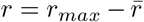 resources and distributing *R* resources now corresponds to distributing 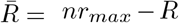 anti-resources. The expected number of surviving offspring is the sum 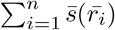.

Now that our optimization problem is in concave-increasing form we can apply the results from the previous section: there exists an optimal anti-resource amount 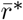 and the parent should produce as many offspring of this size as possible. An optimal strategy is then:

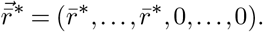

If we use 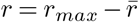 to convert anti-resources back into resources we get

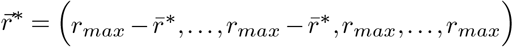

as a solution. The algorithm for finding this optimum is then

1. Convert the problem into one of anti-resources.
2. Find the anti-resource optimum by treating this as a concave-increasing problem.
  - Begin all offspring with zero anti-resources.
  - Find the Smith-Fretwell optimum anti-resource amount 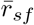, either by finding the tangent point or by solving 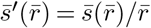.
  - Sequentially provision as many offspring as possible with 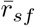 anti-resources.
  - Distribute any remaining anti-resources equally among the offspring with 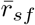 anti-resources. Call this final offspring size 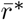.
3. Convert the optimum back into resources.

#### C. Convex-increasing

In the convex-increasing case, resources never harm offspring. This means, like the concave-increasing case, at least one optimal strategy uses all *R* resources. If *R > nr*_*high*_ then any strategy that gives all offspring *r*_*high*_ or more resources is optimal. Otherwise we can show that the optimal strategy is of the form

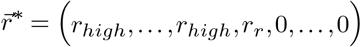

where 0 ≤ *r*_*r*_ *< r*_*high*_.

Consider an optimal strategy. We know by our analysis of convex functions that the offspring on the curved region must be either of size *r*_*low*_ or *r*_*high*_, with an exception of at most one offspring of some intermediate value *r*_*r*_.

Suppose such an intermediate-sized offspring exists. Then there can be no offspring with 0 *< r* ≤ *r*_*low*_. This is because if such an offspring existed we could take resources away from it (which would not reduce fitness) and give those resources to the *r*_*r*_ sized offspring (which would increase fitness). This would give a total increase in fitness and contradict the assumption that the initial strategy was optimal. For the same reason no offspring can have more than *r*_*high*_ resources if there is an offspring of size *r*_*r*_.

Thus, if an intermediate size offspring exists the optimal strategy must have the form:

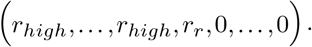

If there is no intermediate offspring then all offspring are either of size less than *r*_*low*_ or of size greater than *r*_*high*_. These are optimal strategies for convex-increasing survival functions. The following algorithm will bring us to one of these optima.

1. Initialize all offspring with zero resources.
2. Increase as many offspring to size *r*_*high*_ as possible.
3. There now remain 0 ≤ *r*_*r*_ *< r*_*high*_ resources. Do one of the following.
  - If all offspring are now of size *r*_*high*_ then we have reached maximum fitness and it does not matter how we distribute the remaining resources.
  - Otherwise allocate the remaining *r*_*r*_ resources to a single offspring.

Under the classic OOS interpretation, *r*_*low*_ is zero and those offspring are never born. Then, if the *r*_*r*_ offspring receives significantly less resources, we would see multiple large offspring and a single small runt.

#### D. Convex-decreasing

In the convex-decreasing case resources never benefit offspring. This can be the case if resources are old components, and is possible in animals under certain conditions concerning size-dependent mortality (Jørgensen et al., 2011).

If the parent may withhold resources it should produce as many offspring as possible and give each no more than *r*_*low*_ resources. This is the optimal strategy as it ensures each offspring has the highest possible survivorship.

Suppose, however, that the parent must distribute all *R* resources and that giving each offspring *r*_*low*_ resources still leaves some resources remaining. To find the optimal strategy we mimic the procedure used for concave-decreasing functions. First, we transform the problem into one of anti-resources. The problem becomes one of optimizing a concave-increasing function where we must use exactly 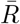 anti-resources. By our previous result for the concave-increasing case, we know the optimum is of the form

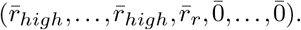

Observing that 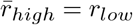 and 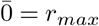 we convert from anti-resources to resources and reorder the terms to find the optimum is

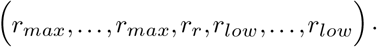

Where 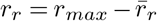. The following algorithm yields this optima.

1. Give every offspring *r*_*low*_ resources. This uses up as many resources as possible without decreasing fitness.
2. If resources remain then sequentially bring as many offspring as possible up to size *r*_*max*_, or as close as possible. All these offspring should end up on the second flat region.
3. If after this resources still remain, then allocate all those remaining resources into one of the *r*_*low*_ offspring. It is now of size *r*_*r*_.

#### E. Concave-increasing-decreasing

In the concave-increasing-decreasing case resources are beneficial or neutral up until a certain point, after which they begin decreasing survival. Let us call this threshold value, after which resources harm the offspring, *r*_*top*_. Functions of this type have been considered in (Marshall et al., 2008; Hendry et al., 2001; Hendry and Day, 2003).

Given the option the parent should never give an offspring more than *r*_*top*_ resources, as this would expend resources to decrease fitness. If the parent can withhold resources, this is straightforward. Even if the parent must spend its resources, as long as *R* ≤ *nr*_*top*_ it is still possible. In either of these cases *s* effectively becomes a concave-increasing function as the parent never produces an offspring that lands beyond the increasing region. The optimal strategy is then the same as in the concave-increasing case; *r*_*top*_ takes the place of *r*_*max*_.

This does not hold when the parent must use up all *R* resources, *and R* is high enough that it must give at least one offspring more than *r*_*top*_ resources. In this case the optimal strategy takes the form

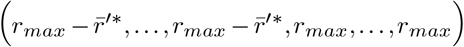

where 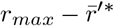 serves as the optimal offspring size and the offspring of size *r*_*max*_ absorbs extra resources. The value of 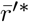 can be calculated as described below in the algorithm for finding this optimal strategy.

1. If *R* ≤ *nr*_*top*_ then set *r*_*max*_ to *r*_*top*_. This restricts the domain of *s* to between zero and *r*_*top*_. Survivorship is now concave-increasing, so find the optimum as one normally would for the concave-increasing case.
2. Otherwise give every offspring *r*_*top*_ resources. This leaves *R*^′^ = *R* − *nr*_*top*_ remaining resources to distribute.
3. Define the function *s*_*d*_ that takes in how many resources *r*^′^ above *r*_*top*_ an offspring is and returns its survivorship.
4. Notice *s*_*d*_ is a concave-decreasing function. Calculate how to optimally distribute the *R*^′^ remaining resources under *s*_*d*_ by following the algorithm given in the concave-decreasing section. Call this strategy 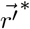.
5. Add *r*_*top*_ to each entry in 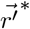.This is an optimal strategy.

The proof that this algorithm gives a optimal strategy can be found in Supplement C.

#### F. Convex-decreasing-increasing

This case describes some resource that is beneficial at low and high levels but detrimental when given in moderation. It could also describe environments that favor specialization (e.g., small offspring can hide, large offspring can fight, and medium offspring can do neither.)

When all *R* resources must be used, and if the optimal strategy contains an offspring with size strictly between *r*_*low*_ and *r*_*high*_, then the optimal strategy is the same as that for either the convex-increasing or the convex-decreasing case. But which of the two it is depends on the exact survivorship function being used.

Heuristically, the convex-decreasing strategy works best when the first flat region is higher, and the convex-increasing strategy works best when the second flat region is higher. But this is only a heuristic, and counterexamples exist (Supplement D.2).

When the parent need not use all *R* (i.e., can withhold resources) and the first flat region is higher or equal to the second flat region then the optimal strategy is to place all offspring in the first flat region. Otherwise the form of the optimum is not easy to classify, and depends on the exact survivorship function.

Supplement D contains the proof of these statements, along with some discussion about the cases where the optimum does not contain a runt. Finding a straightforward algorithmic solution to this case remains an open problem. Unlike the other cases, this one seems to depend on the specific values of the survivorship function.

### Supplement B. Details for constant environments

#### A. Symmetric strategies evolve under concave survivorship functions

In the main paper we showed that the symmetric strategy (having all curved region offspring of the same size) is an optimal strategy. Here we show that mutations towards this optimum are always neutral or beneficial (non-detrimental), implying that the fitness landscape is a simple hill and populations evolve to this optimum.

Our proof strategy is to first consider pairwise mutations: mutations that change resources between two offspring but leave the rest unaffected. We show that these pairwise mutations are non-detrimental only when they make the pair of offspring closer in size. We then show that a mutation in a straight line towards the symmetric strategy can be constructed from a sequence of non-detrimental pairwise mutations, and thus itself must be non-detrimental.

##### Lemma 1.

Suppose that an organism produces *n* offspring under survival function *s*. The curved region of this function, *s*_*c*_, is concave, and at least two of these offspring fall on this region.

Then a pairwise mutation that transfers resources from one curved region offspring to another is either neutral or beneficial if and only if the mutation causes the two offspring to end up closer in size. If *s*_*c*_ is strictly concave then such a mutation is always beneficial.

*Proof:* Suppose that the strategy is not perfectly symmetric. Then at least two offspring on the curved region do not have the same amount of resources. Call these offspring one and two, and without loss of generality, assume that offspring one has fewer resources than offspring two. We write this as *r*_1_ *< r*_2_.

Consider a mutation that takes *ϵ* resources away from offspring two and allocates them to offspring one.

Here *ϵ* is small enough that this action does not move either offspring out of the curved region. We will compare the fitness of the original strategy to that of this new mutant strategy where the resource allocation to offspring one and two is altered.

**Case 1:** Suppose that the mutation does not make offspring one larger than offspring two. The mutant strategy’s fitness is then

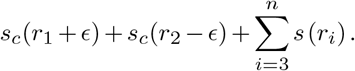

We put in the subscript *c* to emphasize that these offspring remain on the curved region of *s*. Because offspring three and above remain the same between the original and mutant strategies, their contribution to the fitness change can be ignored. The mutant strategy fitness is higher than the original strategy’s when

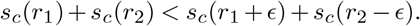

If we let *ϵ* go to zero then this inequality is true if and only if the directional derivative of the fitness *w* increases in the direction of 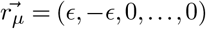.This directional derivative can be written as

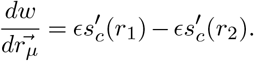

and is either zero or positive when

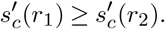

Because *s*_*c*_ is strictly concave, then 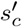 must be monotonically decreasing. This means our assumption that *r*_1_ is less than *r*_2_ implies the above inequality. If *s*_*c*_ is not strictly concave, then equality could be possible, in which case the small mutation would be neutral.

As the derivative in the direction of the mutation will always be non-negative we know that the full mutation for the original non-infinitesimal *ϵ* will be non-detrimental.

**Case 2:** Say the mutation transfers so many resources that offspring one ends up larger than offspring two. That is, *r*_1_ *< r*_2_, but *r*_1_ + *ϵ > r*_2_ − *ϵ*. Such a mutation would be equivalent to switching the offspring indices and transferring a smaller number of resources. This places us back in case one. This is illustrated in S1.

**Case 3:** Suppose a mutation causes the two offspring to have exactly equal resources. In this case (*r*_1_ +*ϵ*) equals (*r*_2_ − *ϵ*), which equals 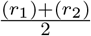. Jensen’s inequality then tells us that this too is either neutral or an improvement.

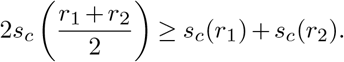

**Figure S1.**
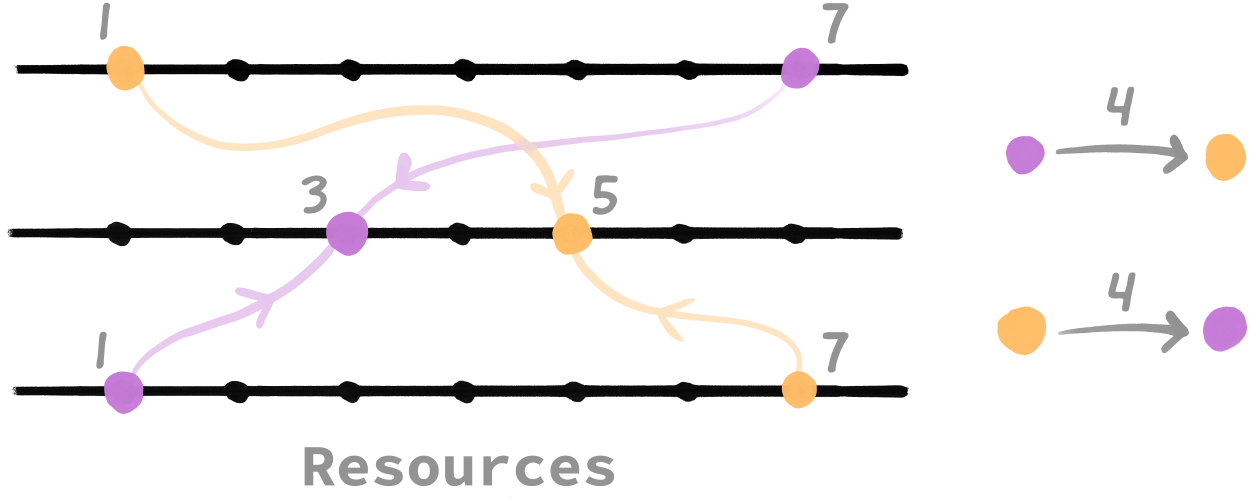
Illustration with two offspring, orange and purple. The position on the line measures of how many resources an offspring has. The transition from the top line to the middle line illustrates a mutation where the purple offspring gives the orange offspring four resources. This causes the purple offspring to become the smaller offspring as in Case 2. Compare this to an equivalent transition from the bottom line to the middle line, which illustrates a mutation where the orange offspring gives four resources to the purple one. This keeps the orange offspring larger than the purple one, placing this transition in case 1. Since a Case 1 transition is non-detrimental, the equivalent Case 2 situation is also non-detrimental.

To summarize, we have considered pairwise mutations that take two offspring and redistribute resources between them. As long as the mutation makes the offspring’s resource values closer together then that mutation is non-detrimental. In the case that *s*_*c*_ is strictly concave, the mutation is always beneficial. On the other hand, pairwise mutations that move the strategy away from symmetry are detrimental if *s*_*c*_ is strictly concave, or neutral/detrimental if *s*_*c*_ is non-strictly concave.

Now let us consider mutations that affect more than a pair of offspring at a time. We can construct any such mutation by composing multiple pairwise mutations together. To see this, consider all possible strategies and realize that this space is spanned by the displacement vector of all pairwise mutations. A quick induction can also yield this result (the lemma proves the base case, for the induction step take the first *n* − 1 elements of the strategy and adjust them such that a pairwise mutation between the first and *n*th element gives the target vector).

In particular this means that we can reach the symmetric strategy by mutating towards it through a series of small pairwise mutations that approximates the path of a straight line. Each of these mutations, if small enough, can be guaranteed to be beneficial or neutral. In the limit the mutation is a straight line towards the symmetric strategy. Doing this from all possible strategies implies that the fitness landscape is a simple “Mt. Fuji-style” one with a peak at the symmetric strategy.

We have purposefully left the definition of “closer to the symmetric strategy” loose. Given an arbitrary mutant strategy 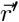 it is difficult to say whether it is detrimental compared to the original strategy 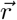. If there exists a pathway of only pairwise non-detrimental/beneficial mutations then the mutant strategy is clearly non-detrimental/beneficial. But if there is only a pathway that makes some detrimental mutations but also some beneficial ones then the picture is less clear as detriments could outweigh benefits or vica versa.

Note that we are assuming mutation operates on the resource distribution vector, and that this vector is what is genetically encoded. We could instead assume that the distribution algorithm is what evolves. This is interesting because then it predicts that the optimal strategy still evolves even if the total resources *R* or maximum number of offspring changes over time, which could be due to evolution, environmental changes, or the ability to accumulate resources increasing as the parent ages. From one perspective this would be plasticity in the number and size of offspring. From another, it would be a non-plastic biological algorithm.

#### B. Asymmetric strategies are optimal under convex survivorship, when restricted to the curved region

Consider a survival function *s* that has a convex curved region. Then an optimal strategy for offspring *restricted to the curved region* is of the form

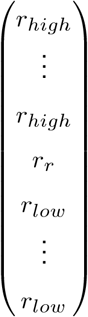

where there is at most one *r*_*r*_ such that *r*_*low*_ *< r*_*r*_ *< r*_*high*_. Furthermore, if the curved region of *s* is strictly convex then this strategy is the strict and unique optimum (up to reorderings of the offspring). Call this optimal strategy 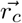. THe subscript *c* indicates that this is the optimum when offspring are restricted to the curved region.

Notice that we do not say how many *r*_*high*_ or *r*_*low*_ there are. This can depend on the exact function. All we claim is that there is some amount (possibly zero) of these, one or zero *r*_*r*_’s, and nothing else.

*Proof:* Proceed by contradiction. Let 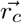 be the optimal strategy of the form given above. Assume there is an alternative strategy 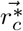 of a different form but that does at least as good as 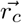 while the same amount of resources. We will let 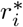 denote the *i*th element in the alternative strategy and *r*_*i*_ the *i*th element in the original. We will say that two such elements with the same index *correspond* with each other.

Since 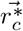 and 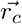 are different at least one of the elements between the two strategies must also be different. Without loss of generality, let *r*_1_ and 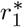 be these differing elements. Say that the difference between these elements has magnitude *ϵ* so that 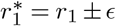. Because *r*_1_ is either *r*_*high*_, *r*_*r*_, or *r*_*low*_ we must be one of the four following cases.

**Table 5.** Possible combinations of 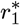 and 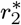. Notice that *r*_1_ is always less than *r*_2_.

| | $r_1^* = r_{low} + \epsilon$ | $r_1^* = r_r + \epsilon$ |
| --- | --- | --- |
| $r_2^* = r_{high} - \delta$ | Possible | Possible |
| $r_2^* = r_r - \delta$ | Possible | Impossible |

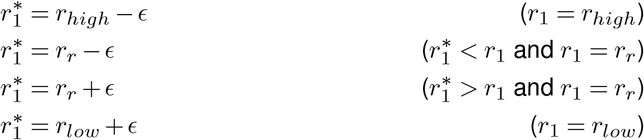

The cases 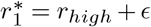 and 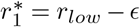 are not possible because we have restricted offspring to the curved region.

Focus on the case where *ϵ* resources have been *added* (the logic for the subtraction case is similar). The extra resources in 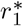 must have come from somewhere, as the total resources used in both strategies are the same. Without loss of generality, say that 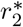 contributed some (or even all) of these resources. This offspring must have *δ* ≤ *ϵ* fewer resources than its corresponding element in 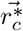. This means *r*_2_ cannot be of value *r*_*low*_, because then 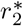 would be outside the curved region. We thus deduce 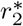 must correspond to either *r*_*high*_ or *r*_*r*_.

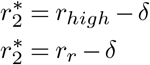

Recall that 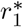 can correspond to either *r*_*r*_ or *r*_*low*_.

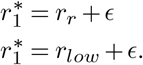

There are three possible combinations of values for 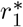 and 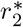. These are shown in the following table. Note that both cannot correspond to *r*_*r*_ at the same time because there is only one *r*_*r*_, so the lower right combination is impossible.

For each possible combination, 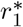 and 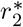 must be closer together than their corresponding entries in the original strategy. For example, taking the upper left possibility we have that the difference between the two entries is

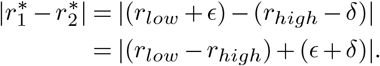

Here the (*r*_*low*_ − *r*_*high*_) gives the difference of the two entries in the original strategy and is negative. Meanwhile the (*ϵ*+*δ*) term is positive. But neither *ϵ* or *δ* can be more than |*r*_*low*_ −*r*_*high*_|, else 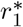 would be out of bounds. At most we can have *ϵ*+*δ* = 2|*r*_*low*_ −*r*_*high*_|. But this is impossible as it would imply the alternative strategy 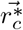 is a swap with 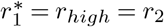 and 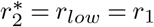, meaning that 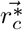 is just a reordering of 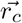 and thus not a different strategy.

Therefore *ϵ*+*δ <* 2 |*r*_*low*_ − *r*_*high*_| and when we compare the original and alternative strategies we can then say

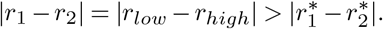

A similar calculation shows that all remaining cases move the two offspring closer together. As convex regions select for moving offspring further apart we expect we can then improve the alternative strategy 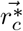 by moving offspring further apart.

Specifically we can move offspring one and two further apart. Consider a new strategy 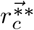, which is the same as 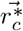 except

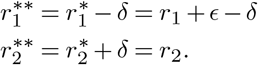

By shuffling resources only between offspring one and two we have made their sizes more like the original strategy, which we know is more spread out than those in 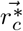.

This action of moving resources between two offspring is the pairwise mutation we considered in lemma 1. We can make a corollary to this lemma for when the survival function is convex instead of concave.

##### Lemma 2.

Suppose that an organism makes *n* offspring under survival function *s*. The curved region of this function, *s*_*c*_, is convex, and at least two of these offspring fall on this region.

Then a pairwise mutation that transfers resources from one curved region offspring to another is either neutral or beneficial if and only if the mutation causes the two offspring to be further apart in size. If *s*_*c*_ is strictly convex then such a mutation is always beneficial.

*Proof:* This is the same proof as for lemma 1, except we let the strategy (*r*_1_ + *ϵ, r*_2_ − *ϵ, r*_3_, …, *r*_*n*_)^*T*^ be the original strategy and (*r*_1_, *r*_2_, *r*_3_, …, *r*_*n*_)^*T*^ be the mutant strategy. Since the curved region *s*_*c*_ is now convex we know 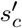 is monotonically increasing and we must also swap the appropriate inequalities.

By this lemma a pairwise mutation that moves offspring further apart increases or maintains fitness. Strategy 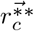 then must have higher fitness than strategy 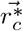. But this contradicts the assumption that 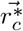 is an optimal strategy.

#### C. Optima under concave-increasing-decreasing survivorship

Say *s* is a survival function with a concave-increasing-decreasing curved region, and say the maximum of the curved region occurs at *r*_*top*_ resources. (In the case where the curved-region is not strict, then let *r*_*top*_ be the lowest resource value that maximizes survival.) If the parent must give at least one offspring more than *r*_*top*_ resources (that is, the parent may not withhold resources and *R > nr*_*top*_) then the optimal strategy is of the form

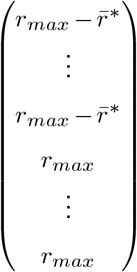

where 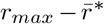 is acting as the optimal offspring size and the offspring of size *r*_*max*_ are used to absorb the extra resources.

Before we begin the proof of this we must show the following lemma.

##### Lemma 3.

If the parent must give at least one offspring more than *r*_*top*_ resources, then it must give all offspring more than *r*_*top*_ resources in the optimal strategy.

*Proof:*

Proceed by contradiction. Say that 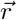 is an optimal strategy where one offspring is of size less than *r*_*top*_. Call this the *small offspring*. We know there is at least one offspring with size larger than *r*_*top*_. Call any such offspring a *large offspring*. Let *r*_*s*_ be the size of the small offspring and *r*_*j*_ be the size of the *j*th large offspring (of which there may be more than one). We can write these resource amount in terms of how far each is from *r*_*top*_.

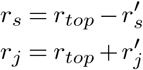

Here 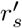 and 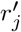 give the distance from *r*_*top*_. Now, sum the 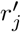 of all the large offspring. This sum must be greater than 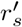 because *R > nr*_*top*_. This means we can take 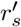 worth of resources from the collection of large offspring and give them to the small offspring, in such a way that each large offspring never goes below *r*_*top*_ resources. Call this new strategy 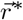.

This process does not lower the survival of any of the large offspring. But, because *r*_*top*_ is the lowest value where *s* reaches its maximum, this process must increase the survival of the small offspring. Thus 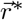has higher fitness than 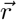,contradicting that 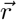 is an optimal strategy.

With this lemma we can prove the main result.

*Proof:* Observe that *s*(*r*_*top*_) is the maximum possible survivorship per offspring. This means a parent who produces *n* offspring and has no resource constraints has a maximum fitness of *ns*(*r*_*top*_). Call this strategy, 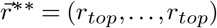, the *ideal strategy*.

Sadly for our parent, the ideal strategy is an impossible strategy, as by assumption the parent must distribute all resources and *R* is high enough that at least one offspring must be larger than *r*_*top*_. The fitness of an optimal strategy then must be less than or equal to the fitness of the ideal strategy. However, the parent can seek to minimize the difference between in total survivorship from this ideal. Call this difference Δ*w* and see that

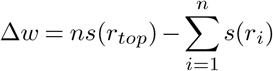

where the *r*_*i*_ are the offspring sizes in the parent’s strategy. Finding an optimal strategy is equivalent to finding a strategy that minimizes Δ*w*.

Because at least one offspring must be larger than *r*_*top*_ our lemma tells us that in an optimal strategy all offspring must be larger than *r*_*top*_. This allows us to rewrite the above equation in terms of how far above *r*_*top*_ each offspring is.

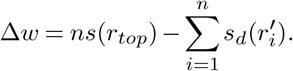

Here 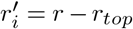 measures how far beyond *r*_*top*_ each offspring is. Call these the *remaining-resources*. There are *R*_*r*_ = *R* − *nr*_*top*_ total remaining-resources. The function *s*_*d*_ gives the survival probability of these offspring in terms of 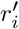. That is, 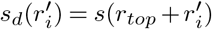.

We can now minimize Δ*w* by maximizing the summation term. This is equivalent to the concave-decreasing problem, which we already know how to solve from Section B. Following that algorithm we transform the problem into one of anti-resources. That is, we wish to maximize 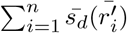 under the constraint that 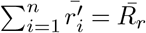 where the over-bar signifies anti-resources as explained in Section B. We find that the optimal distribution of remaining-anti-resources is

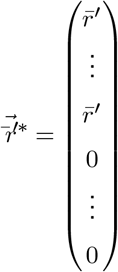

where 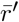 is the optimal anti-resource amount found by optimizing 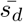. Each offspring can have at most *r*_*max*_ − *r*_*top*_ remaining-resources, so if we transform this strategy from units of remaining-anti-resources to units of remaining-resources we have that the optimal distribution of remaining-resources is

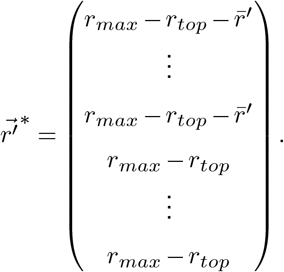

Since this strategy is written in terms of remaining-resources we must add *r*_*top*_ to each entry to get the overall optimal strategy, which is

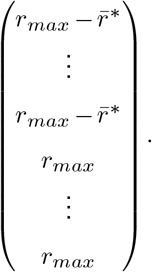

#### D. Optima under convex-decreasing-increasing survivorship

When there is a runt, the optimal strategy for the convex-decreasing-increasing survivorship case is either the optimal strategy for the convex-increasing survivorship case or the optimal strategy for the convex-decreasing survivorship case. To show this we first require a lemma.

##### Lemma 4.

At the optimal strategy, if a runt exists (an offspring with *r*_*low*_ *< r*_*r*_ *< r*_*high*_), then all offspring on the flat region offspring must either all be at 0 and *r*_*high*_ (Option 1) or at *r*_*low*_ and *r*_*max*_ (Option 2).

*Proof:* Given that we have a runt at least one other offspring must be on one of the flat regions (since we have previously proved that for convex cases there is at most one runt.)

Depending on the size of the runt it needs to either lose or gain resources to increase its fitness. Say it needs to increase its resources. If the flat region offspring are not all either at 0 or *r*_*high*_ (Option 1 configuration) then there exists at least one flat region offspring not at 0 or *r*_*high*_. The runt can take a small amount of resources from this offspring to increase its own fitness. This will not decrease the flat region offspring’s fitness as long as it remains on flat region. This increases overall fitness, contradicting that this was an optimal strategy.

The same argument works for the case where the runt requires less resources to increase its fitness. The flat region offspring must all be at *r*_*low*_ or *r*_*high*_ (Option 2 configuration), else the runt could give its resources to one of the flat region offspring without decreasing its own fitness (and potentially increasing total fitness if the recipient moves to a higher flat region or if *r*_*r*_ moves to a better position).

By our lemma we know there are only two possible forms for the optimum (when a runt exists):

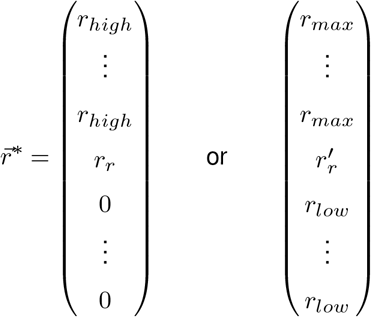

where *r*_*r*_ and 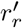 are the respective runt sizes. These forms must be unique in terms of the number of first and second flat region offspring for a given total *R*. For the first one, creating this form in a way that uses up all *R* is equivalent to dividing out *r*_*high*_ from *R* to get the number of size *r*_*high*_ and taking the remainder to be the runt. This is exactly the same as the convex-increasing algorithm. If we convert to anti-resources the argument works for the second optimum form, showing that this form for a given *R*.

**Table 6.** The specific conditions for when following the convex-decreasing strategy gives higher fitness than following the convex-increasing strategy.

| $s(r_r)$ | $s(r'_r)$ | Condition for when convex-decreasing strategy is better |
| --- | --- | --- |
| $s(r_r)$ | $s(r'_r)$ | $s(r_r) - s(r'_r) < \Delta n(\gamma_{low} - \gamma_{high})$ |
| $s(r_r)$ | $\gamma_{low}$ | $s(r_r) < \Delta n(\gamma_{low} - \gamma_{high}) + \gamma_{low}$ |
| $s(r_r)$ | $\gamma_{high}$ | $s(r_r) < \Delta n(\gamma_{low} - \gamma_{high}) + \gamma_{high}$ |
| $\gamma_{low}$ | $s(r'_r)$ | $-s(r'_r) < \Delta n(\gamma_{low} - \gamma_{high}) - \gamma_{low}$ |
| $\gamma_{high}$ | $s(r'_r)$ | $-s(r'_r) < \Delta n(\gamma_{low} - \gamma_{high}) - \gamma_{high}$ |
| $\gamma_{low}$ | $\gamma_{low}$ | $0 < \Delta n(\gamma_{low} - \gamma_{high})$ |
| $\gamma_{low}$ | $\gamma_{high}$ | $0 < (\Delta n - 1)(\gamma_{low} - \gamma_{high})$ |
| $\gamma_{high}$ | $\gamma_{low}$ | $0 < (\Delta n + 1)(\gamma_{low} - \gamma_{high})$ |
| $\gamma_{high}$ | $\gamma_{high}$ | $0 < \Delta n(\gamma_{low} - \gamma_{high})$ |

##### D.1. A heuristic for deciding which strategy performs better

A natural conjecture is that the increasing strategy will work when the first flat region is higher than the second, and vica versa. This is an okay heuristic, but is not true as per the convex-decreasing-increasing counterexample (next subsection).

We can derive a basic condition though. Say we run the increasing algorithm. Let *n*_*l*_ be the number of offspring on the first flat region and let *n*_*h*_ be the number in the second flat region. Let *γ*_*low*_ and *γ*_*high*_ be the fitnesses of those regions respectively. Let *r*_*r*_ be the remainder allocated to the runt. Then the fitness of the increasing strategy is

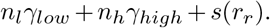

Now consider running the decreasing algorithm. Say that the algorithm produces Δ*n* more offspring on the lower region (i.e., *r*_*low*_ or 0) and correspondingly Δ*n* fewer offspring on the higher region (i.e., *r*_*max*_ or *r*_*high*_). It should always be more or equal as the convex-decreasing survivorship algorithm generates more small offspring. (We have not proved this statement, but it is intuitively true and a large simulation agrees.)

The fitness for the decreasing strategy is then

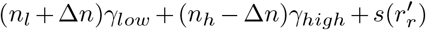

where 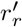 is the size of the runt in the decreasing strategy. Then a basic inequality calculates that the fitness of the decreasing strategy is higher if and only if

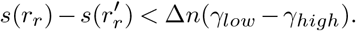

It is possible that the runts are in one of the flat regions so we would still need to consider that.

Temporarily, let us call the very last offspring that the algorithms move the runt. This way the chosen runt is unique and there is always a runt, even it is are technically in the flat regions. This means we can keep the above inequality and simply look at all the combinations of the runt’s fitness and how the inequality simplifies These can tell us about specific cases. But note that we have not proved that the increasing or decreasing strategy are optima for the runt-on-flat-region cases. This table only compares the two to say which is better. There is no guarantee of the choice being the best.

##### D.2. Counterexample to heuristic

This is a counterexample to the heuristic that a higher first flat region implies we should use the convex-increasing survivorship optimum and a higher second flat region implies we should use the convex-decreasing survivorship optimum.

Take a convex-decreasing-increasing survivorship function such that *n* = 4, *R* = 8, *r*_*l*_ = 2, *r*_*h*_ = 5, *r*_*m*_ = 8, and *s*(3) = 0.2. The optimal strategy from applying the convex-increasing algorithm is (5, 3, 0, 0). If we instead apply the convex-decreasing algorithm the optimal strategy is (2, 2, 2, 2).

Next, imagine two situations. Situation A has the left side of the survivorship function higher with *γ*_*low*_ = 1 and *γ*_*high*_ = .9. Situation B has the right side higher with the numbers swapped. We can calculate the fitnesses for both in the following table.

| Situation | Strategy | Fitness |
| --- | --- | --- |
| B | Convex-increasing | 3 |
| B | Convex-decreasing | 3.6 |
| A | Convex-increasing | 3.1 |
| A | Convex-decreasing | 4 |

We find that following the convex-decreasing strategy is always better in these specific examples.

#### E. If the symmetric strategy is best then the asymmetric strategy is worst, and vice-versa

Imagine *s* is a concave function. We know the optimal strategy is symmetric. Instead of measuring offspring in terms of survival probability, let us measure them in terms of death probability. Call this function *d* and see that

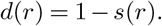

The strategy that minimizes the expected number of surviving offspring is the same strategy that maximizes the expected number of dying offspring. But *d* is a convex function with two flat regions; we already have shown in the main paper that the optimum here is an asymmetric strategy.

Thus we have shown that if *s* is concave, the best possible strategy is symmetric while the worst possible is asymmetric. The argument is similar for when *s* is convex.

#### F. Conjectured strategies for Sigmoidal, Gaussian-like, and other survivorship functions

Below are two conjectures, concerning Gaussian-like and the concave-convex-increasing survivorships. Note that these are conjectures, and have only been numerically checked that they hold against a small handful of “reasonable” functions.

##### F.1. Gaussian-like and Sigmoidal

For sigmoidal or Guassian-like survivorship functions (that is, a convex-increasing region followed by a concave-increasing region followed by a concave-decreasing region followed by a convex-decreasing region) we conjecture another optimization algorithm. We will go through the algorithm for Gaussian-like survivorship functions. The sigmoidal function is special case where we assume that *s*(*r*_*p*_) to *s*(*r*_*max*_) is flat.

Say that the survivorship function has a single maximum point, which we say occurs when we give *r*_*p*_ resources. Then the first phase of the algorithm is to give as many offspring *r*_*p*_ resources as possible. If the parent has enough resources to do this and does not need to distribute them all offspring are on the peak, thus have maximum value, and we are done.

Otherwise if the parent does not have enough resources to do this they will act like an asymmetric distribution strategy and create one runt. The strategy vector will have the form

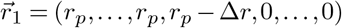

where Δ*r* is how many resources away from the peak the runt offspring is. We have labeled it 1 to denote the first case. Likewise, should we have too many resources to put each offspring at *r*_*p*_ then we can start all offspring at *r*_*p*_ and successively fill each offspring to the maximum number of resources until we run out of resources, since all resources must be distributed. Such a strategy will take the form with a “runt-like” offspring too. We will denote this strategy 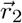.

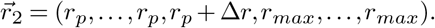

Here now the “runt” is larger than the peak offspring, and Δ*r* measures how much larger.

These are not the final optimized strategies but just the starting points. To further refine our optimization we will take all the offspring with *r*_*p*_ resources and either take or give them all the same *ϵ* amount of resources, and either give or take from the runt to insure the total amount of resources remain the same. Let *k* be the number of offspring with size *r*_*p*_. Then the final optimized strategies are

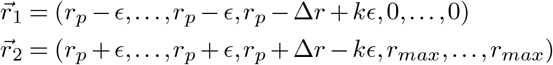

where the exact value of *ϵ* is found by starting it at 0 and continuously increasing it until the total fitness no longer increases. These two equations above are then the conjectured forms of the optimal strategies under Gaussian-like survivorship functions.

Biologically we will have two classes of offspring, small and large, with possibly one offspring in the middle. On the first case the small offspring may not exist and we have many large offspring and one runt as in the convex-increasing case.

Some numerical simulations suggest this “sliding-*ϵ*” technique can be generalized to monotonically increasing then decreasing survivorship functions with an arbitrary finite number of inflection points.

##### F.2. Concave-convex-increasing

One could also ask about sigmoidal and Gaussian-like functions where the convexities are swapped. The convexity-swapped sigmoidal function would then have an increasing concave component followed by an increasing convex component.

For these functions the optimum appears to be to give some number of offspring *r*_*high*_ resources and to distribute the remaining resources equally among the other offspring.

To find the number of offspring with *r*_*high*_ resources is non-trivial. Let *n* be the total number of offspring and *n*^′^ be the number of offspring that receive *r*_*high*_ resources. For every possible *n*^′^ value calculate a corresponding *r*^′^ to be the positive solution to the following equation.

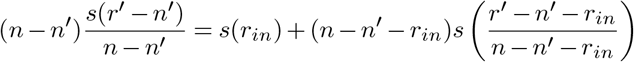

where *r*_*in*_ is the inflection point. Pick the largest of these *r*^′^ which is smaller than *R*. The corresponding *n*^′^ is the number of offspring that receive *r*_*high*_ resources. This latter case could explain situations where parents make some of their offspring large and some small.

#### G. Proof of Smith Fretwell Model

For convenience we provide the standard Smith-Fretwell result and its proof. This section only re-translates results from Smith and Fretwell, 1974 and Charnov et al., 1995 into our terminology. We also provide a proof for completeness.

Say that a parent has *R* resources to equally distribute among *n* offspring so that each offspring receives*R/n* = *r* resources. The survival probability of an offspring given *r* resources is *s*(*r*), where *s* is a concave-increasing survivorship function. A parent’s expected number of surviving offspring, *ns*(*r*), we call the parent’s fitness. Then

1. There exists a unique optimal number of offspring *n*\* and resources-per-offspring *r*\* that maximizes the total fitness, where

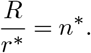

1. On the graph of *s*(*r*) the optimal resources-per-offspring *r*\* corresponds to the point where a line passing through the origin intersects *s*(*r*) at only one point. Say this line has slope *w*\*
2. If a parent makes *n*\* offspring then the parent’s fitness is *Rw*\*.
3. The optimal value of *r*\* satisfies

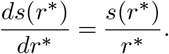

*Proof:* We will derive this with the same method as Smith and Fretwell did.

If the parent makes *n* offspring then each offspring gets *R/n* resources. This means that each offspring has a survival chance of *s*(*R/n*). The fitness of the parent is the expected number of offspring to survive which we can calculate to be *n · s*(*R/n*).

(1 and 2.) We wish to maximize *ns*(*R/n*). This is equivalent to maximizing 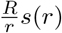, where *r* is the amount of resources given to each offspring. Call this the strategy *r*.

Consider a straight line passing through the origin with slope *w*, as shown in Figure 1. Every point (*x, y*) on this line represents a strategy that gives *x* resources to each offspring so that each offspring has a survival probability of *y*. This gives a total fitness of

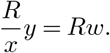

Because *R* is fixed the fitness is controlled entirely by the slope of the line. Points on steeper lines have higher fitness. But not all the points on these lines are possible strategies for our organism. Its possible strategies fall on the curve of *s*(*r*). This means we can find the fitness of the strategy *r* by finding the slope of the line that intersects the point (*r, s*(*r*)) and the origin.

See that the steepest line from the origin that intersects *s*(*r*) is the only one tangent to *s*(*r*). It only touches in one point, which shows that the optimal resource value per offspring *r*\* is unique, and because *r*\* = *R/n*\* we can calculate the optimal number of offspring. Because this is the unique tangent we have that 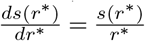. All this shows the last three items.

### Supplement C. Detailed proofs for fluctuating environments

#### A. Optima for fluctuating environments

Here we will prove the results of the fluctuating environments section more formally. Recall that we wish to maximize geometric mean fitness. This can be written both in exponential form and product form.

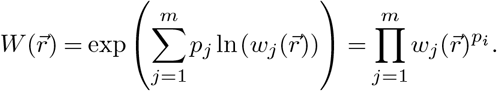

Here *p*_*i*_ is the frequency of the *i*th environment and 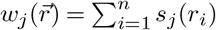 is the fitness of strategy 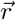 under environment *j*. For all that follows we consider only feasible 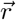, i.e. those that meet our constraints that we can use no more than (or exactly) *R* resources.

In this fitness landscape we have an analogue of the curved region of *s* from the constant environment case: a concave or convex function which we can apply the techniques of convex optimization. But under fluctuating environments there may be many such curved regions, rather than one (see Figure 5).

Our first task is to then to devise a way to refer to regions. This will require a language to describe the *s*_*j*_, as shown below.

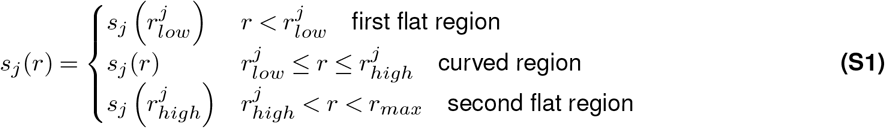

The *j* superscript on *r* is not a power but a indicator of the environment. For example, 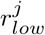 is the lower bound of the curved region of *s*_*j*_. Notice that all environments must share the same *r*_*max*_, as otherwise the geometric mean would be undefined if any offspring was larger than the minimum of the 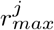.

Observe that the 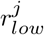 and 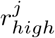 partition the line segment [0, *r*_*max*_]. Because this is equivalent to placing 2*m* points we can have up to 2*m* + 1 segments. As a technical detail will define each segment as open, so that the points 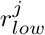 and 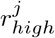 along with the endpoints 0 and *r*_*max*_ are not in any of the segments. For two environments this is shown in the first column of Figure 5.

Enumerate these segments such that the first is 1 and the last is *q*. We can then define an indicator function *f* that takes in a strategy 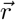 and, for the *i*th offspring, tells us which segment it is in, or if it is in one of boundary points that correspond to no segment.

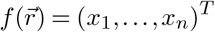

where,

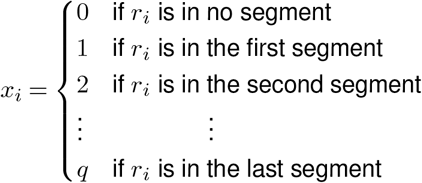

Let 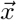 be such a indicator vector. Define the set

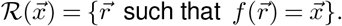

That is, 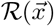 is the set of all resource vectors for which the *r*_*i*_ are restricted to their respective segments defined by 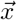. For example, in a three offspring system ℛ (1, 5, 5) gives all strategies such that the first offspring size is restricted to the first segment while the second offspring and third offspring sizes are restricted to the fifth segment. In the two environment case this is shown by the grid in the second column of 5. Each cell of the grid corresponds to a 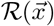.The same idea holds for more than two environments, but the grid is multidimensional.

Why do we do this? Because once a strategy 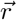 is restricted to a 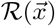 the convexity of 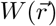 is guaranteed not to change provided that all the curved regions of *s*_*j*_ are concave (we will prove this is true in the next section). This paints a picture of *W* as a partition of *sections*, at most one for each of the *q*^*n*^ possible indicator vectors 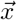. If the curved region of each *s*_*j*_ is concave within the segments defined by 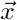, then each of these sections is concave, even if the entirety of *W* is not. There is also a region described by the boundaries of the segments, which is the union of all the 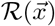 where at least one element of 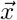 is zero. For two environments this is illustrated in the third column of Figure 5.

To make an analogy, the fitness landscape *W* is a quilt of concave patches, and the boundary region is the stitching between each patch. In the next subsection we prove that this is true.

##### A.1 All the s_j_ are concave

Take the sections and number them arbitrarily and let *W*_*k*_ denote the *k*th section. That is, *W*_*k*_ is the restriction of *W* to 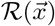, where 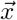 is the *k*th indicator vector. We wish to show the following.

###### Theorem 1.

If the curved region of all the *s*_*j*_ are concave, then each *W*_*k*_ is concave.

Our strategy is to decompose *W*_*k*_ into a composition of concave functions, then take advantage of the conditions where compositions of concave functions are concave. To begin, let *G* be the weighted geometric mean

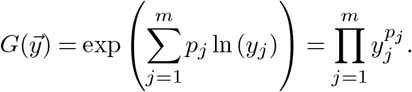

We will consider *G* on non-negative *y*_*j*_, as these will ultimately be the fitness from each environment. *W* can be thought of as the composition of the *w*_*j*_ with *G*. Define the function *w* which outputs a vector of fitnesses for each environment,

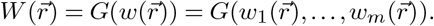

With this setup we can prove that each *W*_*k*_ is concave.

###### Lemma 5.

The weighted geometric mean 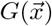, for positive *x*_*i*_, is a non-decreasing function. That is, if we increase any of the *x*_*i*_ then 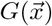 does not decrease.

*Proof:* For positive reals exponentiation by *p*_*i*_ is non-decreasing and multiplication is non-decreasing for positive terms. The composition must also be non-decreasing.

###### Lemma 6.

The weighted geometric mean is concave.

*Proof:* MaoWao, n.d.; Boyd, 2023

###### Lemma 7.

Let *h*: ℝ^*m*^ → ℝand *g*_*i*_ : ℝ^*n*^ → ℝbe twice differentiable functions. Let 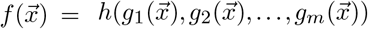. Then *f* is concave if *h* is concave and nondecreasing and all the *g*_*i*_ are concave.

*Proof:* See (Boyd and Vandenberghe, 2004), Chapter 3 for the proof and extensions to situations where *h* and *g*_*i*_ are not differentiable.

Observe that if all the *s*_*j*_ are concave on their respective domains within a section *W*_*k*_, then each 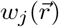 is concave, as sums of concave functions remain concave. By remaining restricted to the domain of *W*_*k*_ we guarantee that the *s*_*j*_ are concave. The *r*_*i*_ in flat regions must remain in flat regions and *r*_*i*_ in curved regions must remain in curved regions. The above lemma then applies as *W*_*k*_ is the composition of the weighted geometric mean with the *w*_*j*_.

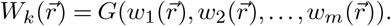

Here the *w*_*j*_ are restricted to the same domain as *W*_*k*_. We know *G* is a non-decreasing concave function by lemmas 6 and 5. We also know the *w*_*i*_ are all concave. Lemma 7 then implies *W*_*k*_ is a concave function.

This tells us every extrema is a local optimum – local to the section it is in. If these are global extrema is a far more difficult question. The Karush-Khun-Tucker conditions are one approach to finding these, as shown in Supplement B.

#### B. Global optima in fluctuating environments case

Finding the optimal strategy for fluctuating environments is difficult. Luckily the Karush-Kuhn-Tucker conditions (KKT conditions) hint at what points could be optima. The KKT conditions generalize the Lagrange multiplier method for constrained optimization. Importantly for us, they check the border conditions and allow constraints to be inequalities (Hillier and Lieberman, 2010).

To apply these conditions we must restrict ourselves to the special case when all *s*_*i*_ share the same *r*_*low*_. This means that all offspring need the same minimal investment before resources affect survival chance, regardless of the environment. This allows us to ignore the first flat region and focus on the curved region, ensuring the function is everywhere differentiable.

Since we have only one constraint (that resources sum to *R*) the KKT conditions simplify and tell us that, for 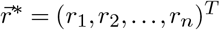 to be a potential global optimum of *w*, there must exist a constant *u* that meets the following six conditions.

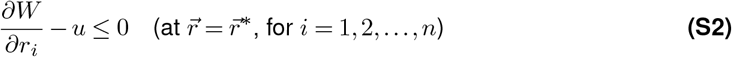

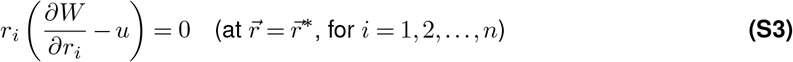

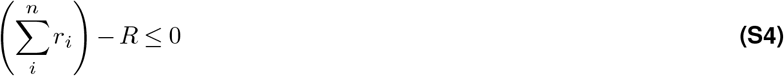

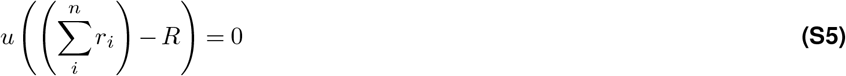

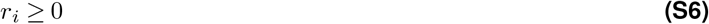

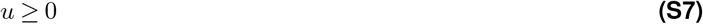

There are further requirements for our constraint 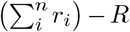, but as it is an affine function we can ignore these.

Condition 3 encodes our resource constraint, and condition 5 encodes that offspring cannot receive negative resources. For conditions 1 and 2 we can calculate the partial derivative *∂W/∂r*_*i*_. This is straightforward if you write the geometric mean in product form and apply a generalized product rule.

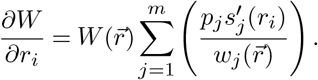

The denominator term 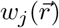 measures the expected survival of strategy 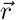 in environment *j*. We can assume that this is not zero, because if for some environment it was zero then there would be an environment where no offspring survive and the geometric mean fitness 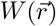 would be zero, putting us in the trivial case where 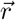 is a worst strategy.

Notice that condition 1 and 2 together say that at *r*_*i*_ or 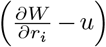 must be zero. This is equivalent to saying

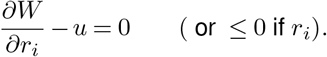

Let us say that the first *k* of the *n* offspring are non-zero-sized and the rest are zero. Then condition 1 and 2 tell us that an optimal strategy is one which solves the following system of equations while keeping with conditions 3 to 6.

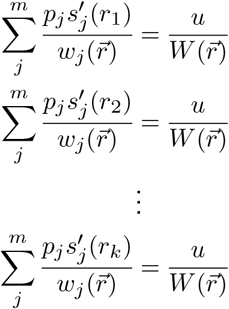

(We have rearranged and divided out by 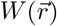 as we assume it is non-zero). Condition 3 and 4 tell us that either *u* is zero or all resources are used. In the former case resources are left over and the equations simplify to

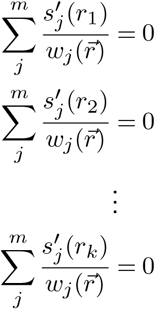

Otherwise *u* is non-zero. But as the KKT requirement is that some positive *u* exists we can rewrite it as 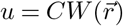 for some positive *C*. This allows us to simplify and say that a strategy 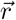 is optimal if there exists a positive *C* that solves the following system.

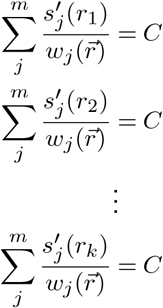

For both cases the constraint that 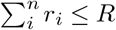 is implies that the set of feasible solutions is convex. This guarantees that all optima of this system are maxima (when the *s*_*i*_ are all concave) or minima (when the *s*_*i*_ are all convex).

This system is difficult to solve in general, but some special cases are possible. In symmetric case where the *k* offspring all get equal amounts of resources all the *r*_*i*_ are equal and thus each equation trivially sums to the same amount, meaning the symmetric strategy is always a global extrema provided the sum is positive.

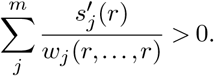

This gives a test which allows one to check if a given symmetric strategy is a global optimum for arbitrary survivorship functions *s* in fluctuating environments.

##### B.1. All the s_j_ are convex

One might naively expect we can mimic the above proof to show the *W*_*k*_ are all convex if the *s*_*j*_ are convex. But this does not work, as we would require that the geometric mean to be a non-*increasing* function. We can in fact find examples where *W*_*k*_ is not convex.

That said, in numerical simulations the *W*_*k*_ are often convex or almost convex (which is to say, it looks convex to the naked eye). Discovering the conditions which guarantee convexity of *W*_*k*_ we leave as an open problem.

### Supplement D. Stage-Structure

Adding age or stage-structure can often modify a model’s predictions (Caswell, 2006). However, our predictions still follow those of tables 2 and 3, provided that the parent has the flexibility to decide on a different distribution strategy for each stage.

Consider a stage-structure defined by the following Leslie matrix, under a constant environment.

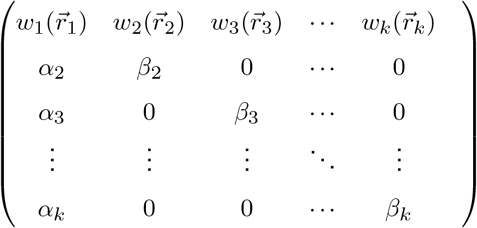

Subscript index the life-stages. Here *α*_*i*_ and *β*_*i*_ are the probabilities of respectively surviving to or staying in the *i*th stage. The 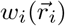 captures the expected number of surviving offspring from a stage *i* individual adopting strategy 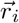. The fitness *w*_*i*_ is subscripted because each stage may have a different fitness landscape, may have a different amount of available resources, or for physiological reasons may have resources affect offspring differently.

We must be careful about what “surviving offspring” means here; they are offspring which survive to the first reproductive stage, stage one. There is implicitly a zeroth juvenile stage through which the offspring immediately pass, and only 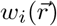 of them survive to stage one. Alternatively one could explicitly adds this juvenile stage and change the interpretation of “surviving offspring” to “offspring that survive the process of birth.”

The dominant eigenvalue of this matrix will give the long-term population growth rate. Because each stage’s strategy 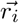 is independent of 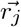 for *j* ≠ *i*, it is best to adopt the optimal strategy for each stage. This maximizes each 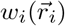 element in the matrix, and it is well known that increasing an individual non-negative element of a non-negative irreducible matrix does not decrease the lead eigenvalue (Berman and Plemmons, 1994). The parent’s overall strategy is then simple: look at the life-stage it is in, the amount of resources it has, and the fitness landscape *w*_*i*_ of its life-stage it is in. Then perform the optimal strategy for that situation as per tables 2 and 3. Furthermore, such a strategy can evolve by small mutations as each of the 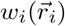 can evolve the optimal strategy, and the evolution of each of the 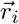 are independent.

This all of course presupposes that the 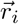 are independent, a reasonable assumption under predictable environments. All the organism requires is a way to measure which life-stage it is in along with the genetic flexibility to encode and trigger a different distribution strategy for each life-stage.

Fluctuating environments complicate this story. Say each environment occurs independently with probability *p*_*j*_. We can imagine that each timestep we select a Leslie matrix *L*_*j*_, corresponding to the current environment, to update the population with.

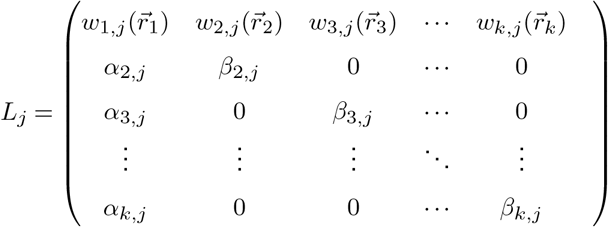

Here the Leslie matrix is the same as in the constant environment case, except the *j* subscript communicate that parameters and fitness landscapes may change across environments. The strategy for each life-stage remain fixed across environments, modeling an organism that can genetically encode its distribution strategy for each life-stage but cannot predict the environment. We cannot easily analyze this as the matrix elements are no longer independent – choosing the best 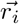 for one environment may reduce fitness in another environment.

We expect the population to evolve the 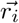 which maximize the average-of-growth-rates. Imagine replicating the evolution of the system many times from a given initial condition. For each replicate we can calculate the growth rate. Take the average of all these growth rates and you find the average-of-growth rates

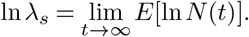

This is the parallel to geometric mean fitness for stage-structure in that it is what natural selection optimizes (Caswell, 2006) since almost all populations grow at the rate ln *λ*. Exact calculation of ln *λ* is difficult but approximations exist (Cohen, 1977; Tuljapurkar, 1982a; Tuljapurkar, 1982b; Caswell, 2006; Tuljapurkar, 2013). If we calculate the related average-population-size growth rate we discover that each 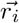 independently evolves over a landscape similar to those found in fluctuating environments without stage-structure (Supplement D). It would be a nice parallel if the maximization of the average-of-growth-rates is equivalent to independently maximizing the geometric mean fitness of each life-stage, but whether this is true we leave as an open question.

#### A. Open Problems

All our stage-structure results assume that each life-stage is independent of the others. What happens when an organism must choose a single strategy for all life-stages? If all the survivorship functions share convexity do we still get the qualitative outcome that concave implies symmetry and convex implies asymmetry? What is the most general biological system where we find this outcome?

Another key problem is to find the optimal strategy when the survivorship function is sigmoidal, an alternative realistic scenario (Brockelman, 1975; Rollinson and Hutchings, 2013). Lin et al. explored Hill-type functions in a bacterial aging model and witnessed a phase-transition from asymmetric to symmetric strategies as protein accumulation rates change (Lin et al., 2019). Our own pilot simulations suggest the optimal strategies tend to have a form similar to an asymmetric strategy (see appendix section F). This hints that protein uptake rates might have a similar role to total resource values. A solution to the sigmoidal case would give us a grip on fluctuating environments that mix concave and convex survivorship functions, as sections of these landscapes can appear as multidimensional sigmoidal functions.

We can extend the sigmoidal problem to the problem of Gaussian survivorship, and the even more ambitious problem of when the survivorship function is made out of a finite number of concave and convex sections. An answer to the latter would let us approximate the optimum to almost all naturally occurring survivorship functions. While problems of this sort are easy to state that does not guarantee they are easy to solve; the general resource distribution problem under arbitrary survivorship functions is equivalent to the NP-hard unbounded knapsack problem (Martello and Toth, 1990; Bretthauer and Shetty, 2002).

No matter how flexible we make the survivorship function our model assumes that all offspring share it. In reality, offspring may have different genotypes. Or sibling competition may create differences in offspring success. A fetus’s placement in the womb, and egg’s placement in a clutch, or a seed’s placement in the plant can all grant different survivorship functions to each offspring (Telfer and Rutberg, 1960; McLaren and Michie, 1960; Harper et al., 1970; McKeown et al., 1976; Roach and Wulff, 1987; Brown and Shine, 2009).

If we give each offspring its own survival function the theory of convex optimization lets us say some mathematical truths. (As long as the functions all remain concave then their sum will be concave. The set of feasible strategies will be the same as before, and this set is both concave and convex. Then any local maxima is in fact global. If parental fitness is strictly concave, which is true when each survival function is strictly concave, then such a maxima is unique. In the case where the survival functions are convex then the above also holds, expect the maxima are now minima.) Organisms will evolve towards the global optimal strategy or away from a global minima. But we cannot always say what these extrema are and have no deep understanding. We cannot even invoke the Purkiss principle if the functions are non-identical.

Just as offspring are not identical neither are resources. Plants require both carbon and nitrogen for their seeds and both elements affect survival in a concave-increasing manner (McGinley and Charnov, 1988). In general there may be multiple resources, each with their own effect on the survivorship function. Such situations are another open problem. We expect the results to remain similar if the (now multidimensional) survivorship function remains concave or convex-increasing. But the situation quickly becomes complicated with different resources affecting survivorship with different kinds of convexity. We might expect that survivorship becomes a multidimensional sigmoidal-like function, again bringing us back to question of optimizing sigmoidal functions.

#### B. Stage-structure in fluctuating environments

There are at least two sensible definitions for population growth rate for matrix models under a stochastic environment. Using the notation of (Caswell, 2006), let *N*(*t*) be a random variable giving the population size at time *t*.

The growth rate discussed in the text is the average-of-growth-rates,

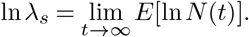

Here we replicate evolution of the system many times from a given initial condition. For each replicate we calculate the growth rate. Take the average of all these growth rates to get ln *λ*_*s*_.

Compare this to the average-population-size growth rate,

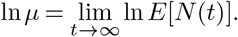

Here at every time *t* we take average population size and measure how fast it grows.

If natural selection maximizes ln *µ*, then we have an answer similar to our constant environment case. Provided that environments are independently distributed, the average-population-size growth rate is equal to the dominant eigenvalue of 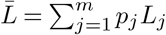, the weighted average of matrices. In particular, the elements of the first row are weighted sums. For example, the first element of the first row is

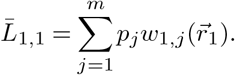

Say that each *w*_1,*j*_ is concave; that is, all of the survivorship functions that make them up are concave and the convexity does not change with environments. Then the entire weighted sum must be concave, as weighted sums preserve convexity. Or, more accurately, the sum will correspond to a fitness landscape made of concave sections, patched together under the same logic as in the non-stage-structured fluctuating environments case (Section A.2). Unlike the non-stage-structured case, if all the survivorship functions are convex the entire sum is *guaranteed* to be made up of convex functions.

Repeating this logic shows that each element in the top row of 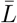 can be written as a function that features a patchwork fitness landscape similar to the non-stage-structured fluctuating environments case. As in the constant environment stage-structured case, the 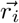 are independent, so we can find the vector which maximizes the weighted sum for each stage.

However, as discussed previously, natural selection likely maximizes the much harder to calculate ln *λ*_*s*_ rather than ln *µ*.

