## Supplementary material for "Resource distribution unifies optimal offspring size and bacterial aging": Figure code: Figures.ipynb

````
# %% [markdown]
# Figures

Here is the code to make the figures for *Unifying optimal offspring size and bacterial aging through resource distribution models, Sakal and Proulx 2023*. 

First we will import the needed packages. Makie will be our plotting software with CairoMakie as the backend. We will also use Algebra Of Graphics theme as the base. We will use the turtle resource distribution figure as inspiration for our color scheme.

- Survivorship Function: A thick orange-red line. If there are multiple survivorship functions they will be the same color but different patterns.
- Region Seperators and other things: Degrees of a blue green shade.
- Maxima: A blue color.

# %% [markdown]
##  Setup

First we set up the default theme and some often used colors and colorshemes. Sometimes we will modify the theme for a certain plot. You can rerun this code to return to the default theme.

# %% [code]
using CairoMakie, AlgebraOfGraphics, GLMakie, LaTeXStrings, ColorSchemes
set_aog_theme!() 


CairoMakie.activate!(type = "svg")

update_theme!(                  # Tweaks the current theme
           fontsize=25,
           markersize=30,
           resolution = (2000, 2000)

       )

#update_theme!(fonts = (; regular = "FiraSans-Medium"))

survivorship_color = "#ff6012"
maxima_marker = :dot
maxima_color = "#0072b2"# :red

num_colors = 6 
color_gradient = ColorSchemes.BuGn_6[1:num_colors]
color_cycle = [color_gradient[i] for i in 1:num_colors]

# %% [markdown]
## Figure 1: Smith Fretwell curves.

This figure gives the standard Smith Fretwell survivorship curve, its derivative, and a dotted line tangent to the survivorship curve which intersects at the maxima of the derivative. This is a standard figure in the literature. 

# %% [code]
CairoMakie.activate!(type = "svg")
update_theme!(               
           resolution = (1800, 600),
       )

# A classic Smith Fretwell survivorship function
function SF_survivorship(r)
	max(1 - exp(-1 * (r-2)), 0)
end

# Fitness when there are twelve resources and they are shared equally amongst R/n offspring.
function SF_fitness(r)
    R = 8  # Total number of resources
    max(R/r * SF_survivorship(r), 0)
end

f = Figure()  # Setup figure
xmax = 7  # How far to plot
lw = 6  # Line width

# First Plot
ax1 = Axis(
    f[1,1],
    title = "Survivorship of single offspring per resource",
    xlabel = "r (resources)",
    ylabel = "s(r) (survival probability)",
    limits = (nothing, nothing, nothing, 1.5),
    xticks = ([3.5], [L"r^*"]),
    yticks = [0, 1],
    aspect = 2,
    xticklabelsize = 30,
    yticklabelsize = 30,
)

lines!(
    0:.1:xmax, 
    SF_survivorship,
    linewidth = 6,
    color = survivorship_color
    )

# Make straight line of constant fitness.
lines!(
    0:.1:xmax, 
    Array(0:.1:xmax) * .221963,  # slope calculated manually
    linestyle = :dash,
    linewidth = lw
)

scatter!(3.5, SF_survivorship(3.5), color=maxima_color, marker=maxima_marker, markersize = 25)  # Put dot at intersection point

# Second Plot
ax2 = Axis(
    f[1,2],
    title = "Parental Fitness",
    xlabel = "Resources per offspring",
    ylabel = "Expected number of surviving offspring",
    xticks = ([3.5], [L"r^*"]),
    aspect = 2,
    xticklabelsize = 30,
    yticklabelsize = 30,
)

lines!(
    0:.1:10, 
    SF_fitness,
    linewidth = 6,
    color = color_cycle[5]
)

scatter!(3.5, SF_fitness(3.5), color=maxima_color, marker=maxima_marker, markersize = 25)  # Put dot at intersection point


f


# %% [code]
save("Smith-Fretwell Curve.pdf", f)  # Save the figure

# %% [markdown]
## Figure 2: Turtles and distribution strategies

The turtles depicted in this figure were created using Midjourney AI then modified in Photoshop. We thank and acknowledge the numerous artists whose artwork contributed to the training of the AI, despite not having granted explicit permission or received compensation. 

# %% [markdown]
## Figure 3: Anatomy of the survivorship function

This figure shows the parts of the survivorship function. 

# %% [code]

# Function for the curved region
function anatomy_curved(r; n=.7, k=.1, a=.7, b=.6)
    ((-r+b)^n)/(k + (-r+b)^n) - ((-r+b)^2)/2 - (-r+b)
end

# The same function, but with constant regions
function anatomy(r)
	if r < 0.2
        return anatomy_curved(.2)
    elseif r >.55
        return anatomy_curved(.55)
    else
        return anatomy_curved(r)
    end

end


# %% [code]
update_theme!(               
           resolution = (1200, 600),
       )
f = Figure()  # Setup figure
xmax = .7  # How far to plot
lw = 6  # Line width


# First Plot
ax1 = Axis(
    f[1,1],
    xlabel = "r (resources)",
    ylabel = "s(r) (survival probability)",
    aspect = 2,
    limits = (.05, .71, .3, .66),
    xticks = ([0.2, 0.55, .7], [ L"r_{low}", L"r_{high}", L"r_{max}"]),  # Custom tick locations for x axis
    yticks = ([anatomy(0), anatomy(1) ], [L"\gamma_{low}", L"\gamma_{high}"]),
    xticklabelsize = 32,
    yticklabelsize = 32,
)

# regions
fontsize = 25

band_color1 = color_cycle[5]
band!([0,.2], [.3, .3], [.66, .66], color = (band_color1, .1))
text!(ax1, "First Flat Region", position = (0.125, 0.63), align = (:center, :center), color = color_cycle[6], fontsize = fontsize, font = :bold)

band_color2 = color_cycle[2]
band!([.2, .55], [.3, .3], [.66, .66],color = (band_color2, .1))
text!(ax1, "Curved region", position = (0.375, 0.63), align = (:center, :center), color = :grey2, fontsize = fontsize, font = :bold)

band_color3 = color_cycle[5]
band!([.55, .7], [.3, .3], [.66, .66], color = (band_color3, .1))
text!(ax1, "Second flat region", position = (0.625, 0.63), align = (:center, :center), color = color_cycle[6], fontsize = fontsize, font = :bold)

vlines_color = (:green, .1)
vlines!(.2, linestyle = :dash, color = vlines_color, linewidth = 3)
vlines!(.55, linestyle = :dash, color = vlines_color, linewidth =3)

lines_color = :lightgrey
lines!([0, .7], [anatomy(1)], linewidth =1, color = lines_color)
lines!([0, .7], [anatomy(0)], linewidth = 1, color = lines_color)

lines!(0:.001:xmax, anatomy, linewidth = lw, color = survivorship_color)
f


# %% [code]
save("Anatomy.pdf", f)  # Save the figure

# %% [markdown]
# Figure 4: Six survivorship functions

This creates a table of the six survivorship functions we consider. 

# %% [code]
# Survivorship function
function s(r, rlow, rhigh, curve_function)
    if r < rlow
        return curve_function(rlow)
    elseif r > rhigh
        return curve_function(rhigh)
    else
        return curve_function(r)
    end
end

# ===============
# Various Base Functions
# For making these functions we use the fact that linear combinations
# of concave (convex) functions are concave (convex)
# ===============

# Default function params
RLOW = 2
RHIGH = 10
RMAX = 12
D = 0.8
GLOW = 0
GHIGH = .95

# ----------------------------------
_curve_concave_increasing_1(r; d=1) = 1 - exp(-d * (r - RLOW))
_curve_concave_increasing_2(r; d=2) = (1 - 1.4^(-d * (r - RLOW))) * 0.7
_curve_concave_increasing_3(r) = _curve_concave_increasing_2(r, d=.7) * 0.7 + _curve_concave_increasing_1(r, d=3) * 0.5 + .1

# --------------------------------------
_curve_concave_decreasing_1(r, rmax=RMAX; d=1) = _curve_concave_increasing_1(rmax - r - 2, d=d)
_curve_concave_decreasing_2(r, rlow=RLOW, rmax=RMAX) = (1 - 1.4^(-1.3 * (rmax - r - rlow))) * .9 - .1
_curve_concave_decreasing_3(r) = (_curve_concave_decreasing_2(r) + _curve_concave_decreasing_1(r, d=5)) * 0.5

# -----------------------------------
_curve_convex_increasing_1(r) = 1 - _curve_concave_decreasing_1(r)
_curve_convex_increasing_2(r) = 1 - _curve_concave_decreasing_2(r)
_curve_convex_increasing_3(r) = 1 - _curve_concave_decreasing_3(r)

# ----------------------------------------------------
_curve_convex_decreasing_1(r) = 1 - _curve_concave_increasing_1(r)
_curve_convex_decreasing_2(r) = 1 - _curve_concave_increasing_2(r)
_curve_convex_decreasing_3(r) = 1 - _curve_concave_increasing_3(r)

# --------------------------------------
_curve_concave_both_1(r) = (.6 * _curve_concave_increasing_1(r, d=2) + 2 * _curve_concave_decreasing_1(r, d=.6)) * .3
_curve_concave_both_2(r) = _curve_concave_increasing_1(.8 * (r + 0.5), d=.9) * .8 * _curve_concave_decreasing_1(r)
_curve_concave_both_3(r) = 0.7 * ((_curve_concave_increasing_1(.8 * (r + 2), d=1) * .8 * _curve_concave_decreasing_1(r)) + (
        _curve_concave_increasing_1(.8 * r, d=1) * .8 * _curve_concave_decreasing_1(r)))

# ---------------------------------------
_curve_convex_both_1Here is the continuation of the translation for the Python code into Julia:

```julia
_curve_convex_both_1(r) = 1 - _curve_concave_both_1(r)
_curve_convex_both_2(r) = 1 - _curve_concave_both_2(r)
_curve_convex_both_3(r) = 1 - _curve_concave_both_3(r)


# %% [code]
CairoMakie.activate!(type = "svg")

rlow = 3
rhigh = 15
rmax = 18
D = 0.8
GLOW = 0
GHIGH = .95

# Define the functions
f1(r) = (1 - 1.4^(-1 * (r - rlow))) * 0.7
f2(r) = (1 - 1.4^(-1.3 * (rmax - r - rlow))) * .9 - .1
f3(r) = -(r-8)^2
f4(r) = - f1(r)
f5(r) = - f2(r)
f6(r) = -f3(r)

# Create a Figure
fig = Figure(resolution = (1000, 800))

# Define a function to generate and plot an Axis
function plotfunc(ax, f)
    x = 2
    overlap = .01
    lines!(ax, 0:0.01:rlow, repeat([f(rlow)], length(0:0.01:rlow)), linewidth = 5, color = color_cycle[5])  # first flat region
    lines!(ax, rhigh:0.01:rmax, repeat([f(rhigh)], length(rhigh:0.01:rmax)), linewidth = 5, color = color_cycle[5])  # second flat region
    lines!(ax, rlow-overlap:0.01:rhigh+overlap, f.(rlow-overlap:0.01:rhigh+overlap), linewidth = 5, color = survivorship_color)  # curved region
    scatter!(rlow, f(rlow-overlap), color = survivorship_color, markersize = 6)
    scatter!(rhigh, f(rhigh+overlap), color = survivorship_color, markersize = 6)

    ax.xticklabelsvisible = false
    ax.yticklabelsvisible = false
    ax.xticksvisible = false
    ax.yticksvisible = false
end


# Generate the text
axtop = Axis(fig[1,2], xticklabelsvisible = false, yticklabelsvisible = false, xticksvisible = false, yticksvisible = false, title="Concave", height = Relative(-1))  
hidespines!(axtop)
hidedecorations!(axtop)

axtop = Axis(fig[1,3], xticklabelsvisible = false, yticklabelsvisible = false, xticksvisible = false, yticksvisible = false, title="Convex", height = Relative(-1))  
hidespines!(axtop)
hidedecorations!(axtop)

axtop = Axis(fig[2,1], xticklabelsvisible = false, yticklabelsvisible = false, xticksvisible = false, yticksvisible = false, title="Increasing", height = Relative(0))  
hidespines!(axtop)
hidedecorations!(axtop)

axtop = Axis(fig[3,1], xticklabelsvisible = false, yticklabelsvisible = false, xticksvisible = false, yticksvisible = false, title="Decreasing", height = Relative(0))  
hidespines!(axtop)
hidedecorations!(axtop)

axtop = Axis(fig[4,1], xticklabelsvisible = false, yticklabelsvisible = false, xticksvisible = false, yticksvisible = false, title="Both", height = Relative(0))  
hidespines!(axtop)
hidedecorations!(axtop)

# Generate the Axes and plot the functions
ax1 = Axis(fig[2, 2]); plotfunc(ax1, f1)
ax2 = Axis(fig[3, 2]); plotfunc(ax2, f2)
ax3 = Axis(fig[4, 2]); plotfunc(ax3, f3)
ax4 = Axis(fig[2, 3]); plotfunc(ax4, f4)
ax5 = Axis(fig[3, 3]); plotfunc(ax5, f5)
ax6 = Axis(fig[4, 3]); plotfunc(ax6, f6)


fig


# %% [markdown]
To finish this figure and  get the text alignment and cropping correct we edited it in Photoshop. 

# %% [code]
save("SixFunctions.pdf", fig)  # Save the figure

# %% [markdown]
# Figure: Geometric Mean Fitness


# %% [markdown]
Below is the code for the figure itself. It is the most complex of figures, and requires many hacky workarounds to get looking correct. 

CairoMakie fails to give the proper transparency effects, so we use GLMakie instead. However, this means we cannot save the figure in vector form and instead must save it as a very high resolution png. However,  Makie appears to have a resolution cap of ~4k? To get the resolution high enough we cut the figure into three parts and glue them together in Photoshop. In Photoshop we also fix minor visual bugs and add the tick labels to the rightmost column, as there appears to be a bug which prevents latex labels to appear in 3d axes.

There is also a bug where sometimes the scale of the survivorship functions is wrong. I don't know why this happens but rerunning the same code multiple times seems to fix it.

First we define the survivorship and geometric mean fitness functions.

# %% [code]
# Define the functions we will use.

rlow = 2
rhigh  = 10
rmax  =  15

# Survivorship in environment 1
function s1(r)
    d = 4
    return max(0.5 * (1 - exp(-d * (r - rlow)) - r * 1/8), 0)
end

# Survivorship in environment 2
function s2(r)
    d = 1
    return max((1 - 1.4^(-d * (r - 2 - rlow))) * 0.7 - 1/16 * (r - 2), 0)
end

# This function is extra and we don't use it.
function s3(r)
    d=1
    return max(1.2 * (1 - exp(-d * (r - 4 - rlow))) - (r - 4) * 1/6, 0)
end


# Geometric mean fitness of strategy r1 and r2, with frequency p1 of environment 1
function W(p1, r1, r2 )
        return exp(p1 * log(s1(r1) + s1(r2)) + (1-p1) * log(s2(r1) + s2(r2)) )    
end

# Plot the survival function, but with flat regions added
function add_flat_regions(s, r, rlow, rhigh)

    if r < rlow
        return s(rlow)
    elseif r > rhigh
        return s(rhigh)
    else
        return s(r)
    end

end


# %% [markdown]
Next we set up some plot options.

# %% [code]
GLMakie.activate!(inline=true)  # We will use GLMakie because it allows finer control of transparency through overdraw 


update_theme!(                  # Tweaks the current theme
           fontsize=80,
           markersize=100,
           resolution = (4500, 1600),

       )

rlow = 2 
rmax = 22

ticksize = 90  # Size of tick text
markersize= 60  # Size of dot markers


xmax = 18  # How far to plot
lw = 12  # Line width
top = .4  # how far the bands reach
wirewidth = 6  # width of the wireframe mesh. This is thick enough that it appears as a solid surface
wirespace = .05 
surfacealpha = .3  # transparecy of grey background surface

gridoffset = .00
gridcolor = :white
gridwidth = 8  # width of the gridlines in the rightmost plot
gridsymbol = :solid  # style of the gridlines
gridtransparecy = true  # whether the grid is transparent
gridoverdrawtransparency = true  # whether the overdraw grid is transparent
gridalpha = .8  # alpha of the 
gridwidthflat = 6  # Size of the gridlines for the middle column plots

symwidth1 =  7 # Width of the nontransparent blue symmetry line. Can't be too big because it hase a white outline
symwidth2 =  10  # Width of the transparent blue symmetry line plotted over the other one. 


# %% [markdown]
### Part 1

# %% [code]

# ===========TWO CONCAVE, SAME RLOW AND RHIGH ====================================================

# ------------- Define the Functions ------------------------------------------

f = Figure()  # Setup figure

function s1_curve(r)
    d=4
    shift = -1.5
    x = .4
    return max(0.5 * (1 - exp(-d * (r*x - shift - rlow)) - (r*x-shift) * 1/8), 0) 
end

function s2_curve(r)
    d = 1
    r = rmax-r  # Mirror the function
    shift = 1.3
    x = .6
    out =max((1 - 1.4^(-d * (r*x - shift - rlow))) * 0.7 - 1/16 * (r*x - shift) , -.075) + .075
end


# Set the parameters
rlow = 2.1
rhigh = 14

# Add the flat regions into the survivorship functions
s1(r) = add_flat_regions(s1_curve, r, rlow, rhigh)
s2(r) = add_flat_regions(s2_curve, r, rlow, rhigh)

# ------------ Survivorship Functions ---------------------------------------------------

ax1 = Axis(
    f[1,1],
    xlabel = "r",
    ylabel = "s₁(r) and s₂(r) ",
    aspect = 1,
    limits = (0, xmax, 0, top),
    xticks = ([rlow, rhigh, xmax],   [L"r_{low}", L"r_{high}",  L"r_{max}"]),  # Have to shift the rlow2 tick a bit. Don't know why.,  
    yticks = [0],
    xticklabelsize = ticksize,
    yticklabelsize = ticksize,

)
  
# Seperate the regions by bands
alpha = .3  # How faint the bands are
band!(
    [0, rlow],
    [0, 0],
    [top , top],
    color = (color_cycle[1], alpha),
)
band!(
    [rlow, rhigh],
    [0, 0],
    [top , top],
    color = ( color_cycle[3], alpha),
)
band!(
    [rhigh, xmax],
    [0, 0],
    [top, top],
    color = ( color_cycle[5], alpha),
)

# Boundry Lines
vlines!(rlow, linestyle = gridsymbol, color = (gridcolor, 1), linewidth = gridwidthflat)
vlines!(rhigh, linestyle = gridsymbol, color = (gridcolor, .6), linewidth = gridwidthflat)

# Plot the survivorship functions (put the fainter line behind for a sense of depth)
lines!(
    0:.01:xmax, 
    s2,
    linewidth = lw,
    color = Cycled(2)  # An orange color in the Algebra of Graphics theme
    )
lines!(
    0:.001:xmax, 
    s1,
    linewidth = lw,
    color = survivorship_color
    )
    
# --------------------- The Grid of regions ---------------------------------------------

ax2 = Axis(
    f[1,2],
    xlabel = "r₁",
    ylabel = "r₂",
    aspect = 1,
    limits = (0, xmax, 0, xmax),
    xticks =  ([rlow, rhigh, xmax],   [L"r_{low}", L"r_{high}",  L"r_{max}"]),
    yticks =  ([rlow, rhigh, xmax],   [L"r_{low}", L"r_{high}",  L"r_{max}"]),
    xticklabelsize = ticksize,
    yticklabelsize = ticksize,
     
)

# Region for surface 
# X-axis
alpha = .5
top = xmax
band!(
    [0, rlow],
    [0, 0],
    [top , top],
    color = (color_cycle[1], alpha),
    transparency = true
)
band!(
    [rlow, rhigh],
    [0, 0],
    [top , top],
    color = ( color_cycle[3], alpha),
    transparency = true
)
band!(
    [rhigh, xmax],
    [0, 0],
    [top, top],
    color = ( color_cycle[5], alpha),
    transparency = true
)

# Y axis
band!(
    [0, xmax],
    [rlow, rlow],
    [0 , 0],
    color = (color_cycle[1], alpha),
    transparency = true
)

band!(
    [0, xmax],
    [rhigh, rhigh],
    [rlow, rlow],
    color = (color_cycle[3], alpha),
    transparency = true
)
band!(
    [0, xmax],
    [xmax, xmax],
    [rhigh, rhigh],
    color = (color_cycle[5], alpha),
    transparency = true
)

# Gridlines 

vlines!(rlow, linestyle = gridsymbol, color = gridcolor, linewidth = gridwidthflat, transparency = true)
vlines!(rhigh, linestyle = gridsymbol, color = gridcolor, linewidth = gridwidthflat, transparency = true)
hlines!(rlow, linestyle = gridsymbol, color = gridcolor, linewidth = gridwidthflat, transparency = true)
hlines!(rhigh, linestyle = gridsymbol, color = gridcolor, linewidth = gridwidthflat, transparency = true)

# ---------- Fitness Landscape ---------------------------------------------------

R = 25
# I can't get Latex to work with the tick lables for the 3D plot. So we will add those in after with photoshop
ax3 = Axis3(
    f[1,3 ],
    azimuth =  -π/2  -.2,
    elevation = .9,
    xlabel = "r₁",
    ylabel = "r₂",
    zlabel = "W(r₁, r₂)",
    #aspect = (1,1,1),
    xticklabelsvisible = false,
    yticklabelsvisible = false,
    zticklabelsvisible = false,
    zticksvisible = false,
    xticks = [rlow, rhigh, xmax],  
    yticks = [rlow, rhigh, xmax],   
    xticklabelsize = ticksize,
    yticklabelsize = ticksize,
)

# Make the surface. Enumerate through each region of the middle plot
regions = [0, rlow,  rhigh, xmax]
for (i, r1) in enumerate(regions)
    for (j, r2) in enumerate(regions)
        space = wirespace
        if i < 4 && j < 4
            r1s = regions[i]:space:regions[i+1]  # x axis
            r2s = regions[j]:space:regions[j+1] # y axis
            zs = [W(1/2,  r1,  r2) for r1 in r1s, r2 in r2s]  # z axis

            #Plot the three of these
           
            wireframe!(
                    r1s, 
                    r2s, 
                    zs,
                    color = (color_cycle[i*2-1], .5),
                    transparency = true,
                    linewidth = wirewidth
                    )
                wireframe!(
                r1s, 
                r2s, 
                zs,
                color = (color_cycle[j*2-1], .5),
                linewidth = wirewidth,
                transparency = true,
                )
                surface!(
                    r1s, 
                    r2s, 
                    zs,
                    colormap = (:greys, surfacealpha),
                    )
    
        end
    end
end


# Plot the section boundries

r1_values = 0:0.001:xmax
r2_values = repeat([rlow], length(r1_values))
z_values = [W(1/2, rlow, r)+gridoffset for r in r1_values]
lines!(ax3, r1_values, r2_values, z_values, linewidth = gridwidth, linestyle = gridsymbol, color = (gridcolor, .5), overdraw=false, transparency = gridtransparecy)
lines!(ax3, r1_values, r2_values, z_values, linewidth = gridwidth, linestyle = gridsymbol, color = (gridcolor, .2), overdraw=true, transparency = gridoverdrawtransparency)

r1_values = 0:0.001:xmax
r2_values = repeat([rhigh], length(r1_values))
z_values = [W(1/2, rhigh, r)+gridoffset for r in r1_values]
lines!(ax3, r1_values, r2_values, z_values, linewidth = gridwidth, linestyle = gridsymbol, color = (gridcolor, .5), overdraw=false, transparency = gridtransparecy)
lines!(ax3, r1_values, r2_values, z_values, linewidth = gridwidth, linestyle = gridsymbol, color = (gridcolor, .2), overdraw=true, transparency = gridoverdrawtransparency)

r2_values = 0:0.001:xmax
r1_values = repeat([rhigh], length(r2_values))
z_values = [W(1/2, r, rhigh)+gridoffset for r in r2_values]
lines!(ax3, r1_values, r2_values, z_values, linewidth = gridwidth, linestyle = gridsymbol, color = (gridcolor, .5), overdraw=false, transparency = gridtransparecy)
lines!(ax3, r1_values, r2_values, z_values, linewidth = gridwidth, linestyle = gridsymbol, color = (gridcolor, .2), overdraw=true, transparency = gridoverdrawtransparency)

r2_values = 0:0.001:xmax
r1_values = repeat([rlow], length(r2_values))
z_values = [W(1/2, r, rlow)+gridoffset for r in r2_values]
lines!(ax3, r1_values, r2_values, z_values, linewidth = gridwidth, linestyle = gridsymbol, color = (gridcolor, .5), overdraw=false, transparency = gridtransparecy)
lines!(ax3, r1_values, r2_values, z_values, linewidth = gridwidth, linestyle = gridsymbol, color = (gridcolor, .2), overdraw=true, transparency = gridoverdrawtransparency)


# Find the maxima 
max_val = -Inf
max_r1, max_r2 = -1, -1

for r1 in rlow:.1:rhigh
    for r2 in rlow:.1:rhigh
        val = W(1/2, r1, r2)
        if val > max_val
            max_val = val
            max_r1, max_r2 = r1, r2
        end
    end
end

max_val += .004

# Plot the maxima. There will be two because the landscape is symmetric
scatter!(ax3, [max_r1], [max_r2], [max_val], markersize = markersize*.8, color = (maxima_color, .8), overdraw = false, marker = maxima_marker)
scatter!(ax3, [max_r2], [max_r1], [max_val], markersize = markersize*.8, color = (maxima_color, .8), overdraw = false, marker = maxima_marker)
# Replot with transparency and overdraw for asthetics 
scatter!(ax3, [max_r1], [max_r2], [max_val], markersize = markersize, color = (maxima_color, .4), overdraw = false, marker = maxima_marker, transparency = true, ssao = true)
scatter!(ax3, [max_r2], [max_r1], [max_val], markersize = markersize, color = (maxima_color, .4), overdraw = false, marker = maxima_marker, transparency = true, ssao = true)
scatter!(ax3, [max_r1], [max_r2], [max_val], markersize = markersize, color = (maxima_color, .4), overdraw = true, marker = maxima_marker, transparency = true, ssao = true)
scatter!(ax3, [max_r2], [max_r1], [max_val], markersize = markersize, color = (maxima_color, .4), overdraw = true, marker = maxima_marker, transparency = true, ssao = true)

# generate points where r1 = r2 to show symmetric strategies
r_values = 0:0.001:xmax-.1
z_values = [W(1/2, r, r)+.004 for r in r_values]

# plot the line where r1=r2. 
# we need to plot multiple lines with varying values and options to get the transparency to work out correctly
lines!(ax3, r_values, r_values, z_values, linewidth = symwidth1, color = (maxima_color, .2), overdraw=false, ssao = true)
lines!(ax3, r_values, r_values, z_values, linewidth = symwidth1, color = (maxima_color, .2), overdraw=true, ssao = true)
lines!(ax3, r_values, r_values, z_values, linewidth = symwidth2, color = (maxima_color, .4), overdraw=false, transparency =true, ssao = true)
lines!(ax3, r_values, r_values, z_values, linewidth = symwidth2, color = (maxima_color, .6), overdraw=true, transparency=true, ssao = true)


save("Landscapes1.png", f)

f  

# %% [markdown]
### Part 2

# %% [code]


# ===========================================================================
# ===========TWO CONCAVE, DIFFERENT RLOW AND RHIGH==============================
# ===========================================================================


f = Figure()  # Setup figure


function s1_curve(r)
    d=4
    shift = 0
     return max(0.5 * (1 - exp(-d * (r - shift - rlow)) - (r-shift) * 1/8), 0) + .05
end

function s2_curve(r)
    d = 1
    r = rmax-r  # Mirror the function
    shift = 7
    out = max((1 - 1.4^(-d * (r - shift - rlow))) * 0.7 - 1/16 * (r - shift) , 0) + .025
end


# Set the parameters
rlow1 = 2.25
rhigh1 = 8
rlow2 = 4
rhigh2 = 11.55

s1(r) = add_flat_regions(s1_curve, r, rlow1, rhigh1)
s2(r) = add_flat_regions(s2_curve, r, rlow2, rhigh2)


# ------------ Survivorship Functions ---------------------------------------------------
top = .4  # how far the bands reach

ax1 = Axis(
    f[1,1],
    xlabel = "r",
    ylabel = "s₁(r) and s₂(r) ",
    aspect = 1,
    limits = (0, xmax, 0, top),
    xticks = ([rlow1, rhigh1, rlow2  + .45, rhigh2, xmax],   [L"r_{low}", L"r_{high}", L"r'_{low}", L"r'_{high}", L"r_{max}"]),  # Have to shift the rlow2 tick a bit. Don't know why.,  
    yticks = [0],
    xticklabelsize = ticksize,
    yticklabelsize = ticksize,

)
  
# Seperate the regions by bands
alpha = .3  # How faint the bands are
band!(
    [0, rlow1],
    [0, 0],
    [top , top],
    color = (color_cycle[1], alpha),
)
band!(
    [rlow1, rlow2 + .45],
    [0, 0],
    [top , top],
    color = ( color_cycle[2], alpha)
)
band!(
    [rlow2 + .45, rhigh1],
    [0, 0],
    [top, top],
    color = ( color_cycle[3], alpha)
)
band!(
    [rhigh1, rhigh2],
    [0, 0],
    [top, top],
    color = ( color_cycle[4], alpha)
)
band!(
    [rhigh2, xmax],
    [0, 0],
    [top, top],
    color = (color_cycle[5], alpha)
)

# Boundry Lines
# Change the alpha manually so that the lines remain preceptually constant between the regions
vlines!(rlow1, linestyle = gridsymbol, color = (gridcolor, 1), linewidth = gridwidthflat)
vlines!(rhigh1, linestyle = gridsymbol, color = (gridcolor, .6), linewidth = gridwidthflat)
vlines!(rlow2 + .45, linestyle = gridsymbol, color = (gridcolor, .9), linewidth = gridwidthflat)
vlines!(rhigh2, linestyle = gridsymbol, color = (gridcolor, .5), linewidth = gridwidthflat)


# Plot the survivorship functions (put the fainter line behind for a sense of depth)
lines!(
    0:.01:xmax, 
    s2,
    linewidth = lw,
    color = Cycled(2)  # An orange color in the Algebra of Graphics theme
    )
lines!(
    0:.001:xmax, 
    s1,
    linewidth = lw,
    color = survivorship_color
    )
    
# --------------------- The Grid of regions ---------------------------------------------

ax2 = Axis(
    f[1,2],
    xlabel = "r₁",
    ylabel = "r₂",
    aspect = 1,
    limits = (0, xmax, 0, xmax),
    xticks = ([rlow1, rhigh1, rlow2  + .45, rhigh2, xmax],  [L"r_{low}", L"r_{high}", L"r'_{low}", L"r'_{high}", L"r_{max}"]),  
    yticks = ([rlow1, rhigh1, rlow2  + .45, rhigh2, xmax], [L"r_{low}", L"r_{high}", L"r'_{low}", L"r'_{high}", L"r_{max}"]),  
    xticklabelsize = ticksize,
    yticklabelsize = ticksize,
     
)

# Region for surface 
# X-axis
alpha2 = .5
top = xmax
band!(
    [0, rlow1],
    [0, 0],
    [top , top],
    color = (color_cycle[1], alpha2),
    transparency = true
)
band!(
    [rlow1, rlow2 + .45],
    [0, 0],
    [top , top],
    color = ( color_cycle[2], alpha2),
    transparency = true
)
band!(
    [rlow2 + .45, rhigh1],
    [0, 0],
    [top, top],
    color = ( color_cycle[3], alpha2),
    transparency = true
)
band!(
    [rhigh1, rhigh2],
    [0, 0],
    [top, top],
    color = ( color_cycle[4], alpha2),
    transparency = true
)
band!(
    [rhigh2, xmax],
    [0, 0],
    [top, top],
    color = (color_cycle[5], alpha2),
    transparency = true
)

# Y axis
band!(
    [0, xmax],
    [rlow1, rlow1],
    [0 , 0],
    color = (color_cycle[1], alpha2),
    transparency = true
)
band!(
    [0, xmax],
    [rlow2+.45, rlow2+.45],
    [rlow1, rlow1],
    color = (color_cycle[2], alpha2),
    transparency = true
)
band!(
    [0, xmax],
    [rhigh1, rhigh1],
    [rlow2 + .45, rlow2 + .45],
    color = (color_cycle[3], alpha2),
    transparency = true
)
band!(
    [0, xmax],
    [rhigh2, rhigh2],
    [rhigh1, rhigh1],
    color = (color_cycle[4], alpha2),
    transparency = true
)
band!(
    [0, xmax],
    [xmax, xmax],
    [rhigh2, rhigh2],
    color = (color_cycle[5], alpha2),
    transparency = true
)

# Gridlines
vlines!(rlow1, linestyle = gridsymbol, color = gridcolor, linewidth = gridwidthflat, transparency = true)
vlines!(rhigh1, linestyle = gridsymbol, color = gridcolor, linewidth = gridwidthflat, transparency = true)
hlines!(rlow1, linestyle = gridsymbol, color = gridcolor, linewidth = gridwidthflat, transparency = true)
hlines!(rhigh1, linestyle = gridsymbol, color = gridcolor, linewidth = gridwidthflat, transparency = true)
vlines!(rlow2+.45, linestyle = gridsymbol, color = gridcolor, linewidth = gridwidthflat, transparency = true)
vlines!(rhigh2, linestyle = gridsymbol, color = gridcolor, linewidth = gridwidthflat, transparency = true)
hlines!(rlow2+.45, linestyle = gridsymbol, color = gridcolor, linewidth = gridwidthflat, transparency = true)
hlines!(rhigh2, linestyle = gridsymbol, color = gridcolor, linewidth = gridwidthflat, transparency = true)
# ---------- Fitness Landscape ---------------------------------------------------

R = 25
# I can't get Latex to work with the tick lables for the 3D plot. So we will add those in after with photoshop
ax3 = Axis3(
    f[1,3 ],
    azimuth =  -π/2  -.2,
    elevation = .9,
    xlabel = "r₁",
    ylabel = "r₂",
    zlabel = "W(r₁, r₂)",
    #aspect = (1,1,1),
    xticklabelsvisible = false,
    yticklabelsvisible = false,
    zticklabelsvisible = false,
    zticksvisible = false,
    xticks = [rlow1, rhigh1, rlow2  + .45, rhigh2, xmax],  
    yticks = [rlow1, rhigh1, rlow2  + .45, rhigh2, xmax], 
    xticklabelsize = ticksize,
    yticklabelsize = ticksize,
)

# Make the surface
regions = [0, rlow1, rlow2  + .45, rhigh1,  rhigh2, xmax]
for (i, r1) in enumerate(regions)
    for (j, r2) in enumerate(regions)
        space = wirespace
        if i < 6 && j < 6
            r1s = regions[i]:space:regions[i+1]  # x axis
            r2s = regions[j]:space:regions[j+1] # y axis
            zs = [W(1/2,  r1,  r2) for r1 in r1s, r2 in r2s]  # z axis

            # Plot the three of these
            
            wireframe!(
                    r1s, 
                    r2s, 
                    zs,
                    color = (color_cycle[i], .5),
                    linewidth = wirewidth,
                    transparency = true
                    )
            wireframe!(
            r1s, 
            r2s, 
            zs,
            color = (color_cycle[j], .5),
            linewidth = wirewidth,
            transparency = true
            )
            surface!(
                r1s, 
                r2s, 
                zs,
                colormap = (:greys, surfacealpha),
                transparency = false
                )
        end
    end
end


# Plot the section boundries 
r1_values = 0:0.001:xmax
r2_values = repeat([rlow1], length(r1_values))
z_values = [W(1/2, rlow1, r)+gridoffset for r in r1_values]
lines!(ax3, r1_values, r2_values, z_values, linewidth = gridwidth, linestyle = gridsymbol, color = (gridcolor, gridalpha), overdraw=false, transparency = gridtransparecy)
lines!(ax3, r1_values, r2_values, z_values, linewidth = gridwidth, linestyle = gridsymbol, color = (gridcolor, .2), overdraw=true, transparency = gridoverdrawtransparency)

r1_values = 0:0.001:xmax
r2_values = repeat([rhigh1], length(r1_values))
z_values = [W(1/2, rhigh1, r)+gridoffset for r in r1_values]
lines!(ax3, r1_values, r2_values, z_values, linewidth = gridwidth, linestyle = gridsymbol, color = (gridcolor, gridalpha), overdraw=false, transparency = gridtransparecy)
lines!(ax3, r1_values, r2_values, z_values, linewidth = gridwidth, linestyle = gridsymbol, color = (gridcolor, .2), overdraw=true, transparency = gridoverdrawtransparency)

r2_values = 0:0.001:xmax
r1_values = repeat([rhigh1], length(r2_values))
z_values = [W(1/2, r, rhigh1)+gridoffset for r in r2_values]
lines!(ax3, r1_values, r2_values, z_values, linewidth = gridwidth, linestyle = gridsymbol, color = (gridcolor, gridalpha), overdraw=false, transparency = gridtransparecy)
lines!(ax3, r1_values, r2_values, z_values, linewidth = gridwidth, linestyle = gridsymbol, color = (gridcolor, .2), overdraw=true, transparency = gridoverdrawtransparency)

r2_values = 0:0.001:xmax
r1_values = repeat([rlow1], length(r2_values))
z_values = [W(1/2, r, rlow1)+gridoffset for r in r2_values]
lines!(ax3, r1_values, r2_values, z_values, linewidth = gridwidth, linestyle = gridsymbol, color = (gridcolor, gridalpha), overdraw=false, transparency = gridtransparecy)
lines!(ax3, r1_values, r2_values, z_values, linewidth = gridwidth, linestyle = gridsymbol, color = (gridcolor, .2), overdraw=true, transparency = gridoverdrawtransparency)

r1_values = 0:0.001:xmax
r2_values = repeat([rlow2 + .45], length(r1_values))
z_values = [W(1/2, rlow2 + .45, r)+gridoffset for r in r1_values]
lines!(ax3, r1_values, r2_values, z_values, linewidth = gridwidth, linestyle = gridsymbol, color = (gridcolor, gridalpha), overdraw=false, transparency = gridtransparecy)
lines!(ax3, r1_values, r2_values, z_values, linewidth = gridwidth, linestyle = gridsymbol, color = (gridcolor, .2), overdraw=true, transparency = gridoverdrawtransparency)

r1_values = 0:0.001:xmax
r2_values = repeat([rhigh2], length(r1_values))
z_values = [W(1/2, rhigh2, r)+gridoffset for r in r1_values]
lines!(ax3, r1_values, r2_values, z_values, linewidth = gridwidth, linestyle = gridsymbol, color = (gridcolor, gridalpha), overdraw=false, transparency = gridtransparecy)
lines!(ax3, r1_values, r2_values, z_values, linewidth = gridwidth, linestyle = gridsymbol, color = (gridcolor, .2), overdraw=true, transparency = gridoverdrawtransparency)

r2_values = 0:0.001:xmax
r1_values = repeat([rhigh2], length(r2_values))
z_values = [W(1/2, r, rhigh2)+gridoffset for r in r2_values]
lines!(ax3, r1_values, r2_values, z_values, linewidth = gridwidth, linestyle = gridsymbol, color = (gridcolor, gridalpha), overdraw=false, transparency = gridtransparecy)
lines!(ax3, r1_values, r2_values, z_values, linewidth = gridwidth, linestyle = gridsymbol, color = (gridcolor, .2), overdraw=true, transparency = gridoverdrawtransparency)

r2_values = 0:0.001:xmax
r1_values = repeat([rlow2+.45], length(r2_values))
z_values = [W(1/2, r, rlow2+.45)+gridoffset for r in r2_values]
lines!(ax3, r1_values, r2_values, z_values, linewidth = gridwidth, linestyle = gridsymbol, color = (gridcolor, gridalpha), overdraw=false, transparency = gridtransparecy)
lines!(ax3, r1_values, r2_values, z_values, linewidth = gridwidth, linestyle = gridsymbol, color = (gridcolor, .2), overdraw=true, transparency = gridoverdrawtransparency)


# Find the maxima 
max_val = -Inf
max_r1, max_r2 = -1, -1

for r1 in rlow:.1:rhigh
    for r2 in rlow:.1:rhigh
        val = W(1/2, r1, r2)
        if val > max_val
            max_val = val
            max_r1, max_r2 = r1, r2
        end
    end
end

max_val += .003

# Plot the maxima. There will be two because the landscape is symmetric
scatter!(ax3, [max_r1], [max_r2], [max_val], markersize = markersize*.8, color = (maxima_color, .8), overdraw = false, marker = maxima_marker)
scatter!(ax3, [max_r2], [max_r1], [max_val], markersize = markersize*.8, color = (maxima_color, .8), overdraw = false, marker = maxima_marker)
# Replot with transparency and overdraw for asthetics 
scatter!(ax3, [max_r1], [max_r2], [max_val], markersize = markersize, color = (maxima_color, .4), overdraw = false, marker = maxima_marker, transparency = true, ssao = true)
scatter!(ax3, [max_r2], [max_r1], [max_val], markersize = markersize, color = (maxima_color, .4), overdraw = false, marker = maxima_marker, transparency = true, ssao = true)
scatter!(ax3, [max_r1], [max_r2], [max_val], markersize = markersize, color = (maxima_color, .4), overdraw = true, marker = maxima_marker, transparency = true, ssao = true)
scatter!(ax3, [max_r2], [max_r1], [max_val], markersize = markersize, color = (maxima_color, .4), overdraw = true, marker = maxima_marker, transparency = true, ssao = true)

# generate points where r1 = r2 to show symmetric strategies
r_values = 0:0.001:xmax-.1
z_values = [W(1/2, r, r)+.004 for r in r_values]

# plot the line where r1=r2. 
# we need to plot multiple lines with varying values and options to get the transparency to work out correctly
lines!(ax3, r_values, r_values, z_values, linewidth = symwidth1, color = (maxima_color, .2), overdraw=false, ssao = true)
lines!(ax3, r_values, r_values, z_values, linewidth = symwidth1, color = (maxima_color, .2), overdraw=true, ssao = true)
lines!(ax3, r_values, r_values, z_values, linewidth = symwidth2, color = (maxima_color, .4), overdraw=false, transparency =true, ssao = true)
lines!(ax3, r_values, r_values, z_values, linewidth = symwidth2, color = (maxima_color, .6), overdraw=true, transparency=true, ssao = true)


save("Landscapes2.png", f)

f  

# %% [markdown]
### Part 3

# %% [code]


# ===========================================================================
# ===========ONE CONVEX, ONE CONCAVE, DIFFERENT RLOW AND RHIGH===============
# ===========================================================================

f = Figure()

function s1_curve(r)
    d=4
    shift = 0
     return .3 - max(0.5 * (1 - exp(-d * (r - shift - rlow)) - (r-shift) * 1/8), 0) + .05
end

function s2_curve(r)
    d = 1
    r = rmax-r  # Mirror the function
    shift = 7
    out = max((1 - 1.4^(-d * (r - shift - rlow))) * 0.7 - 1/16 * (r - shift) , 0) + .2
end


# Set the parameters
rlow1 = 2
rhigh1 = 6
rlow2 = 4
rhigh2 = 11.55 

s1(r) = add_flat_regions(s1_curve, r, rlow1, rhigh1)
s2(r) = add_flat_regions(s2_curve, r, rlow2, rhigh2)


# ------------ Survivorship Functions ---------------------------------------------------
top = .4  # how far the bands reach

ax1 = Axis(
    f[1,1],
    xlabel = "r",
    ylabel = "s₁(r) and s₂(r) ",
    aspect = 1,
    limits = (0, xmax, 0, top),
    xticks = ([rlow1, rhigh1, rlow2  + .45, rhigh2, xmax],   [L"r_{low}", L"r_{high}", L"r'_{low}", L"r'_{high}", L"r_{max}"]),  # Have to shift the rlow2 tick a bit. Don't know why.,  
    yticks = [0],
    xticklabelsize = ticksize,
    yticklabelsize = ticksize,

)
  
# Seperate the regions by bands
alpha = .3  # How faint the bands are
band!(
    [0, rlow1],
    [0, 0],
    [top , top],
    color = (color_cycle[1], alpha)
)
band!(
    [rlow1, rlow2 + .45],
    [0, 0],
    [top , top],
    color = ( color_cycle[2], alpha)
)
band!(
    [rlow2 + .45, rhigh1],
    [0, 0],
    [top, top],
    color = ( color_cycle[3], alpha)
)
band!(
    [rhigh1, rhigh2],
    [0, 0],
    [top, top],
    color = ( color_cycle[4], alpha)
)
band!(
    [rhigh2, xmax],
    [0, 0],
    [top, top],
    color = (color_cycle[5], alpha)
)

# Boundry Lines
# Change the alpha manually so that the lines remain preceptually constant between the regions
vlines!(rlow1, linestyle = gridsymbol, color = (gridcolor, 1), linewidth = gridwidthflat)
vlines!(rhigh1, linestyle = gridsymbol, color = (gridcolor, .6), linewidth = gridwidthflat)
vlines!(rlow2 + .45, linestyle = gridsymbol, color = (gridcolor, .9), linewidth = gridwidthflat)
vlines!(rhigh2, linestyle = gridsymbol, color = (gridcolor, .5), linewidth = gridwidthflat)

# Plot the survivorship functions (put the fainter line behind for a sense of depth)
lines!(
    0:.01:xmax, 
    s2,
    linewidth = lw,
    color = Cycled(2)  # An orange color in the Algebra of Graphics theme
    )
lines!(
    0:.001:xmax, 
    s1,
    linewidth = lw,
    color = survivorship_color
    )
    
# --------------------- The Grid of regions ---------------------------------------------

ax2 = Axis(
    f[1,2],
    xlabel = "r₁",
    ylabel = "r₂",
    aspect = 1,
    limits = (0, xmax, 0, xmax),
    xticks = ([rlow1, rhigh1, rlow2  + .45, rhigh2, xmax],  [L"r_{low}", L"r_{high}", L"r'_{low}", L"r'_{high}", L"r_{max}"]),  
    yticks = ([rlow1, rhigh1, rlow2  + .45, rhigh2, xmax], [L"r_{low}", L"r_{high}", L"r'_{low}", L"r'_{high}", L"r_{max}"]),  
    xticklabelsize = ticksize,
    yticklabelsize = ticksize,
     
)

# Region for surface 
# X-axis
alpha2 = .5
top = xmax
band!(
    [0, rlow1],
    [0, 0],
    [top , top],
    color = (color_cycle[1], alpha2),
    transparency = true
)
band!(
    [rlow1, rlow2 + .45],
    [0, 0],
    [top , top],
    color = ( color_cycle[2], alpha2),
    transparency = true
)
band!(
    [rlow2 + .45, rhigh1],
    [0, 0],
    [top, top],
    color = ( color_cycle[3], alpha2),
    transparency = true
)
band!(
    [rhigh1, rhigh2],
    [0, 0],
    [top, top],
    color = ( color_cycle[4], alpha2),
    transparency = true
)
band!(
    [rhigh2, xmax],
    [0, 0],
    [top, top],
    color = (color_cycle[5], alpha2),
    transparency = true
)

# Y axis
band!(
    [0, xmax],
    [rlow1, rlow1],
    [0 , 0],
    color = (color_cycle[1], alpha2),
    transparency = true
)
band!(
    [0, xmax],
    [rlow2+.45, rlow2+.45],
    [rlow1, rlow1],
    color = (color_cycle[2], alpha2),
    transparency = true
)
band!(
    [0, xmax],
    [rhigh1, rhigh1],
    [rlow2 + .45, rlow2 + .45],
    color = (color_cycle[3], alpha2),
    transparency = true
)
band!(
    [0, xmax],
    [rhigh2, rhigh2],
    [rhigh1, rhigh1],
    color = (color_cycle[4], alpha2),
    transparency = true
)
band!(
    [0, xmax],
    [xmax, xmax],
    [rhigh2, rhigh2],
    color = (color_cycle[5], alpha2),
    transparency = true
)

# Gridlines
vlines!(rlow1, linestyle = gridsymbol, color = gridcolor, linewidth = gridwidthflat, transparency = true)
vlines!(rhigh1, linestyle = gridsymbol, color = gridcolor, linewidth = gridwidthflat, transparency = true)
hlines!(rlow1, linestyle = gridsymbol, color = gridcolor, linewidth = gridwidthflat, transparency = true)
hlines!(rhigh1, linestyle = gridsymbol, color = gridcolor, linewidth = gridwidthflat, transparency = true)
vlines!(rlow2+.45, linestyle = gridsymbol, color = gridcolor, linewidth = gridwidthflat, transparency = true)
vlines!(rhigh2, linestyle = gridsymbol, color = gridcolor, linewidth = gridwidthflat, transparency = true)
hlines!(rlow2+.45, linestyle = gridsymbol, color = gridcolor, linewidth = gridwidthflat, transparency = true)
hlines!(rhigh2, linestyle = gridsymbol, color = gridcolor, linewidth = gridwidthflat, transparency = true)

# ---------- Fitness Landscape ---------------------------------------------------

R = 25
# I can't get Latex to work with the tick lables for the 3D plot. So we will add those in after with photoshop
ax3 = Axis3(
    f[1,3 ],
    azimuth =  -π/2  -.2,
    elevation = .9,
    xlabel = "r₁",
    ylabel = "r₂",
    zlabel = "W(r₁, r₂)",
    #aspect = (1,1,1),
    xticklabelsvisible = false,
    yticklabelsvisible = false,
    zticklabelsvisible = false,
    zticksvisible = false,
    xticks = [rlow1, rhigh1, rlow2  + .45, rhigh2, xmax],  
    yticks = [rlow1, rhigh1, rlow2  + .45, rhigh2, xmax], 
    xticklabelsize = ticksize,
    yticklabelsize = ticksize,
)

# Make the surface
regions = [0, rlow1, rlow2  + .45, rhigh1,  rhigh2, xmax]
for (i, r1) in enumerate(regions)
    for (j, r2) in enumerate(regions)
        space = wirespace
        if i < 6 && j < 6
            r1s = regions[i]:space:regions[i+1]  # x axis
            r2s = regions[j]:space:regions[j+1] # y axis
            zs = [W(1/2,  r1,  r2) for r1 in r1s, r2 in r2s]  # z axis

            # Plot the three of these
            
            wireframe!(
                    r1s, 
                    r2s, 
                    zs,
                    color = (color_cycle[i], .5),
                    linewidth = wirewidth,
                    transparency = true
                    )
            wireframe!(
            r1s, 
            r2s, 
            zs,
            color = (color_cycle[j], .5),
            linewidth = wirewidth,
            transparency = true
            )
            surface!(
                r1s, 
                r2s, 
                zs,
                colormap = (:greys, surfacealpha),
                )
        end
    end
end


# Plot the section boundries 
r1_values = 0:0.001:xmax
r2_values = repeat([rlow1], length(r1_values))
z_values = [W(1/2, rlow1, r)+gridoffset for r in r1_values]
lines!(ax3, r1_values, r2_values, z_values, linewidth = gridwidth, linestyle = gridsymbol, color = (gridcolor, gridalpha), overdraw=false, transparency = gridtransparecy)
lines!(ax3, r1_values, r2_values, z_values, linewidth = gridwidth, linestyle = gridsymbol, color = (gridcolor, .2), overdraw=true, transparency = gridoverdrawtransparency)

r1_values = 0:0.001:xmax
r2_values = repeat([rhigh1], length(r1_values))
z_values = [W(1/2, rhigh1, r)+gridoffset for r in r1_values]
lines!(ax3, r1_values, r2_values, z_values, linewidth = gridwidth, linestyle = gridsymbol, color = (gridcolor, gridalpha), overdraw=false, transparency = gridtransparecy)
lines!(ax3, r1_values, r2_values, z_values, linewidth = gridwidth, linestyle = gridsymbol, color = (gridcolor, .2), overdraw=true, transparency = gridoverdrawtransparency)

r2_values = 0:0.001:xmax
r1_values = repeat([rhigh1], length(r2_values))
z_values = [W(1/2, r, rhigh1)+gridoffset for r in r2_values]
lines!(ax3, r1_values, r2_values, z_values, linewidth = gridwidth, linestyle = gridsymbol, color = (gridcolor, gridalpha), overdraw=false, transparency = gridtransparecy)
lines!(ax3, r1_values, r2_values, z_values, linewidth = gridwidth, linestyle = gridsymbol, color = (gridcolor, .2), overdraw=true, transparency = gridoverdrawtransparency)

r2_values = 0:0.001:xmax
r1_values = repeat([rlow1], length(r2_values))
z_values = [W(1/2, r, rlow1)+gridoffset for r in r2_values]
lines!(ax3, r1_values, r2_values, z_values, linewidth = gridwidth, linestyle = gridsymbol, color = (gridcolor, gridalpha), overdraw=false, transparency = gridtransparecy)
lines!(ax3, r1_values, r2_values, z_values, linewidth = gridwidth, linestyle = gridsymbol, color = (gridcolor, .2), overdraw=true, transparency = gridoverdrawtransparency)

r1_values = 0:0.001:xmax
r2_values = repeat([rlow2 + .45], length(r1_values))
z_values = [W(1/2, rlow2 + .45, r)+gridoffset for r in r1_values]
lines!(ax3, r1_values, r2_values, z_values, linewidth = gridwidth, linestyle = gridsymbol, color = (gridcolor, gridalpha), overdraw=false, transparency = gridtransparecy)
lines!(ax3, r1_values, r2_values, z_values, linewidth = gridwidth, linestyle = gridsymbol, color = (gridcolor, .2), overdraw=true, transparency = gridoverdrawtransparency)

r1_values = 0:0.001:xmax
r2_values = repeat([rhigh2], length(r1_values))
z_values = [W(1/2, rhigh2, r)+gridoffset for r in r1_values]
lines!(ax3, r1_values, r2_values, z_values, linewidth = gridwidth, linestyle = gridsymbol, color = (gridcolor, gridalpha), overdraw=false, transparency = gridtransparecy)
lines!(ax3, r1_values, r2_values, z_values, linewidth = gridwidth, linestyle = gridsymbol, color = (gridcolor, .2), overdraw=true, transparency = gridoverdrawtransparency)

r2_values = 0:0.001:xmax
r1_values = repeat([rhigh2], length(r2_values))
z_values = [W(1/2, r, rhigh2)+gridoffset for r in r2_values]
lines!(ax3, r1_values, r2_values, z_values, linewidth = gridwidth, linestyle = gridsymbol, color = (gridcolor, gridalpha), overdraw=false, transparency = gridtransparecy)
lines!(ax3, r1_values, r2_values, z_values, linewidth = gridwidth, linestyle = gridsymbol, color = (gridcolor, .2), overdraw=true, transparency = gridoverdrawtransparency)

r2_values = 0:0.001:xmax
r1_values = repeat([rlow2+.45], length(r2_values))
z_values = [W(1/2, r, rlow2+.45)+gridoffset for r in r2_values]
lines!(ax3, r1_values, r2_values, z_values, linewidth = gridwidth, linestyle = gridsymbol, color = (gridcolor, gridalpha), overdraw=false, transparency = gridtransparecy)
lines!(ax3, r1_values, r2_values, z_values, linewidth = gridwidth, linestyle = gridsymbol, color = (gridcolor, .2), overdraw=true, transparency = gridoverdrawtransparency)


# Find the maxima 
max_val = -Inf
max_r1, max_r2 = -1, -1

for r1 in rlow:.1:rhigh
    for r2 in rlow:.05:rhigh
        val = W(1/2, r1, r2)
        if val > max_val
            max_val = val
            max_r1, max_r2 = r1, r2
        end
    end
end

max_val += .01


# Plot the maxima. There will be two because the landscape is symmetric
scatter!(ax3, [max_r1], [max_r2], [max_val], markersize = markersize*.8, color = (maxima_color, .8), overdraw = false, marker = maxima_marker)
scatter!(ax3, [max_r2], [max_r1], [max_val], markersize = markersize*.8, color = (maxima_color, .8), overdraw = false, marker = maxima_marker)
# Replot with transparency and overdraw for asthetics 
scatter!(ax3, [max_r1], [max_r2], [max_val], markersize = markersize, color = (maxima_color, .4), overdraw = false, marker = maxima_marker, transparency = true, ssao = true)
scatter!(ax3, [max_r2], [max_r1], [max_val], markersize = markersize, color = (maxima_color, .4), overdraw = false, marker = maxima_marker, transparency = true, ssao = true)
scatter!(ax3, [max_r1], [max_r2], [max_val], markersize = markersize, color = (maxima_color, .4), overdraw = true, marker = maxima_marker, transparency = true, ssao = true)
scatter!(ax3, [max_r2], [max_r1], [max_val], markersize = markersize, color = (maxima_color, .4), overdraw = true, marker = maxima_marker, transparency = true, ssao = true)

# generate points where r1 = r2 to show symmetric strategies
r_values = 0:0.001:xmax-.1
z_values = [W(1/2, r, r)+.004 for r in r_values]

# plot the line where r1=r2. 
# we need to plot multiple lines with varying values and options to get the transparency to work out correctly
lines!(ax3, r_values, r_values, z_values, linewidth = symwidth1, color = (maxima_color, .2), overdraw=false, ssao = true)
lines!(ax3, r_values, r_values, z_values, linewidth = symwidth1, color = (maxima_color, .2), overdraw=true, ssao = true)
lines!(ax3, r_values, r_values, z_values, linewidth = symwidth2, color = (maxima_color, .4), overdraw=false, transparency =true, ssao = true)
lines!(ax3, r_values, r_values, z_values, linewidth = symwidth2, color = (maxima_color, .6), overdraw=true, transparency=true, ssao = true)


save("Landscapes3.png", f)

f


# %% [markdown]
### All three parts together

Here is the code for all three parts together to make an interactive version.


# %% [code]
GLMakie.activate!(inline=false)  # We will use GLMakie because it allows finer control of transparency through overdraw 


update_theme!(                  # Tweaks the current theme
           fontsize=38,
           markersize=50,
           resolution = (2000, 2000),

       )

rlow = 2
rmax = 22

ticksize = 38


# Plot the survival function, but with flat regions added
function add_flat_regions(s, r, rlow, rhigh)

    if r < rlow
        return s(rlow)
    elseif r > rhigh
        return s(rhigh)
    else
        return s(r)
    end

end


f = Figure()  # Setup figure
xmax = 18  # How far to plot
lw = 6  # Line width
top = .4  # how far the bands reach
wirewidth = 5  # width of the wireframe mesh. This is thick enough that it appears as a solid surface
wirespace = .05 
surfacealpha = .3  # transparecy of grey background surface

gridoffset = .00
gridcolor = :white
gridwidth = 6  # width of the gridlines in the rightmost plot
gridsymbol = :solid  # style of the gridlines
gridtransparecy = true  # whether the grid is transparent
gridoverdrawtransparency = true  # whether the overdraw grid is transparent
gridalpha = .8  # alpha of the 
gridwidthflat = 3  # Size of the gridlines for the middle column plots

# ===========TWO CONCAVE, SAME RLOW AND RHIGH ====================================================

# ------------- Define the Functions ------------------------------------------

function s1_curve(r)
    d=4
    shift = -1.5
    x = .4
    return max(0.5 * (1 - exp(-d * (r*x - shift - rlow)) - (r*x-shift) * 1/8), 0) 
end

function s2_curve(r)
    d = 1
    r = rmax-r  # Mirror the function
    shift = 1.3
    x = .6
    out =max((1 - 1.4^(-d * (r*x - shift - rlow))) * 0.7 - 1/16 * (r*x - shift) , -.075) + .075
end


# Set the parameters
rlow = 2.1
rhigh = 14

# Add the flat regions into the survivorship functions
s1(r) = add_flat_regions(s1_curve, r, rlow, rhigh)
s2(r) = add_flat_regions(s2_curve, r, rlow, rhigh)

# ------------ Survivorship Functions ---------------------------------------------------

ax1 = Axis(
    f[1,1],
    xlabel = "r",
    ylabel = "s₁(r) and s₂(r) ",
    aspect = 1,
    limits = (0, xmax, 0, top),
    xticks = ([rlow, rhigh, xmax],   [L"r_{low}", L"r_{high}",  L"r_{max}"]),  # Have to shift the rlow2 tick a bit. Don't know why.,  
    yticks = [0],
    xticklabelsize = ticksize,
    yticklabelsize = ticksize,

)
  
# Seperate the regions by bands
alpha = .3  # How faint the bands are
band!(
    [0, rlow],
    [0, 0],
    [top , top],
    color = (color_cycle[1], alpha),
)
band!(
    [rlow, rhigh],
    [0, 0],
    [top , top],
    color = ( color_cycle[3], alpha),
)
band!(
    [rhigh, xmax],
    [0, 0],
    [top, top],
    color = ( color_cycle[5], alpha),
)

# Boundry Lines
vlines!(rlow, linestyle = gridsymbol, color = (gridcolor, 1), linewidth = gridwidthflat)
vlines!(rhigh, linestyle = gridsymbol, color = (gridcolor, .6), linewidth = gridwidthflat)

# Plot the survivorship functions (put the fainter line behind for a sense of depth)
lines!(
    0:.01:xmax, 
    s2,
    linewidth = lw,
    color = Cycled(2)  # An orange color in the Algebra of Graphics theme
    )
lines!(
    0:.001:xmax, 
    s1,
    linewidth = lw,
    color = survivorship_color
    )
    
# --------------------- The Grid of regions ---------------------------------------------

ax2 = Axis(
    f[1,2],
    xlabel = "r₁",
    ylabel = "r₂",
    aspect = 1,
    limits = (0, xmax, 0, xmax),
    xticks =  ([rlow, rhigh, xmax],   [L"r_{low}", L"r_{high}",  L"r_{max}"]),
    yticks =  ([rlow, rhigh, xmax],   [L"r_{low}", L"r_{high}",  L"r_{max}"]),
    xticklabelsize = ticksize,
    yticklabelsize = ticksize,
     
)

# Region for surface 
# X-axis
alpha = .5
top = xmax
band!(
    [0, rlow],
    [0, 0],
    [top , top],
    color = (color_cycle[1], alpha),
    transparency = true
)
band!(
    [rlow, rhigh],
    [0, 0],
    [top , top],
    color = ( color_cycle[3], alpha),
    transparency = true
)
band!(
    [rhigh, xmax],
    [0, 0],
    [top, top],
    color = ( color_cycle[5], alpha),
    transparency = true
)

# Y axis
band!(
    [0, xmax],
    [rlow, rlow],
    [0 , 0],
    color = (color_cycle[1], alpha),
    transparency = true
)

band!(
    [0, xmax],
    [rhigh, rhigh],
    [rlow, rlow],
    color = (color_cycle[3], alpha),
    transparency = true
)
band!(
    [0, xmax],
    [xmax, xmax],
    [rhigh, rhigh],
    color = (color_cycle[5], alpha),
    transparency = true
)

# Gridlines 

vlines!(rlow, linestyle = gridsymbol, color = gridcolor, linewidth = gridwidthflat, transparency = true)
vlines!(rhigh, linestyle = gridsymbol, color = gridcolor, linewidth = gridwidthflat, transparency = true)
hlines!(rlow, linestyle = gridsymbol, color = gridcolor, linewidth = gridwidthflat, transparency = true)
hlines!(rhigh, linestyle = gridsymbol, color = gridcolor, linewidth = gridwidthflat, transparency = true)

# ---------- Fitness Landscape ---------------------------------------------------

R = 25
# I can't get Latex to work with the tick lables for the 3D plot. So we will add those in after with photoshop
ax3 = Axis3(
    f[1,3 ],
    azimuth =  -π/2  -.2,
    elevation = .9,
    xlabel = "r₁",
    ylabel = "r₂",
    zlabel = "W(r₁, r₂)",
    #aspect = (1,1,1),
    xticklabelsvisible = false,
    yticklabelsvisible = false,
    zticklabelsvisible = false,
    zticksvisible = false,
    xticks = [rlow, rhigh, xmax],  
    yticks = [rlow, rhigh, xmax],   
    xticklabelsize = ticksize,
    yticklabelsize = ticksize,
)

# Make the surface. Enumerate through each region of the middle plot
regions = [0, rlow,  rhigh, xmax]
for (i, r1) in enumerate(regions)
    for (j, r2) in enumerate(regions)
        space = wirespace
        if i < 4 && j < 4
            r1s = regions[i]:space:regions[i+1]  # x axis
            r2s = regions[j]:space:regions[j+1] # y axis
            zs = [W(1/2,  r1,  r2) for r1 in r1s, r2 in r2s]  # z axis

            #Plot the three of these
           
            wireframe!(
                    r1s, 
                    r2s, 
                    zs,
                    color = (color_cycle[i*2-1], .5),
                    transparency = true,
                    linewidth = wirewidth
                    )
                wireframe!(
                r1s, 
                r2s, 
                zs,
                color = (color_cycle[j*2-1], .5),
                linewidth = wirewidth,
                transparency = true,
                )
                surface!(
                    r1s, 
                    r2s, 
                    zs,
                    colormap = (:greys, surfacealpha),
                    )
    
        end
    end
end


# Plot the section boundries

r1_values = 0:0.001:xmax
r2_values = repeat([rlow], length(r1_values))
z_values = [W(1/2, rlow, r)+gridoffset for r in r1_values]
lines!(ax3, r1_values, r2_values, z_values, linewidth = gridwidth, linestyle = gridsymbol, color = (gridcolor, .5), overdraw=false, transparency = gridtransparecy)
lines!(ax3, r1_values, r2_values, z_values, linewidth = gridwidth, linestyle = gridsymbol, color = (gridcolor, .2), overdraw=true, transparency = gridoverdrawtransparency)

r1_values = 0:0.001:xmax
r2_values = repeat([rhigh], length(r1_values))
z_values = [W(1/2, rhigh, r)+gridoffset for r in r1_values]
lines!(ax3, r1_values, r2_values, z_values, linewidth = gridwidth, linestyle = gridsymbol, color = (gridcolor, .5), overdraw=false, transparency = gridtransparecy)
lines!(ax3, r1_values, r2_values, z_values, linewidth = gridwidth, linestyle = gridsymbol, color = (gridcolor, .2), overdraw=true, transparency = gridoverdrawtransparency)

r2_values = 0:0.001:xmax
r1_values = repeat([rhigh], length(r2_values))
z_values = [W(1/2, r, rhigh)+gridoffset for r in r2_values]
lines!(ax3, r1_values, r2_values, z_values, linewidth = gridwidth, linestyle = gridsymbol, color = (gridcolor, .5), overdraw=false, transparency = gridtransparecy)
lines!(ax3, r1_values, r2_values, z_values, linewidth = gridwidth, linestyle = gridsymbol, color = (gridcolor, .2), overdraw=true, transparency = gridoverdrawtransparency)

r2_values = 0:0.001:xmax
r1_values = repeat([rlow], length(r2_values))
z_values = [W(1/2, r, rlow)+gridoffset for r in r2_values]
lines!(ax3, r1_values, r2_values, z_values, linewidth = gridwidth, linestyle = gridsymbol, color = (gridcolor, .5), overdraw=false, transparency = gridtransparecy)
lines!(ax3, r1_values, r2_values, z_values, linewidth = gridwidth, linestyle = gridsymbol, color = (gridcolor, .2), overdraw=true, transparency = gridoverdrawtransparency)


# Find the maxima 
max_val = -Inf
max_r1, max_r2 = -1, -1

for r1 in rlow:.1:rhigh
    for r2 in rlow:.1:rhigh
        val = W(1/2, r1, r2)
        if val > max_val
            max_val = val
            max_r1, max_r2 = r1, r2
        end
    end
end

max_val += .004

# Plot the maxima. There will be two because the landscape is symmetric
scatter!(ax3, [max_r1], [max_r2], [max_val], markersize = 16, color = (maxima_color, .8), overdraw = false, marker = maxima_marker)
scatter!(ax3, [max_r2], [max_r1], [max_val], markersize = 16, color = (maxima_color, .8), overdraw = false, marker = maxima_marker)
# Replot with transparency and overdraw for asthetics 
scatter!(ax3, [max_r1], [max_r2], [max_val], markersize = 20, color = (maxima_color, .4), overdraw = false, marker = maxima_marker, transparency = true, ssao = true)
scatter!(ax3, [max_r2], [max_r1], [max_val], markersize = 20, color = (maxima_color, .4), overdraw = false, marker = maxima_marker, transparency = true, ssao = true)
scatter!(ax3, [max_r1], [max_r2], [max_val], markersize = 20, color = (maxima_color, .4), overdraw = true, marker = maxima_marker, transparency = true, ssao = true)
scatter!(ax3, [max_r2], [max_r1], [max_val], markersize = 20, color = (maxima_color, .4), overdraw = true, marker = maxima_marker, transparency = true, ssao = true)

# generate points where r1 = r2 to show symmetric strategies
r_values = 0:0.001:xmax-.1
z_values = [W(1/2, r, r)+.004 for r in r_values]

# plot the line where r1=r2. 
# we need to plot multiple lines with varying values and options to get the transparency to work out correctly
lines!(ax3, r_values, r_values, z_values, linewidth = 3, color = (maxima_color, .2), overdraw=false, ssao = true)
lines!(ax3, r_values, r_values, z_values, linewidth = 3, color = (maxima_color, .2), overdraw=true, ssao = true)
lines!(ax3, r_values, r_values, z_values, linewidth = 8, color = (maxima_color, .4), overdraw=false, transparency =true, ssao = true)
lines!(ax3, r_values, r_values, z_values, linewidth = 8, color = (maxima_color, .6), overdraw=true, transparency=true, ssao = true)


# ===========================================================================
# ===========TWO CONCAVE, DIFFERENT RLOW AND RHIGH==============================
# ===========================================================================

function s1_curve(r)
    d=4
    shift = 0
     return max(0.5 * (1 - exp(-d * (r - shift - rlow)) - (r-shift) * 1/8), 0) + .05
end

function s2_curve(r)
    d = 1
    r = rmax-r  # Mirror the function
    shift = 7
    out = max((1 - 1.4^(-d * (r - shift - rlow))) * 0.7 - 1/16 * (r - shift) , 0) + .025
end


# Set the parameters
rlow1 = 2.25
rhigh1 = 8
rlow2 = 4
rhigh2 = 11.55

s1(r) = add_flat_regions(s1_curve, r, rlow1, rhigh1)
s2(r) = add_flat_regions(s2_curve, r, rlow2, rhigh2)


# ------------ Survivorship Functions ---------------------------------------------------
top = .4  # how far the bands reach

ax1 = Axis(
    f[2,1],
    xlabel = "r",
    ylabel = "s₁(r) and s₂(r) ",
    aspect = 1,
    limits = (0, xmax, 0, top),
    xticks = ([rlow1, rhigh1, rlow2  + .45, rhigh2, xmax],   [L"r_{low}", L"r_{high}", L"r'_{low}", L"r'_{high}", L"r_{max}"]),  # Have to shift the rlow2 tick a bit. Don't know why.,  
    yticks = [0],
    xticklabelsize = ticksize,
    yticklabelsize = ticksize,

)
  
# Seperate the regions by bands
alpha = .3  # How faint the bands are
band!(
    [0, rlow1],
    [0, 0],
    [top , top],
    color = (color_cycle[1], alpha),
)
band!(
    [rlow1, rlow2 + .45],
    [0, 0],
    [top , top],
    color = ( color_cycle[2], alpha)
)
band!(
    [rlow2 + .45, rhigh1],
    [0, 0],
    [top, top],
    color = ( color_cycle[3], alpha)
)
band!(
    [rhigh1, rhigh2],
    [0, 0],
    [top, top],
    color = ( color_cycle[4], alpha)
)
band!(
    [rhigh2, xmax],
    [0, 0],
    [top, top],
    color = (color_cycle[5], alpha)
)

# Boundry Lines
# Change the alpha manually so that the lines remain preceptually constant between the regions
vlines!(rlow1, linestyle = gridsymbol, color = (gridcolor, 1), linewidth = gridwidthflat)
vlines!(rhigh1, linestyle = gridsymbol, color = (gridcolor, .6), linewidth = gridwidthflat)
vlines!(rlow2 + .45, linestyle = gridsymbol, color = (gridcolor, .9), linewidth = gridwidthflat)
vlines!(rhigh2, linestyle = gridsymbol, color = (gridcolor, .5), linewidth = gridwidthflat)


# Plot the survivorship functions (put the fainter line behind for a sense of depth)
lines!(
    0:.01:xmax, 
    s2,
    linewidth = lw,
    color = Cycled(2)  # An orange color in the Algebra of Graphics theme
    )
lines!(
    0:.001:xmax, 
    s1,
    linewidth = lw,
    color = survivorship_color
    )
    
# --------------------- The Grid of regions ---------------------------------------------

ax2 = Axis(
    f[2,2],
    xlabel = "r₁",
    ylabel = "r₂",
    aspect = 1,
    limits = (0, xmax, 0, xmax),
    xticks = ([rlow1, rhigh1, rlow2  + .45, rhigh2, xmax],  [L"r_{low}", L"r_{high}", L"r'_{low}", L"r'_{high}", L"r_{max}"]),  
    yticks = ([rlow1, rhigh1, rlow2  + .45, rhigh2, xmax], [L"r_{low}", L"r_{high}", L"r'_{low}", L"r'_{high}", L"r_{max}"]),  
    xticklabelsize = ticksize,
    yticklabelsize = ticksize,
     
)

# Region for surface 
# X-axis
alpha2 = .5
top = xmax
band!(
    [0, rlow1],
    [0, 0],
    [top , top],
    color = (color_cycle[1], alpha2),
    transparency = true
)
band!(
    [rlow1, rlow2 + .45],
    [0, 0],
    [top , top],
    color = ( color_cycle[2], alpha2),
    transparency = true
)
band!(
    [rlow2 + .45, rhigh1],
    [0, 0],
    [top, top],
    color = ( color_cycle[3], alpha2),
    transparency = true
)
band!(
    [rhigh1, rhigh2],
    [0, 0],
    [top, top],
    color = ( color_cycle[4], alpha2),
    transparency = true
)
band!(
    [rhigh2, xmax],
    [0, 0],
    [top, top],
    color = (color_cycle[5], alpha2),
    transparency = true
)

# Y axis
band!(
    [0, xmax],
    [rlow1, rlow1],
    [0 , 0],
    color = (color_cycle[1], alpha2),
    transparency = true
)
band!(
    [0, xmax],
    [rlow2+.45, rlow2+.45],
    [rlow1, rlow1],
    color = (color_cycle[2], alpha2),
    transparency = true
)
band!(
    [0, xmax],
    [rhigh1, rhigh1],
    [rlow2 + .45, rlow2 + .45],
    color = (color_cycle[3], alpha2),
    transparency = true
)
band!(
    [0, xmax],
    [rhigh2, rhigh2],
    [rhigh1, rhigh1],
    color = (color_cycle[4], alpha2),
    transparency = true
)
band!(
    [0, xmax],
    [xmax, xmax],
    [rhigh2, rhigh2],
    color = (color_cycle[5], alpha2),
    transparency = true
)

# Gridlines
vlines!(rlow1, linestyle = gridsymbol, color = gridcolor, linewidth = gridwidthflat, transparency = true)
vlines!(rhigh1, linestyle = gridsymbol, color = gridcolor, linewidth = gridwidthflat, transparency = true)
hlines!(rlow1, linestyle = gridsymbol, color = gridcolor, linewidth = gridwidthflat, transparency = true)
hlines!(rhigh1, linestyle = gridsymbol, color = gridcolor, linewidth = gridwidthflat, transparency = true)
vlines!(rlow2+.45, linestyle = gridsymbol, color = gridcolor, linewidth = gridwidthflat, transparency = true)
vlines!(rhigh2, linestyle = gridsymbol, color = gridcolor, linewidth = gridwidthflat, transparency = true)
hlines!(rlow2+.45, linestyle = gridsymbol, color = gridcolor, linewidth = gridwidthflat, transparency = true)
hlines!(rhigh2, linestyle = gridsymbol, color = gridcolor, linewidth = gridwidthflat, transparency = true)
# ---------- Fitness Landscape ---------------------------------------------------

R = 25
# I can't get Latex to work with the tick lables for the 3D plot. So we will add those in after with photoshop
ax3 = Axis3(
    f[2,3 ],
    azimuth =  -π/2  -.2,
    elevation = .9,
    xlabel = "r₁",
    ylabel = "r₂",
    zlabel = "W(r₁, r₂)",
    #aspect = (1,1,1),
    xticklabelsvisible = false,
    yticklabelsvisible = false,
    zticklabelsvisible = false,
    zticksvisible = false,
    xticks = [rlow1, rhigh1, rlow2  + .45, rhigh2, xmax],  
    yticks = [rlow1, rhigh1, rlow2  + .45, rhigh2, xmax], 
    xticklabelsize = ticksize,
    yticklabelsize = ticksize,
)

# Make the surface
regions = [0, rlow1, rlow2  + .45, rhigh1,  rhigh2, xmax]
for (i, r1) in enumerate(regions)
    for (j, r2) in enumerate(regions)
        space = wirespace
        if i < 6 && j < 6
            r1s = regions[i]:space:regions[i+1]  # x axis
            r2s = regions[j]:space:regions[j+1] # y axis
            zs = [W(1/2,  r1,  r2) for r1 in r1s, r2 in r2s]  # z axis

            # Plot the three of these
            
            wireframe!(
                    r1s, 
                    r2s, 
                    zs,
                    color = (color_cycle[i], .5),
                    linewidth = wirewidth,
                    transparency = true
                    )
            wireframe!(
            r1s, 
            r2s, 
            zs,
            color = (color_cycle[j], .5),
            linewidth = wirewidth,
            transparency = true
            )
            surface!(
                r1s, 
                r2s, 
                zs,
                colormap = (:greys, surfacealpha),
                transparency = false
                )
        end
    end
end


# Plot the section boundries 
r1_values = 0:0.001:xmax
r2_values = repeat([rlow1], length(r1_values))
z_values = [W(1/2, rlow1, r)+gridoffset for r in r1_values]
lines!(ax3, r1_values, r2_values, z_values, linewidth = gridwidth, linestyle = gridsymbol, color = (gridcolor, gridalpha), overdraw=false, transparency = gridtransparecy)
lines!(ax3, r1_values, r2_values, z_values, linewidth = gridwidth, linestyle = gridsymbol, color = (gridcolor, .2), overdraw=true, transparency = gridoverdrawtransparency)

r1_values = 0:0.001:xmax
r2_values = repeat([rhigh1], length(r1_values))
z_values = [W(1/2, rhigh1, r)+gridoffset for r in r1_values]
lines!(ax3, r1_values, r2_values, z_values, linewidth = gridwidth, linestyle = gridsymbol, color = (gridcolor, gridalpha), overdraw=false, transparency = gridtransparecy)
lines!(ax3, r1_values, r2_values, z_values, linewidth = gridwidth, linestyle = gridsymbol, color = (gridcolor, .2), overdraw=true, transparency = gridoverdrawtransparency)

r2_values = 0:0.001:xmax
r1_values = repeat([rhigh1], length(r2_values))
z_values = [W(1/2, r, rhigh1)+gridoffset for r in r2_values]
lines!(ax3, r1_values, r2_values, z_values, linewidth = gridwidth, linestyle = gridsymbol, color = (gridcolor, gridalpha), overdraw=false, transparency = gridtransparecy)
lines!(ax3, r1_values, r2_values, z_values, linewidth = gridwidth, linestyle = gridsymbol, color = (gridcolor, .2), overdraw=true, transparency = gridoverdrawtransparency)

r2_values = 0:0.001:xmax
r1_values = repeat([rlow1], length(r2_values))
z_values = [W(1/2, r, rlow1)+gridoffset for r in r2_values]
lines!(ax3, r1_values, r2_values, z_values, linewidth = gridwidth, linestyle = gridsymbol, color = (gridcolor, gridalpha), overdraw=false, transparency = gridtransparecy)
lines!(ax3, r1_values, r2_values, z_values, linewidth = gridwidth, linestyle = gridsymbol, color = (gridcolor, .2), overdraw=true, transparency = gridoverdrawtransparency)

r1_values = 0:0.001:xmax
r2_values = repeat([rlow2 + .45], length(r1_values))
z_values = [W(1/2, rlow2 + .45, r)+gridoffset for r in r1_values]
lines!(ax3, r1_values, r2_values, z_values, linewidth = gridwidth, linestyle = gridsymbol, color = (gridcolor, gridalpha), overdraw=false, transparency = gridtransparecy)
lines!(ax3, r1_values, r2_values, z_values, linewidth = gridwidth, linestyle = gridsymbol, color = (gridcolor, .2), overdraw=true, transparency = gridoverdrawtransparency)

r1_values = 0:0.001:xmax
r2_values = repeat([rhigh2], length(r1_values))
z_values = [W(1/2, rhigh2, r)+gridoffset for r in r1_values]
lines!(ax3, r1_values, r2_values, z_values, linewidth = gridwidth, linestyle = gridsymbol, color = (gridcolor, gridalpha), overdraw=false, transparency = gridtransparecy)
lines!(ax3, r1_values, r2_values, z_values, linewidth = gridwidth, linestyle = gridsymbol, color = (gridcolor, .2), overdraw=true, transparency = gridoverdrawtransparency)

r2_values = 0:0.001:xmax
r1_values = repeat([rhigh2], length(r2_values))
z_values = [W(1/2, r, rhigh2)+gridoffset for r in r2_values]
lines!(ax3, r1_values, r2_values, z_values, linewidth = gridwidth, linestyle = gridsymbol, color = (gridcolor, gridalpha), overdraw=false, transparency = gridtransparecy)
lines!(ax3, r1_values, r2_values, z_values, linewidth = gridwidth, linestyle = gridsymbol, color = (gridcolor, .2), overdraw=true, transparency = gridoverdrawtransparency)

r2_values = 0:0.001:xmax
r1_values = repeat([rlow2+.45], length(r2_values))
z_values = [W(1/2, r, rlow2+.45)+gridoffset for r in r2_values]
lines!(ax3, r1_values, r2_values, z_values, linewidth = gridwidth, linestyle = gridsymbol, color = (gridcolor, gridalpha), overdraw=false, transparency = gridtransparecy)
lines!(ax3, r1_values, r2_values, z_values, linewidth = gridwidth, linestyle = gridsymbol, color = (gridcolor, .2), overdraw=true, transparency = gridoverdrawtransparency)


# Find the maxima 
max_val = -Inf
max_r1, max_r2 = -1, -1

for r1 in rlow:.1:rhigh
    for r2 in rlow:.1:rhigh
        val = W(1/2, r1, r2)
        if val > max_val
            max_val = val
            max_r1, max_r2 = r1, r2
        end
    end
end

max_val += .003

# Plot the maxima. There will be two because the landscape is symmetric
scatter!(ax3, [max_r1], [max_r2], [max_val], markersize = 16, color = (maxima_color, .8), overdraw = false, marker = maxima_marker)
scatter!(ax3, [max_r2], [max_r1], [max_val], markersize = 16, color = (maxima_color, .8), overdraw = false, marker = maxima_marker)
# Replot with transparency and overdraw for asthetics 
scatter!(ax3, [max_r1], [max_r2], [max_val], markersize = 20, color = (maxima_color, .4), overdraw = false, marker = maxima_marker, transparency = true, ssao = true)
scatter!(ax3, [max_r2], [max_r1], [max_val], markersize = 20, color = (maxima_color, .4), overdraw = false, marker = maxima_marker, transparency = true, ssao = true)
scatter!(ax3, [max_r1], [max_r2], [max_val], markersize = 20, color = (maxima_color, .4), overdraw = true, marker = maxima_marker, transparency = true, ssao = true)
scatter!(ax3, [max_r2], [max_r1], [max_val], markersize = 20, color = (maxima_color, .4), overdraw = true, marker = maxima_marker, transparency = true, ssao = true)


# generate points where r1 = r2 to show symmetric strategies
r_values = 0:0.001:xmax-.1
z_values = [W(1/2, r, r)+.004 for r in r_values]

# plot the line where r1=r2. 
# we need to plot multiple lines with varying values and options to get the transparency to work out correctly
lines!(ax3, r_values, r_values, z_values, linewidth = 3, color = (maxima_color, .2), overdraw=false, ssao = true)
lines!(ax3, r_values, r_values, z_values, linewidth = 3, color = (maxima_color, .2), overdraw=true, ssao = true)
lines!(ax3, r_values, r_values, z_values, linewidth = 8, color = (maxima_color, .4), overdraw=false, transparency =true, ssao = true)
lines!(ax3, r_values, r_values, z_values, linewidth = 8, color = (maxima_color, .6), overdraw=true, transparency=true, ssao = true)


# ===========================================================================
# ===========ONE CONVEX, ONE CONCAVE, DIFFERENT RLOW AND RHIGH===============
# ===========================================================================

function s1_curve(r)
    d=4
    shift = 0
     return .3 - max(0.5 * (1 - exp(-d * (r - shift - rlow)) - (r-shift) * 1/8), 0) + .05
end

function s2_curve(r)
    d = 1
    r = rmax-r  # Mirror the function
    shift = 7
    out = max((1 - 1.4^(-d * (r - shift - rlow))) * 0.7 - 1/16 * (r - shift) , 0) + .2
end


# Set the parameters
rlow1 = 2
rhigh1 = 6
rlow2 = 4
rhigh2 = 11.55 

s1(r) = add_flat_regions(s1_curve, r, rlow1, rhigh1)
s2(r) = add_flat_regions(s2_curve, r, rlow2, rhigh2)


# ------------ Survivorship Functions ---------------------------------------------------
top = .4  # how far the bands reach

ax1 = Axis(
    f[3,1],
    xlabel = "r",
    ylabel = "s₁(r) and s₂(r) ",
    aspect = 1,
    limits = (0, xmax, 0, top),
    xticks = ([rlow1, rhigh1, rlow2  + .45, rhigh2, xmax],   [L"r_{low}", L"r_{high}", L"r'_{low}", L"r'_{high}", L"r_{max}"]),  # Have to shift the rlow2 tick a bit. Don't know why.,  
    yticks = [0],
    xticklabelsize = ticksize,
    yticklabelsize = ticksize,

)
  
# Seperate the regions by bands
alpha = .3  # How faint the bands are
band!(
    [0, rlow1],
    [0, 0],
    [top , top],
    color = (color_cycle[1], alpha)
)
band!(
    [rlow1, rlow2 + .45],
    [0, 0],
    [top , top],
    color = ( color_cycle[2], alpha)
)
band!(
    [rlow2 + .45, rhigh1],
    [0, 0],
    [top, top],
    color = ( color_cycle[3], alpha)
)
band!(
    [rhigh1, rhigh2],
    [0, 0],
    [top, top],
    color = ( color_cycle[4], alpha)
)
band!(
    [rhigh2, xmax],
    [0, 0],
    [top, top],
    color = (color_cycle[5], alpha)
)

# Boundry Lines
# Change the alpha manually so that the lines remain preceptually constant between the regions
vlines!(rlow1, linestyle = gridsymbol, color = (gridcolor, 1), linewidth = gridwidthflat)
vlines!(rhigh1, linestyle = gridsymbol, color = (gridcolor, .6), linewidth = gridwidthflat)
vlines!(rlow2 + .45, linestyle = gridsymbol, color = (gridcolor, .9), linewidth = gridwidthflat)
vlines!(rhigh2, linestyle = gridsymbol, color = (gridcolor, .5), linewidth = gridwidthflat)

# Plot the survivorship functions (put the fainter line behind for a sense of depth)
lines!(
    0:.01:xmax, 
    s2,
    linewidth = lw,
    color = Cycled(2)  # An orange color in the Algebra of Graphics theme
    )
lines!(
    0:.001:xmax, 
    s1,
    linewidth = lw,
    color = survivorship_color
    )
    
# --------------------- The Grid of regions ---------------------------------------------

ax2 = Axis(
    f[3,2],
    xlabel = "r₁",
    ylabel = "r₂",
    aspect = 1,
    limits = (0, xmax, 0, xmax),
    xticks = ([rlow1, rhigh1, rlow2  + .45, rhigh2, xmax],  [L"r_{low}", L"r_{high}", L"r'_{low}", L"r'_{high}", L"r_{max}"]),  
    yticks = ([rlow1, rhigh1, rlow2  + .45, rhigh2, xmax], [L"r_{low}", L"r_{high}", L"r'_{low}", L"r'_{high}", L"r_{max}"]),  
    xticklabelsize = ticksize,
    yticklabelsize = ticksize,
     
)

# Region for surface 
# X-axis
alpha2 = .5
top = xmax
band!(
    [0, rlow1],
    [0, 0],
    [top , top],
    color = (color_cycle[1], alpha2),
    transparency = true
)
band!(
    [rlow1, rlow2 + .45],
    [0, 0],
    [top , top],
    color = ( color_cycle[2], alpha2),
    transparency = true
)
band!(
    [rlow2 + .45, rhigh1],
    [0, 0],
    [top, top],
    color = ( color_cycle[3], alpha2),
    transparency = true
)
band!(
    [rhigh1, rhigh2],
    [0, 0],
    [top, top],
    color = ( color_cycle[4], alpha2),
    transparency = true
)
band!(
    [rhigh2, xmax],
    [0, 0],
    [top, top],
    color = (color_cycle[5], alpha2),
    transparency = true
)

# Y axis
band!(
    [0, xmax],
    [rlow1, rlow1],
    [0 , 0],
    color = (color_cycle[1], alpha2),
    transparency = true
)
band!(
    [0, xmax],
    [rlow2+.45, rlow2+.45],
    [rlow1, rlow1],
    color = (color_cycle[2], alpha2),
    transparency = true
)
band!(
    [0, xmax],
    [rhigh1, rhigh1],
    [rlow2 + .45, rlow2 + .45],
    color = (color_cycle[3], alpha2),
    transparency = true
)
band!(
    [0, xmax],
    [rhigh2, rhigh2],
    [rhigh1, rhigh1],
    color = (color_cycle[4], alpha2),
    transparency = true
)
band!(
    [0, xmax],
    [xmax, xmax],
    [rhigh2, rhigh2],
    color = (color_cycle[5], alpha2),
    transparency = true
)

# Gridlines
vlines!(rlow1, linestyle = gridsymbol, color = gridcolor, linewidth = gridwidthflat, transparency = true)
vlines!(rhigh1, linestyle = gridsymbol, color = gridcolor, linewidth = gridwidthflat, transparency = true)
hlines!(rlow1, linestyle = gridsymbol, color = gridcolor, linewidth = gridwidthflat, transparency = true)
hlines!(rhigh1, linestyle = gridsymbol, color = gridcolor, linewidth = gridwidthflat, transparency = true)
vlines!(rlow2+.45, linestyle = gridsymbol, color = gridcolor, linewidth = gridwidthflat, transparency = true)
vlines!(rhigh2, linestyle = gridsymbol, color = gridcolor, linewidth = gridwidthflat, transparency = true)
hlines!(rlow2+.45, linestyle = gridsymbol, color = gridcolor, linewidth = gridwidthflat, transparency = true)
hlines!(rhigh2, linestyle = gridsymbol, color = gridcolor, linewidth = gridwidthflat, transparency = true)

# ---------- Fitness Landscape ---------------------------------------------------

R = 25
# I can't get Latex to work with the tick lables for the 3D plot. So we will add those in after with photoshop
ax3 = Axis3(
    f[3,3 ],
    azimuth =  -π/2  -.2,
    elevation = .9,
    xlabel = "r₁",
    ylabel = "r₂",
    zlabel = "W(r₁, r₂)",
    #aspect = (1,1,1),
    xticklabelsvisible = false,
    yticklabelsvisible = false,
    zticklabelsvisible = false,
    zticksvisible = false,
    xticks = [rlow1, rhigh1, rlow2  + .45, rhigh2, xmax],  
    yticks = [rlow1, rhigh1, rlow2  + .45, rhigh2, xmax], 
    xticklabelsize = ticksize,
    yticklabelsize = ticksize,
)

# Make the surface
regions = [0, rlow1, rlow2  + .45, rhigh1,  rhigh2, xmax]
for (i, r1) in enumerate(regions)
    for (j, r2) in enumerate(regions)
        space = wirespace
        if i < 6 && j < 6
            r1s = regions[i]:space:regions[i+1]  # x axis
            r2s = regions[j]:space:regions[j+1] # y axis
            zs = [W(1/2,  r1,  r2) for r1 in r1s, r2 in r2s]  # z axis

            # Plot the three of these
            
            wireframe!(
                    r1s, 
                    r2s, 
                    zs,
                    color = (color_cycle[i], .5),
                    linewidth = wirewidth,
                    transparency = true
                    )
            wireframe!(
            r1s, 
            r2s, 
            zs,
            color = (color_cycle[j], .5),
            linewidth = wirewidth,
            transparency = true
            )
            surface!(
                r1s, 
                r2s, 
                zs,
                colormap = (:greys, surfacealpha),
                )
        end
    end
end


# Plot the section boundries 
r1_values = 0:0.001:xmax
r2_values = repeat([rlow1], length(r1_values))
z_values = [W(1/2, rlow1, r)+gridoffset for r in r1_values]
lines!(ax3, r1_values, r2_values, z_values, linewidth = gridwidth, linestyle = gridsymbol, color = (gridcolor, gridalpha), overdraw=false, transparency = gridtransparecy)
lines!(ax3, r1_values, r2_values, z_values, linewidth = gridwidth, linestyle = gridsymbol, color = (gridcolor, .2), overdraw=true, transparency = gridoverdrawtransparency)

r1_values = 0:0.001:xmax
r2_values = repeat([rhigh1], length(r1_values))
z_values = [W(1/2, rhigh1, r)+gridoffset for r in r1_values]
lines!(ax3, r1_values, r2_values, z_values, linewidth = gridwidth, linestyle = gridsymbol, color = (gridcolor, gridalpha), overdraw=false, transparency = gridtransparecy)
lines!(ax3, r1_values, r2_values, z_values, linewidth = gridwidth, linestyle = gridsymbol, color = (gridcolor, .2), overdraw=true, transparency = gridoverdrawtransparency)

r2_values = 0:0.001:xmax
r1_values = repeat([rhigh1], length(r2_values))
z_values = [W(1/2, r, rhigh1)+gridoffset for r in r2_values]
lines!(ax3, r1_values, r2_values, z_values, linewidth = gridwidth, linestyle = gridsymbol, color = (gridcolor, gridalpha), overdraw=false, transparency = gridtransparecy)
lines!(ax3, r1_values, r2_values, z_values, linewidth = gridwidth, linestyle = gridsymbol, color = (gridcolor, .2), overdraw=true, transparency = gridoverdrawtransparency)

r2_values = 0:0.001:xmax
r1_values = repeat([rlow1], length(r2_values))
z_values = [W(1/2, r, rlow1)+gridoffset for r in r2_values]
lines!(ax3, r1_values, r2_values, z_values, linewidth = gridwidth, linestyle = gridsymbol, color = (gridcolor, gridalpha), overdraw=false, transparency = gridtransparecy)
lines!(ax3, r1_values, r2_values, z_values, linewidth = gridwidth, linestyle = gridsymbol, color = (gridcolor, .2), overdraw=true, transparency = gridoverdrawtransparency)

r1_values = 0:0.001:xmax
r2_values = repeat([rlow2 + .45], length(r1_values))
z_values = [W(1/2, rlow2 + .45, r)+gridoffset for r in r1_values]
lines!(ax3, r1_values, r2_values, z_values, linewidth = gridwidth, linestyle = gridsymbol, color = (gridcolor, gridalpha), overdraw=false, transparency = gridtransparecy)
lines!(ax3, r1_values, r2_values, z_values, linewidth = gridwidth, linestyle = gridsymbol, color = (gridcolor, .2), overdraw=true, transparency = gridoverdrawtransparency)

r1_values = 0:0.001:xmax
r2_values = repeat([rhigh2], length(r1_values))
z_values = [W(1/2, rhigh2, r)+gridoffset for r in r1_values]
lines!(ax3, r1_values, r2_values, z_values, linewidth = gridwidth, linestyle = gridsymbol, color = (gridcolor, gridalpha), overdraw=false, transparency = gridtransparecy)
lines!(ax3, r1_values, r2_values, z_values, linewidth = gridwidth, linestyle = gridsymbol, color = (gridcolor, .2), overdraw=true, transparency = gridoverdrawtransparency)

r2_values = 0:0.001:xmax
r1_values = repeat([rhigh2], length(r2_values))
z_values = [W(1/2, r, rhigh2)+gridoffset for r in r2_values]
lines!(ax3, r1_values, r2_values, z_values, linewidth = gridwidth, linestyle = gridsymbol, color = (gridcolor, gridalpha), overdraw=false, transparency = gridtransparecy)
lines!(ax3, r1_values, r2_values, z_values, linewidth = gridwidth, linestyle = gridsymbol, color = (gridcolor, .2), overdraw=true, transparency = gridoverdrawtransparency)

r2_values = 0:0.001:xmax
r1_values = repeat([rlow2+.45], length(r2_values))
z_values = [W(1/2, r, rlow2+.45)+gridoffset for r in r2_values]
lines!(ax3, r1_values, r2_values, z_values, linewidth = gridwidth, linestyle = gridsymbol, color = (gridcolor, gridalpha), overdraw=false, transparency = gridtransparecy)
lines!(ax3, r1_values, r2_values, z_values, linewidth = gridwidth, linestyle = gridsymbol, color = (gridcolor, .2), overdraw=true, transparency = gridoverdrawtransparency)


# Find the maxima 
max_val = -Inf
max_r1, max_r2 = -1, -1

for r1 in rlow:.1:rhigh
    for r2 in rlow:.05:rhigh
        val = W(1/2, r1, r2)
        if val > max_val
            max_val = val
            max_r1, max_r2 = r1, r2
        end
    end
end

max_val += .01

# Plot the maxima. There will be two because the landscape is symmetric
scatter!(ax3, [max_r1], [max_r2], [max_val], markersize = 16, color = (maxima_color, .8), overdraw = false, marker = maxima_marker)
scatter!(ax3, [max_r2], [max_r1], [max_val], markersize = 16, color = (maxima_color, .8), overdraw = false, marker = maxima_marker)
# Replot with transparency and overdraw for asthetics 
scatter!(ax3, [max_r1], [max_r2], [max_val], markersize = 20, color = (maxima_color, .4), overdraw = false, marker = maxima_marker, transparency = true, ssao = true)
scatter!(ax3, [max_r2], [max_r1], [max_val], markersize = 20, color = (maxima_color, .4), overdraw = false, marker = maxima_marker, transparency = true, ssao = true)
scatter!(ax3, [max_r1], [max_r2], [max_val], markersize = 20, color = (maxima_color, .4), overdraw = true, marker = maxima_marker, transparency = true, ssao = true)
scatter!(ax3, [max_r2], [max_r1], [max_val], markersize = 20, color = (maxima_color, .4), overdraw = true, marker = maxima_marker, transparency = true, ssao = true)


# generate points where r1 = r2 to show symmetric strategies
r_values = 0:0.001:xmax-.1
z_values = [W(1/2, r, r)+.004 for r in r_values]

# plot the line where r1=r2. 
# we need to plot multiple lines with varying values and options to get the transparency to work out correctly
lines!(ax3, r_values, r_values, z_values, linewidth = 3, color = (maxima_color, .2), overdraw=false, ssao = true)
lines!(ax3, r_values, r_values, z_values, linewidth = 3, color = (maxima_color, .2), overdraw=true, ssao = true)
lines!(ax3, r_values, r_values, z_values, linewidth = 8, color = (maxima_color, .4), overdraw=false, transparency =true, ssao = true)
lines!(ax3, r_values, r_values, z_values, linewidth = 8, color = (maxima_color, .6), overdraw=true, transparency=true, ssao = true)


f
````
